# Putative *cis*-regulatory elements predict iron deficiency responses in Arabidopsis roots

**DOI:** 10.1101/603290

**Authors:** Birte Schwarz, Christina B. Azodi, Shin-Han Shiu, Petra Bauer

**Affiliations:** Institute of Botany, Heinrich Heine University, Universitätsstr. 1, Düsseldorf, Germany; Department of Plant Biology, Michigan State University, East Lansing, MI, USA; Department of Computational, Mathematics, Science, and Engineering, Michigan State University, East Lansing, MI, USA; Cluster of Excellence on Plant Science (CEPLAS), Heinrich Heine University, Düsseldorf, Germany

**Keywords:** Arabidopsis, Fe deficiency, machine learning, Random Forest, co-expression clustering, *cis*-regulatory element, transcription factor binding motif, FIT, IDE1, coumarin

## Abstract

Iron (Fe) is a key cofactor in many cellular redox processes, including respiration and photosynthesis. Plant Fe deficiency (-Fe) activates a complex regulatory network which coordinates root Fe uptake and distribution to sink tissues, while avoiding over-accumulation of Fe and other metals to toxic levels. In Arabidopsis (*Arabidopsis thaliana*), FIT (FER-LIKE FE DEFICIENCY-INDUCED TRANSCRIPTION FACTOR), a bHLH transcription factor (TF), is required for up-regulation of root Fe acquisition genes. However, other root and shoot -Fe-induced genes involved in Fe allocation and signaling are FIT-independent. The *cis*-regulatory code, i.e. the *cis*-regulatory elements (CREs) and their combinations that regulate plant -Fe-responses, remains largely elusive. Using Arabidopsis genome and transcriptome data, we identified over 100 putative CREs (pCREs) that were predictive of -Fe-induced up-regulation of genes in root tissue. We used large-scale *in vitro* TF binding data, association with FIT-dependent or FIT-independent co-expression clusters, positional bias, and evolutionary conservation to assess pCRE properties and possible functions. In addition to bHLH and MYB TFs, also B3, NAC, bZIP, and TCP TFs might be important regulators for -Fe responses. Our approach uncovered IDE1 (Iron Deficiency-responsive Element 1), a -Fe response CRE in grass species, to be conserved in regulating genes for biosynthesis of Fe-chelating compounds also in Arabidopsis. Our findings provide a comprehensive source of *cis*-regulatory information for -Fe-responsive genes, that advances our mechanistic understanding and informs future efforts in engineering plants with more efficient Fe uptake or transport systems.

**One sentence summary:** >100 putative *cis*-regulatory elements robustly predict Arabidopsis root Fe deficiency-responses in computational models, and shed light on the mechanisms of transcriptional regulation.

## Introduction

The micronutrient iron (Fe) is crucial for survival of all organisms. Plants encounter Fe deficiency (-Fe) on calcareous and alkaline soils or during developmental phases with increased sink demands. As a central component of heme and Fe-sulfur (FeS) clusters, Fe acts in redox processes in plants in basically all important metabolic processes, such as the respiratory and photosynthetic electron transport chains, chlorophyll biosynthesis, DNA replication and repair, and nitrogen and sulfur assimilation. Consequently, plants react to -Fe with a range of molecular, physiological and morphological adjustments, which is reflected in transcriptional alterations of more than 1000 genes in Arabidopsis (*Arabidopsis thaliana*) (Dinneny et al., 2008; Rodríguez-Celma et al., 2013; Mai et al., 2016). In the shoots, the photosynthetic machinery is remodeled, leading to visible leaf chlorosis symptoms, and essential Fe-requiring processes are prioritized, which can be achieved through break-down of dispensable Fe-bound proteins and Fe redistribution between organelles (Blaby-Haas and Merchant, 2013; Balk and Schaedler, 2014; Hantzis et al., 2018). In the roots, genes controlling soil Fe uptake and detoxification of other transition metal ions acquired along with Fe are up-regulated. Additionally, Fe is mobilized from internal storages and distributed to Fe sinks. -Fe also leads to changes in root architecture and root hair morphology (Brumbarova et al., 2015; Curie and Mari, 2017; Jeong et al., 2017; Li and Lan, 2017).

To acquire soil Fe, grasses secrete mugineic acid (MA) family phytosiderophores and import Fe^3+^-MA complexes into the root (“Strategy II”). In contrast, non-grass monocots and dicots, such as Arabidopsis, acquire Fe via a reduction-based mechanism, in which soil Fe^3+^ is solubilized by lowering the local pH through proton extrusion, followed by reduction to plant-accessible Fe^2+^ at the root epidermis and Fe^2+^ uptake (“Strategy I”) (Marschner and Römheld, 1994). In Strategy I, secreted chelators (mainly phenylpropanoid-derived coumarins or riboflavin derivatives) aid efficient Fe^3+^ solubilization and reduction (Fourcroy et al., 2014; Schmid et al., 2014). Thus, Fe chelation is important during acquisition in both strategies.

Transcriptional control plays an important role in -Fe responses. A regulatory cascade ultimately controls a set of -Fe response genes. In both, Strategy I and II, the current cascade model involves related subgroups of basic helix-loop-helix (bHLH) transcription factors (TFs). When rice and Arabidopsis plants experience -Fe, subgroup IVc bHLH proteins activate subgroup Ib and IVb *BHLH* genes (Zhang et al., 2015; Liang et al., 2017). Downstream from IVc bHLH TFs (ILR3/bHLH34/bHLH104/bHLH115 in Arabidopsis, PRI1 in rice), subgroup Ib bHLH TFs (bHLH38/39/100/101 in Arabidopsis, IRO2 in rice) and subgroup IVb bHLH TFs (PYE in Arabidopsis, IRO3 in rice) regulate responses further downstream (Ogo et al., 2007; Yuan et al., 2008; Long et al., 2010; Zheng et al., 2010). In addition, IVc bHLH protein levels are controlled by Fe-regulated E3 ligases (Selote et al., 2015; Zhang et al., 2017).

Despite these conserved regulatory and functional interactions of subgroup IVc, Ib, and IVb bHLH TFs between grass and non-grass species, it remains unclear if other regulatory components between Strategy I and II are conserved. For example, in grasses, IDEF1 (IRON DEFICIENCY-RESPONSIVE ELEMENT BINDING FACTOR1, ABI3VP1 subfamily of B3 TF) and IDEF2 (NAC TF) coordinate -Fe responses through binding to IDE1 (Iron Deficiency-responsive Element 1) and IDE2 (Kobayashi et al., 2007; Ogo et al., 2008). IDE1 has been connected to induction of genes involved in Strategy II MA biosynthesis and Fe-MA uptake (Kobayashi et al., 2005; Ogo et al., 2007). However, while barley IDE1 can drive reporter gene expression in tobacco in a -Fe-dependent manner and IDE1-like motifs are present in several Arabidopsis -Fe response genes, a function for IDE1 has not been shown in Strategy I plants (Kobayashi et al., 2003; Kobayashi et al., 2005; Kobayashi et al., 2007; Murgia et al., 2011). Strategy I Fe acquisition requires the bHLH TF FIT (FER-LIKE IRON DEFICIENCY-INDUCED TRANSCRIPTION FACTOR) that is absent in rice (Colangelo and Guerinot, 2004; Jakoby et al., 2004), and is activated upon -Fe mainly through interaction with subgroup Ib TFs (Yuan et al., 2008; Sivitz et al., 2012; Wang et al., 2013). FIT is essential for up-regulation of Fe^3+^ reduction, Fe^2+^ uptake, and chelator biosynthesis and export (Colangelo and Guerinot, 2004; Jakoby et al., 2004; Sivitz et al., 2012; Schmid et al., 2014; Mai et al., 2016).

A co-expression network built with -Fe-responsive genes gives insight into the complex -Fe regulatory system in Arabidopsis (Ivanov et al., 2012). Among co-expression clusters, one contains root-specific and FIT-dependent genes involved in Fe acquisition, while another one is composed of root- and shoot-expressed FIT-independent genes. In this work, we refer to robust (i.e. consistently identified in different studies) FIT-dependent and FIT-independent genes as the “gold standard” (GS) -Fe-induced genes. The concept to discriminate FIT-dependent and FIT-independent co-expression clusters has proven very informative for interpreting mutant phenotypes and to place novel regulators into the -Fe response cascade (e.g. Zhang et al., 2015; Liang et al., 2017; Gratz et al., 2019). FIT-independent network genes mostly act in sub-cellular and long-distance transport and distribution of Fe and in Fe signaling and they include subgroup Ib *BHLH* genes and *PYE* (Ivanov et al., 2012). Only few upstream regulators for FIT-independent gene expression have been identified yet, namely bHLH IVc TFs, controlling Ib *BHLH* and *PYE*, and PYE controlling *NAS4*, *ZIF1* and *FRO3* of the same co-expression regulon (Long et al., 2010).

For most -Fe-responsive genes, including reliable marker genes (Ivanov et al., 2012; Mai et al., 2016), the specific *cis*-regulatory elements (CREs) which coordinate their expression are unknown. Computational approaches uncover regulatory connections on a genome-wide scale, such as through elucidating the *cis*-regulatory code, i.e. the collection of CREs and the genes they regulate in a given regulatory context (Yáñez-Cuna et al., 2013). Putative CREs (pCREs) could be identified computationally by the over-representation of sequences in the promoter regions of co-regulated genes. Combining with data for TF binding motifs (TFBMs) in Arabidopsis (Weirauch et al., 2014; O’Malley et al., 2016), regulatory connections can be made between TFs, binding sequences, and target genes. To further improve the confidence of computationally derived *cis*-regulatory code, machine learning algorithms (reviewed in Ma et al., 2014) can be applied to build models with pCREs to predict gene expression or transcriptional responses. These models can be evaluated by making predictions on expression of genes that are not part of the model training. Most importantly, a good model indicates that the pCREs used are most likely important for regulating the expression/response of interest. Previous work has demonstrated the suitability of machine learning for elucidating the *cis*-regulatory code of environmental stress responses in Arabidopsis (Zou et al., 2011; Uygun et al., 2017).

To get a deeper understanding of -Fe response regulation, we elucidate the underlying *cis*-regulatory code. Because some TFs have well established roles in -Fe response, we can use these to validate our findings. We combined genome, transcriptome, and *in vitro* protein-DNA interaction data to uncover links between pCREs controlling -Fe responses and their upstream TFs. With pCREs over-represented in promoters of co-expressed genes we modeled -Fe-induced up-regulation and identified over 100 informative pCREs of -Fe-responsive processes.

## Results and Discussion

### Overview of approach and functions of -Fe-responsive genes

To identify root -Fe-associated CREs at a genome-wide scale, we defined root -Fe response co-expression clusters, then we identified *k*-mers enriched in the promoter regions of those genes, and finally we modeled -Fe response on the basis of the enriched promoter *k*-mers. An overview of our complete workflow including functional analysis of the identified pCREs is shown in **Figure 1A**. Because many factors, such as the choice of data set or the measure used to define expression similarity, impact the discovery of functional connections between genes (Uygun et al., 2016), we used multiple expression data combinations and algorithms with varying parameters (see **Methods**).

**Figure 1.**
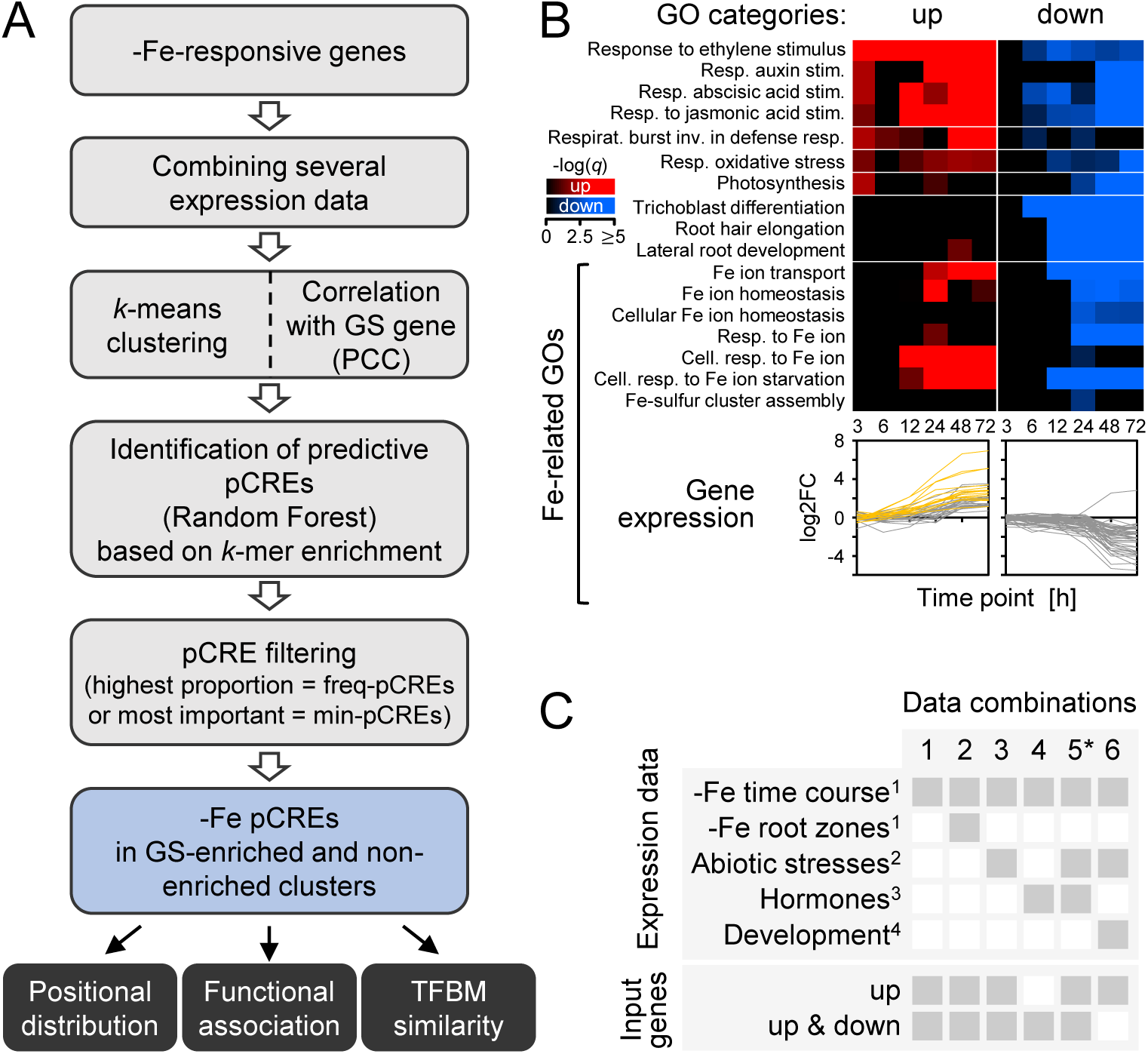
-Fe pCRE identification workflow and transcriptomic data. **A:** pCRE identification workflow. **B:** Heatmap of enrichment (FET, *q*<0.05) of selected GO terms in genes that were significantly up- (red) or down-regulated (blue) (*q*<0.05) at ≥1 of 6 time points in -Fe-treated roots of 6 d-old seedlings (Dinneny et al., 2008). Differential regulation was defined as log_2_ fold-change (log2FC) >1 or <-1 (treatment vs. control). GOs are sorted by category, and expression patterns of genes corresponding to Fe-related GOs are shown below the heatmap. Yellow genes indicate -Fe GS genes. **C:** Transcriptomic data combinations which were used for clustering of co-expressed genes. Gray filled boxes in columns depict (**top**) expression data used in the combination and (**bottom**) if up (up-regulated only) or up & down (up- and down-regulated) genes were included. ^1^(Dinneny et al., 2008), ^2^(Kilian et al., 2007), ^3^(Goda et al., 2008), ^4^(Schmid et al., 2005), *Tested with (5a) and without (5b) genotoxic stress data, (5b) input only up-regulated genes.

-Fe-responsive genes (log_2_ fold-change (log2FC) >1 or <-1, *q*<0.05) were identified using transcriptomic data available for six time points after an -Fe treatment in Arabidopsis seedling roots (Dinneny et al., 2008). Enrichment analysis (Fisher’s exact test (FET), *q*<0.05) of biological process gene ontologies (GOs) showed that, next to Fe-related GOs (e.g. Fe transport, homeostasis and FeS cluster assembly), responses to several hormones, including auxin, ethylene, abscisic acid and jasmonic acid, were over-represented (**Figure 1B**). This is consistent with the roles of hormones in -Fe response (Brumbarova et al., 2015) and in root and root hair morphology and development (Schmidt et al., 2000), which were also enriched GOs. -Fe affects the photosynthetic machinery and often correlates with oxidative stress responses (Rodríguez-Celma et al., 2013), which is reflected in enrichment of GOs regarding oxidative stress, photosynthesis, and primary metabolism even in roots (**Figure 1B**; **Supplemental Figure S1**).

### Using multiple expression data sets to define -Fe co-expression clusters

We next grouped differentially regulated -Fe response genes into co-expression clusters using two approaches: *k*-means clustering and correlation to gold standard. *K*-means clustering was based on the transcriptional responses to -Fe alone and combined with different responses to other stress and developmental conditions (**Figure 1C**) (Schmid et al., 2005; Kilian et al., 2007; Dinneny et al., 2008; Goda et al., 2008). Correlation-based clusters were generated for each gene in our curated list of gold standard (GS) -Fe response genes (see **Methods**; **Supplemental Table S1**), by selecting the differentially regulated -Fe response genes with a significantly similar (Pearson’s Correlation Coefficient (PCC); see **Methods**) expression pattern to the GS gene, also using the different combinations of transcriptional data (**Figure 1C**). To identify co-expression clusters with similar biological functions, we grouped them according to their enriched GOs (FET, *q*<0.05) into “superclusters” (**Figure 2A**; **Supplemental Figure S2A**; see **Methods**), which were defined as groups of at least 20 clusters with significantly higher similarity to each other than the average similarity of all clusters (all Mann-Whitney U, *p<*2.2e-16; **Supplemental Figure S2B, C**). While *k*-means supercluster C was enriched in an Fe-related GO (cellular response to Fe, GO shared by ≥75% of co-expression clusters within each supercluster), *k*-means superclusters A and B shared GOs related to different stress responses.

**Figure 2.**
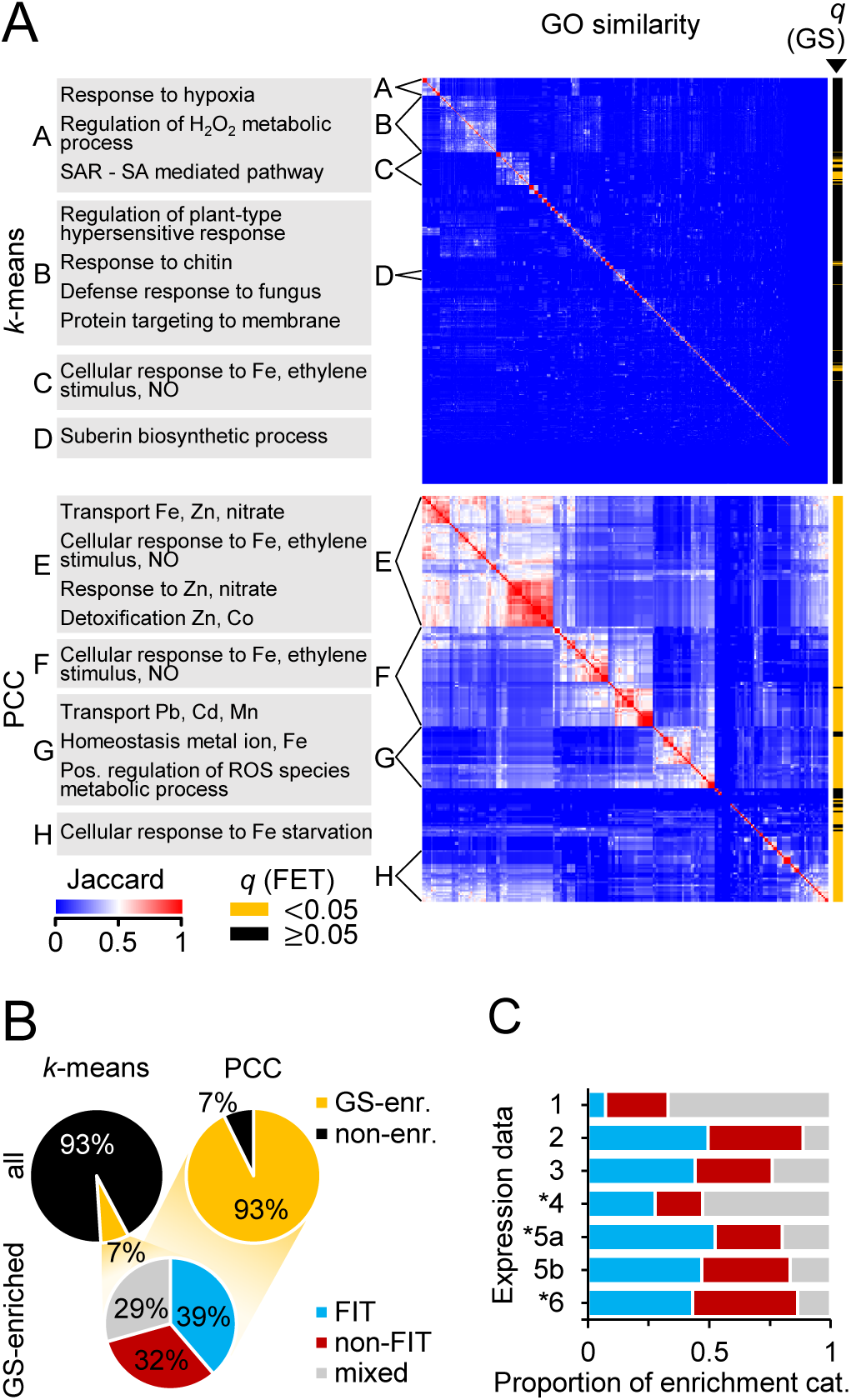
Characterization of the defined co-expression clusters by GO terms, -Fe GS gene content and FIT-dependent/FIT-independent gene content. **A:** Heatmap of GO similarity between co-expression clusters from *k*-means clustering (**top**, n=985 clusters) and GS gene correlation (PCC; **bottom**, n=238), containing up-regulated genes (Figure 1C; up- and down-regulated genes: **Supplemental Figure S2A**). Clusters were grouped by hierarchical clustering and superclusters (A-F) were defined as groups of >20 clusters that have a within-mean Jaccard Index significantly higher than the mean Jaccard Index of all clusters. Enriched GO terms shared by ≥75% (*k*-means) and ≥90% (PCC) of the clusters in each supercluster are shown (**left**). Co-expression clusters enriched for -Fe GS genes are designated (yellow, **right**). **B:** Proportions of all *k*-means (**top left**) and PCC (**top right**) co-expression clusters in which -Fe GS genes are significantly over-represented (yellow). Of those (**bottom**), the proportion enriched for FIT-dependent genes (FIT, blue), FIT-independent genes (non-FIT, red) or for both (mixed, gray) was calculated. **C:** Proportion of FIT, non-FIT, and mixed clusters found using each expression data combination (as in Figure 1C). 1: -Fe time course, 2: time course + root zones, 3: time course + abiotic stresses, 4: time course + hormone treatments, 5a: time course + abiotic stresses + hormones, 5b: as 5a, genotoxic stress deleted, 6: time course + abiotic stresses + developmental data. *PCC clusters only. All enrichment analyses: FET, *q*<0.05.

Because the current TAIR GO annotation for -Fe response-related processes does not contain all -Fe-responsive genes of interest (e.g. *MYB10*, *UGT72E1*, *AT3G07720*, *FEP3* or *NAS4*, (Ivanov et al., 2012)), we also determined if *k*-means-generated co-expression clusters were enriched for GS genes (“GS-enriched”; FET, *q*<0.05; **right**, **Figure 2A**). While many of these clusters were part of the Fe-related GO supercluster A, the GS approach allowed us to identify an additional 23 Fe-related co-expression clusters that would have been overlooked by conventional GO enrichment analysis. In total, 7% of the *k*-means-generated clusters were GS-enriched (**Figure 2B**). Applying this same analysis to the correlation-based clusters (**Figure 2A**; **Supplemental Figure S2A**), we found higher levels of similarity between correlation-based clusters compared to *k*-means clusters (Mann-Whitney U, *p*<2.2e-16; **Supplemental Figure S2D**), because we pre-condition their identification on GS genes, some of which are tightly co-regulated (Ivanov et al., 2012). Accordingly, we found that 93% were GS-enriched (**right**, **Figure 2B**).

GS genes are either FIT target (“FIT-dependent”) or FIT-independent Fe homeostasis (“FIT-independent”) genes, which we found reflected in our GS-enriched clusters: 71% of the GS-enriched clusters were more specifically enriched for FIT-dependent (39%) and/or FIT-independent genes (32%) (FET, *q*<0.05; **bottom**, **Figure 2B**). The remaining GS-enriched clusters were enriched for both (“mixed”). Interestingly, clusters based on combined expression data (i.e. data combinations (dc) 2, 3, 5a/b, 6) were more often enriched for FIT-dependent or FIT-independent genes, while -Fe time course data alone (dc1) produced mainly mixed category clusters (**Figure 2C**). The utility of including spatial or developmental data (dc2, 6) to define co-expression clusters reflects that -Fe-responsive genes act at different time points and in different tissues and organs (Dinneny et al., 2008; Ivanov et al., 2012; Jeong et al., 2017). Finally, genes in clusters not enriched for GS-genes (“non-enriched”) tended to respond to particular abiotic stresses, for example cold (**Supplemental Figure S8**; e.g. clusters 937, 973) or salt (e.g. clusters 900, 915, 936, 987), whereas gene expression for GS-enriched cluster genes tended to randomly oscillate under different abiotic stresses (e.g. clusters 818, 835, 858, 889), which might indicate different regulatory networks and highlight the usefulness of incorporating additional abiotic stress data (as in dc3, 5, 6) when defining co-expression clusters that are likely co-regulated.

In summary, by using different expression data sets and clustering methods we defined 1,959 -Fe co-expression clusters, many of which were enriched for FIT-dependent and/or FIT-independent GS genes. These represent possibly co-regulated functional units in Fe acquisition and Fe homeostasis processes, well-suited to identify pCREs which can explain -Fe-induced up-regulation. Genes in co-expression clusters that were enriched in -Fe-responsive genes but not GS genes (non-enriched clusters) are presumably regulated by mechanisms different from GS-enriched clusters.

### A machine learning approach to model regulation of -Fe responsive co-expression clusters

The machine learning algorithm Random Forest (RF) has been successfully used to model stress transcriptional response using *cis*-regulatory sequences in plants (Zou et al., 2011; Deng et al., 2017; Uygun et al., 2017). Here, for each co-expression cluster, we used pCREs (enriched *k*-mers in putative promoter sequences; see **Methods**) to build a RF model that classifies genes as belonging to the cluster in question or as a non-responsive gene (see **Methods**). The pCREs from models performing above a defined threshold (F1≥0.7; see **Methods**) were then considered further. Out of 1,959 co-expression clusters, 28% of the models passed the performance threshold, 60% performed poorly, and for 12% no model could be built due to small size (median size=2 genes; **Supplemental Figure S3A, B**). Poor performing models (median F1=0.62) were mostly for small clusters (median size=12) (**Supplemental Figure S3B**; **Supplemental Table S2**) likely due to the lack of training data. Nonetheless, 66 large clusters (>100 genes, median size=135) also performed poorly (median F1=0.64) – this is likely because these large clusters are too heterogeneous containing genes with multiple regulatory codes (Uygun et al., 2016; Uygun et al., 2017), and/or are co-regulated but not at the transcriptional level (e.g. post-translationally controlled). Interestingly, of the 28% of clusters with models above the threshold, only 36% were GS-enriched clusters (**Supplemental Figure S3A**). Nonetheless, models built for GS-enriched clusters (median F1=0.68) tended to perform better than models built for non-enriched clusters (median F1=0.65; Mann-Whitney U, *p*<2.358e-09; **Figure 3A-C**).

**Figure 3.**
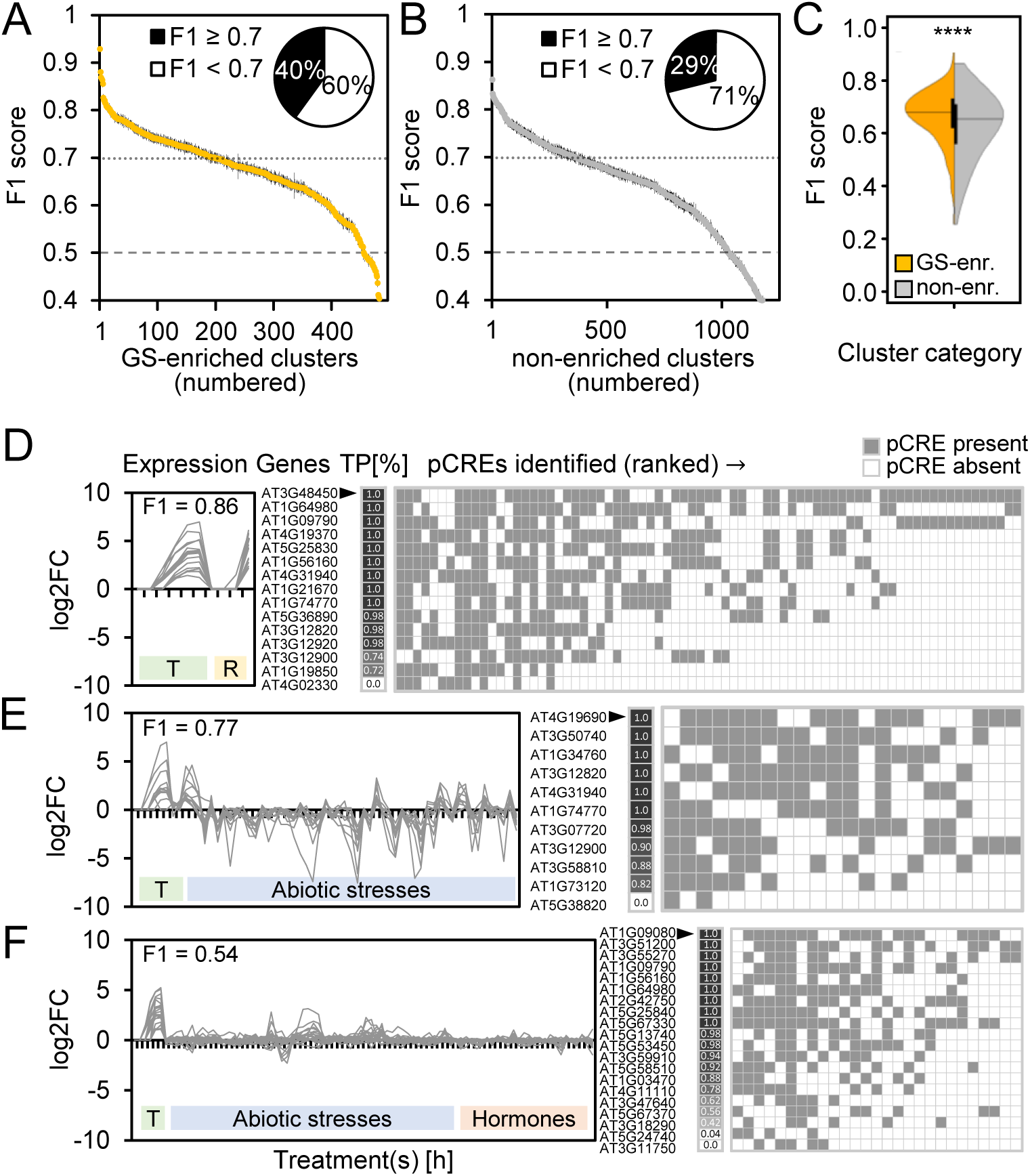
Performance of -Fe response RF prediction models. **A:** F1 scores of all GS-enriched clusters (n=495). **Inset**: Proportions of well-performing (F1≥0.7) and poorly performing clusters among the GS-enriched clusters. **B:** F1 scores of all non-enriched clusters (n=1,240). **Inset**: Proportions of well-performing and poorly performing clusters among the non-enriched clusters. **C:** Mean F1 score distributions of all GS-enriched clusters (yellow) and non-enriched clusters (gray). Statistical analysis: Mann-Whitney U (**** *p*<2.358e-09). **D-F**: Example GS-enriched co-expression clusters with good (**D**, **E**; cluster IDs: 493, 823) and bad (**F**; cluster ID: 1297) model performance. (**Left**) Expression (log_2_ fold-change: log2FC) profile of all genes in the co-expression cluster. (**Center**) Percent of times across RF replicates each gene was correctly predicted as -Fe-responsive (true positive (TP); black=100%, white=0%). (**Right**) pCREs sorted by importance rank (top ranked pCRE on the left) with heatmap designating when pCRE was present (gray) or absent (white) in a gene’s promoter. T: -Fe treatment time course. R: -Fe-treated root zones 1-4. F1 score: harmonic mean of precision and recall, with 1=perfect prediction and 0.5=random guessing. Cluster IDs and details: **Supplemental Table S2**.

Good model performance indicates that genes in a cluster are more likely co-regulated, and, because pCREs were used to build the model, these pCREs are likely the regulatory sequences contributing to the co-regulation. Taken together, we identified 5,639 pCREs enriched in promoters of -Fe-responsive genes that may be predictive of -Fe-induced up- or down-regulation. To further evaluate the biological relevance of pCREs, in the following sections, we assess pCREs based on their association with GS-enriched or non-enriched clusters, importance for model performance, and similarity to known TF binding sites. Known -Fe CREs from Arabidopsis and also from grasses, for example E-/G-boxes (bHLH TF binding sites) and IDE1, will serve as positive controls.

### Identifying common pCREs across co-expression clusters

We expect true -Fe response CREs to be: (i) important for building models with good performance in predicting -Fe response, and (ii) reliably identified in co-expression clusters with similar gene content. Therefore, for each pCRE we calculated the proportion of clusters enriched for the pCRE and its average importance rank across those clusters (**Supplemental Table S3**). The importance rank of a pCRE was derived from an importance score for the pCRE in question from the trained RF models that reflects how useful a pCRE was for predicting -Fe response genes in a cluster. This allowed us to get a snapshot of pattern of presence and absence of important pCREs for genes correctly predicted (true positives (TP)) in co-expression clusters with good (**Figure 3D, E**) and poor (**Figure 3F**) performance. The pCREs were enriched in between 1 (0.6%) and 56 (35%) GS-enriched clusters and in between 1 (0.3%) and 54 (15%) non-enriched clusters, with 173 pCREs considered frequent pCREs (freq-pCREs, enriched in >5% of GS-enriched or non-enriched clusters) (**Supplemental Table S3**). Across GS-enriched clusters, pCREs tended to have higher proportions with higher importance ranks than across clusters that were not GS-enriched (Mann-Whitney U, *p*<2.2e-16; **Figure 4A, B**; **Supplemental Figure S4A, B**). The higher proportion of GS-enriched cluster pCREs can be explained partly by the fact that those clusters are more homogenous in terms of gene contents than non-enriched clusters (Mann-Whitney U, *p*<2.2e-16; **Supplemental Figure S3C**).

**Figure 4.**
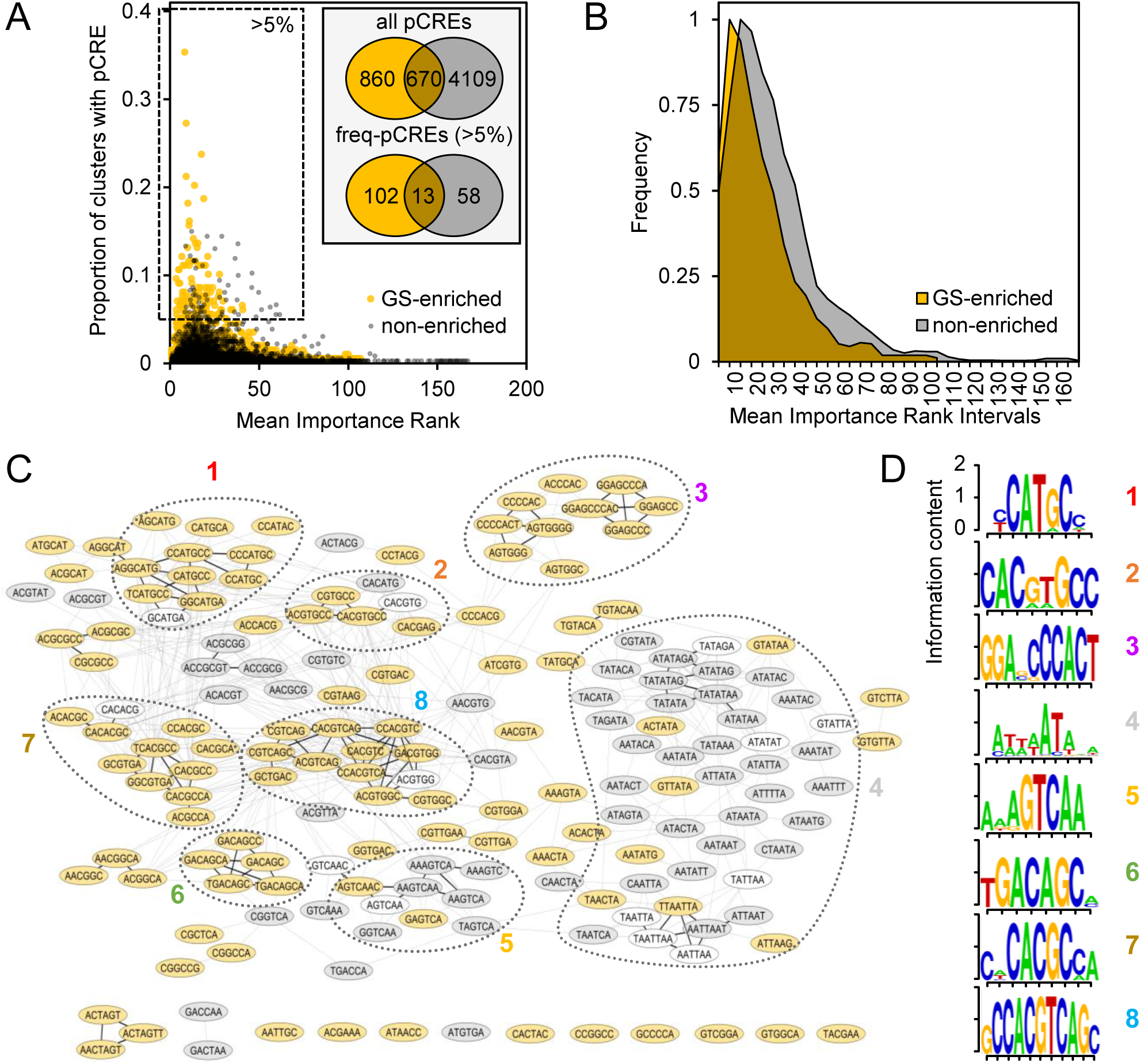
Analysis of pCREs predictive of -Fe co-expression clusters. **A:** Proportion of GS-enriched (yellow) and non-enriched (gray) clusters in which each pCRE (total n=5,639) was identified (y-axis) and mean importance rank (1=most important) of that pCRE in those clusters (x-axis). **Inset**: Numbers of unique and shared pCREs of GS-enriched and non-enriched cluster categories. **Upper**: all 5,639 pCREs. **Lower**: pCREs identified in >5% of GS-enriched or non-enriched clusters (n=173; freq-pCREs). **B:** Frequency of normalized mean importance ranks across all pCREs in GS-enriched (yellow) and non-enriched (gray) clusters. **C:** Cytoscape network of the 173 freq-pCREs based on sequence similarity, where similar pCREs (nodes) are connected by edges representing pair-wise correlation (PCC) distance of freq-pCRE PWMs. Bold black edges: distance=0. Light gray edges: distance ≤0.22. Highly interconnected freq-pCREs were arranged in groups and numbered. Hierarchical clustering representation of PCC distances: **Supplemental Figure S5E**. Yellow filled: freq-pCRE unique for GS-enriched clusters, gray filled: freq-pCRE unique for non-enriched clusters, not filled: shared freq-pCRE. **D:** PWMs of merged freq-pCREs from the same group (as in 4C).

We next determined whether GS-enriched clusters are regulated by a different set of pCREs than non-enriched clusters. Out of the 5,639 pCREs, 15% (n=860) were unique to GS-enriched clusters, while 73% (n=4109) were unique to non-enriched clusters and 12% (n=670) were found in both GS-enriched and non-enriched clusters (**inset**, **Figure 4A**). This indicates that GS-enriched clusters and non-enriched clusters are regulated partly by different pCREs, but also by a fraction of shared pCREs. However, 43% (n=286) of the 670 shared pCREs were predominant to GS-enriched clusters (i.e. having only low proportion and low importance rank in non-enriched clusters; **top**, **Supplemental Figure S4C**). This indicates that pCREs that were categorized as shared might not be equally important for regulating both GS-enriched and non-enriched clusters. Interestingly, unique GS-enriched freq-pCREs represented 59% (n=102) of the 173 freq-pCREs, while 35.5% (n=58) were unique non-enriched, and only 7.5% (n=13) freq-pCREs were shared between GS-enriched and non-enriched clusters, indicating that pCREs with high proportion are also the ones which seem to regulate almost exclusively either GS-enriched or non-enriched cluster functions, but not both (**inset**; **Figure 4A**; **bottom**, **Supplemental Figure S4C**). Furthermore, freq-pCREs tended to have higher importance ranks than non-frequent pCREs (Mann-Whitney U, *p*<1.924e-14; **Supplemental Figure S5D**). Together, this suggests that freq-pCREs could be particularly relevant for regulation of -Fe response mechanisms.

To characterize the freq-pCREs, we grouped them according to sequence similarity using pair-wise PCC distances of pCRE position weight matrices (PWM; see **Methods**). 62% (n=107) of all freq-pCREs could be placed into one of eight pCRE groups (**Figure 4C, D**; **Supplemental Figure S4E**). Freq-pCREs of the same group tended to be predictors of the same cluster category (GS-enriched/non-enriched).

In summary, we identified more than 100 -Fe pCREs that were reliably associated either exclusively to GS-enriched or non-enriched co-expression clusters or with high preference for one of the categories. Those pCREs were also ranked as important for machine learning models and might therefore be candidates for functionally relevant motifs to different responses to -Fe.

### Similarity of -Fe pCREs to known TFBMs

CREs are recognized by TFs to modulate gene expression. To identify what types of TFs may bind to the identified pCREs, we examined the similarities between the -Fe pCREs and known TF binding motifs (TFBMs) from two sources (see **Methods**). Based on threshold similarities, we were able to match a specific TF and/or a specific TF family to each of the 173 freq-pCREs (see **Methods**; **Figure 5A**; **Supplemental Figure S5**). To gain an overview which TF families might be associated with GS-enriched clusters and how specific these TF families are, we asked which families contained over-represented numbers of TFs that likely bound pCREs from GS-enriched and non-enriched cluster categories. We found that most TF families were found with higher proportion in either GS-enriched clusters (14 TF families) or non-enriched clusters (12), while only four were similarly distributed between both categories (**Figure 5B**).

**Figure 5.**
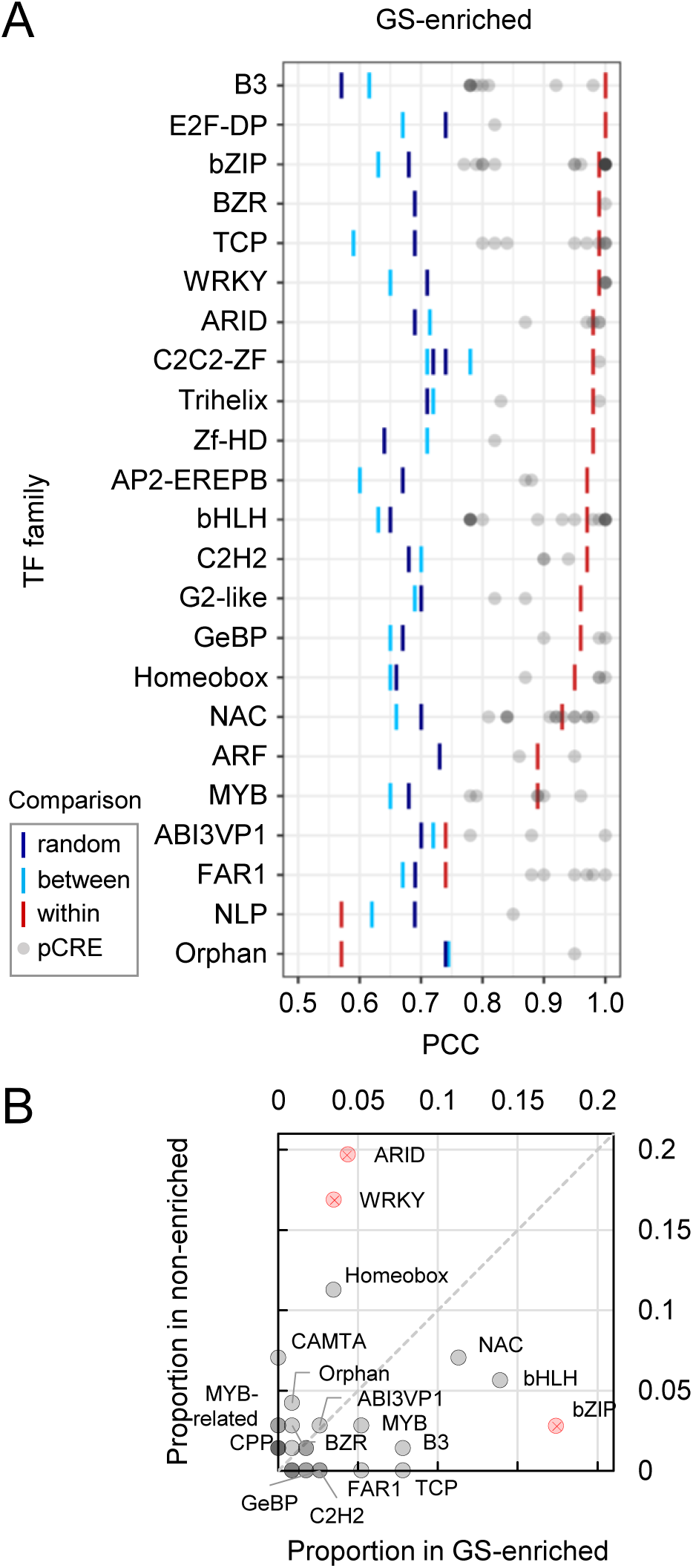
Similarity of freq-pCREs to *in vitro* TFBMs. **A:** Significance of sequence similarity for freq-pCREs from GS-enriched clusters and the best matching known TFBM. Bars represent 95^th^ percentile (PCC) significance thresholds for within TF family (red, pCRE sequence is more similar to a specific TFBM than other TFBMs from the same family), between TF families (light blue, pCRE sequence is more similar to a TFBM in a TF family than TFBMs from other TF families), or random (dark blue, pCRE sequence is more similar to a TFBM from a family than random 6-mers). Similarity of freq-pCREs from non-enriched clusters to TFBMs: **Supplemental Figure S5**. **B:** Proportion of TF family TFBMs (representing freq-pCRE matches meeting at least “between” threshold) in GS-enriched clusters (x-axis) and non-enriched clusters (y-axis). TFBM matches significantly over-represented (FET, *q*<0.05) in the GS-enriched or non-enriched cluster category are depicted in red and marked with “X”. Dashed line marks theoretical position for TF family TFBMs with the same proportion in both categories.

Most known -Fe regulators in Arabidopsis are bHLH TFs (FIT, subgroup Ib and IVc bHLH proteins, PYE, e.g. (Jakoby et al., 2004; Wang et al., 2007; Long et al., 2010; Palmer et al., 2013; Zhang et al., 2015)). bHLH and MYB TF families were identified, and even with higher proportion in GS-enriched clusters than in non-enriched clusters, which is indeed consistent with their role in -Fe response regulation. Other matching TF families over-represented in GS-enriched clusters were bZIP (FET, *q*<0.05), B3, TCP and NAC. Although a B3 TF (ABI3VP1 subfamily; IDEF1) and a NAC TF (IDEF2) are important regulators of Strategy II Fe acquisition in grasses (Kobayashi et al., 2007; Ogo et al., 2008), the role for these TF families in Strategy I non-grass plant species has not yet been described. In contrast, ARID, WRKY (both FET, *q*<0.05), Homeobox, and CAMTA TF families were matched more in non-enriched than GS-enriched clusters, pointing towards roles during -Fe stress other than Fe uptake or homeostasis, in which GS genes are mostly involved.

Next, freq-pCRE-TFBM matches from GS-enriched clusters served to infer specific upstream regulators of -Fe-responsive modules. More than 50% freq-pCREs (60 out of 115) matched TFBMs of a specific TF (**Figure 5A**). Of those, 29 freq-pCREs shared perfect sequence similarity (PCC=1) to the TFBM, which were then of particular interest. From these perfect matches, 23 pCREs were unique for GS-enriched clusters. Example TF candidates for these 23 cases were FUS3 (an ABI3VP1/B3 TF), bHLH104, bZIP3, 16 and 42, TCP13 (PTF1), and FAR1 (**Supplemental Table S4A**). While the DAP-seq and CIS-BP TFBM databases contain binding information for many TFs, they are far from exhaustive. For example, out of 162 known Arabidopsis bHLHs (Bailey et al., 2003), only 46 were available to be included in the analysis. Therefore, some TF families were likely under-represented in our analysis and some top match TFBMs may not accurately reflect the binding partner for certain pCREs. Consequently, some important pCRE-TFBM matches might not be detectable at this time. However, as new experimental TF binding data is collected, we might gain more biological insight into our -Fe pCREs.

### Inferring upstream regulators of the -Fe response

Because we believe our genome-wide approach for identifying regulatory elements may shed light on areas of -Fe response that are less well understood, we next put our findings in context with open questions in the field. For example, the ABI3VP1/B3-type TF IDEF1, a key regulatory factor of -Fe responses in rice and barley roots, recognizes the CATGC core of IDE1 (Kobayashi et al., 2003; Kobayashi et al., 2005; Kobayashi et al., 2007; Kobayashi et al., 2009; Kobayashi et al., 2010). With ten of our freq-pCREs having an IDE1 CATGC (or GCATG) core and matching ABI3VP1/B3 family TFBMs, IDE1-likes were fairly dominant among the freq-pCREs and unique to GS-enriched clusters (**Supplemental Table S3**). This strongly suggests an important function for IDE1-like motifs in Arabidopsis. Arabidopsis AFLs (B3 family TFs ABI3/FUS3/LEC2), are the closest homologs of the rice IDEF1, and may bind to the IDE1-likes. In fact, ABI3 and FUS3 bind to RY-like elements (CATGCA), regulating FeS cluster subunit formation during seed maturation (Roschzttardtz et al., 2009). However, ABI3 or FUS3 functions during later developmental stages, particularly in the root during -Fe response, remain to be elucidated. Since the FUS3 TFBM matched our top most abundant IDE1-like (CATGCC; **Supplemental Table S4A**), and because *FUS3* is expressed in the root epidermis and in lateral root primordia during later developmental stages (Boulard et al., 2017; Tang et al., 2017), FUS3 might be an IDEF1 homolog in Strategy I plants.

Another -Fe response-related TF in rice and barley, IDEF2, belongs to the NAC family and binds to the CA(A/C)G(T/C)(T/C/A)(T/C/A) core in IDE2 (Ogo et al., 2008). Although we did not have a perfect (PCC=1) pCRE-NAC TFBM match, we found matched NAC TFBMs slightly over-represented in GS-enriched clusters (**Figure 5B**). Furthermore, two of the top ten most abundant freq-pCREs unique to GS-enriched clusters matched NAC TFBMs (PCC>0.9), with one freq-pCRE being highly similar to the IDE2 core (CACGCC). This indicates that IDE2-like motifs might also play a role during Arabidopsis -Fe responses.

One freq-pCRE (CGTGCC) perfectly matched to a bHLH104 TFBM (**Supplemental Table S4A**). bHLH104 binds to the promoters of subgroup Ib *BHLH* genes *BHLH38*/*39*/*100*/*101* (Zhang et al., 2015; Li et al., 2016), positively regulating Fe uptake. Consistently, we found CGTGCC in clusters containing *BHLH101* (*AT5G04150*; **Supplemental Table S2**).

Other freq-pCREs matched to known TFBMs from TFs with unknown roles in -Fe response. For example, bZIP TFBMs were significantly over-represented in GS-enriched clusters, but have no known direct roles in -Fe response. However, bZIP TFs are known regulators of the Zn deficiency response, which, together with the fact that one GS gene, *ZIP9*, is also responsive to Zn deficiency, could indicate an interdependency of Zn and Fe homeostasis (Assunção et al., 2010; Sinclair et al., 2018). Furthermore, two matched TFs, bZIP3 and bZIP16, are involved in ABA signaling, which is connected to -Fe response amongst others by modulating root growth (Séguéla et al., 2008; Matiolli et al., 2011; Hsieh et al., 2012). Possible functions of bZIP TFs in response to -Fe stress should be explored in the future.

TCP13 (PTF1) and FAR1 TFBMs are two more examples for perfect freq-pCRE matches with yet unknown specific roles of the TFs during -Fe, although their specificity to GS-enriched clusters points towards important roles in regulating GS genes. TCPs are involved in plant development, but also act in signaling of hormones that influence -Fe responses (Davière et al., 2014; Brumbarova et al., 2015; Resentini et al., 2015; Nicolas and Cubas, 2016). For example, TCP20 was reported to bind to the *BHLH39* promoter (Andriankaja et al., 2014), indicating a possible connection of TCPs and -Fe responses during plant development. TCP13 is involved in regulating responses to light shade signals through *PHYTOCHROME INTERACTING FACTORS* (*PIF*s) (Zhou et al., 2018). Interestingly, FAR1 and its homolog FHY3 also act in phytochrome-PIF signaling (Wang and Wang, 2015). Together, this suggests a connection of light perception and -Fe responses mediated through these TFs, which is consistent with the known diurnal influence on Fe uptake (Vert et al., 2003; Santi and Schmidt, 2009; Hong et al., 2013; Salomé et al., 2013). In addition, FAR1/FHY3 act in the regulation of phosphate starvation response, together with ethylene regulator EIN3 (Liu et al., 2017), which also binds FIT to promote Fe uptake (Lingam et al., 2011). Therefore, FAR1 might also regulate Fe acquisition via the ethylene pathway.

Finally, a perfect freq-pCRE-WRKY11 match indicates that WRKY TFs, although significantly over-represented in non-enriched clusters, are also important for regulating GS-enriched clusters. WRKY11 is involved in abiotic stress tolerance in Arabidopsis (Ali et al., 2018), with no specific role known during -Fe response yet. However, WRKYs in general have already been connected to -Fe, for example as putative regulators of the coumarin transporter gene *PDR9* (Ito and Gray, 2006) and of *PYE* (Koryachko et al., 2015). Furthermore, WRKY46 negatively regulates the vacuolar Fe importer gene *VTL1*/*VITL1* (Gollhofer et al., 2011; Gollhofer et al., 2014; Yan et al., 2016). We found a WRKY TFBM (GTCAAC) in several non-enriched clusters containing down-regulated Fe-responsive genes, including the *VTL1* homolog *VTL5* (*AT3G25190*; **Supplemental Table S2**), indicating that some of the TFs matching non-enriched cluster pCREs might act as repressors of Fe excess genes.

In summary, many pCREs commonly found among GS-enriched clusters shared significant sequence similarity with known -Fe CREs, such as IDE1, or with binding sites of known -Fe-associated TF families, such as ABI3VP1/B3, NAC, MYB and bHLH. Notably, we found evidence for IDE1-like motifs being relevant not only in Strategy II plants, but also in the Strategy I plant Arabidopsis. Our results also suggest novel associations, such as the role of bZIPs or TCPs in -Fe responses. We assessed in the next paragraph in which specific -Fe response processes pCREs of particular interest, such as IDE1-likes, might be involved in.

### Associating important pCREs with FIT-dependent or FIT-independent functions

After identifying novel potential regulators in the -Fe response, we pinpointed some of those which could best explain models of -Fe-responsive up-regulation and explored their potential functions.

More than 1,500 pCREs were identified in total in GS-enriched clusters, raising the question of a core set of important pCREs needed to robustly predict -Fe response in each cluster. Using pCRE abundance (freq-pCREs) among GS-enriched clusters as the only criteria for selecting informative motifs for those clusters could result in missing motifs simply due to the fact that some co-expression clusters were more unique than others. This is supported by the fact that rare pCREs still can have a high importance rank (**Figure 4A**), meaning that those pCREs were not included in the set of freq-pCREs although they seem to be important for regulating individual GS-enriched clusters. We identified the most important pCREs (defined as the minimum set of pCREs; min-pCREs) by building RF models iteratively with successively deleting the least important pCREs in each round (**Supplemental Figure S6**). Applying this approach to the 159 GS-enriched clusters resulted in a collective set of 615 min-pCREs. They were part of the minimum sets of between 1 (0.6%) and 48 (30%) GS-enriched clusters, with the IDE1-like CATGCC being the top most abundant min-pCRE. Together with <u>CATGC</u>C, two more IDE1-like motifs, T<u>CATGC</u> and C<u>CATGC</u>, were among the top ten most abundant min-pCREs (**Supplemental Table S3**). This supports a previous computational analysis of rice promoters, in which IDE1-like was among the top scoring motifs (Kakei et al., 2013). Together with our previous finding typing IDE1-like ABI3VP1/B3 TFBMs to GS-enriched clusters, it suggests an important, yet unknown, function of IDE1-like motifs in Arabidopsis -Fe response regulation. Min-pCREs matching a bHLH (CGTGAC), a MYB (TAACTA), and the IDE2-like NAC TFBM (CACGCC; all **Supplemental Table S4A**) were also among the top ten most abundant min-pCREs (**Supplemental Table S3**), further demonstrating the utility of our approach.

To determine in which processes min-pCREs might function during -Fe, we tested if min-pCREs were more likely to be found in FIT-dependent or FIT-independent co-expression clusters. More than 60% of the 159 GS-enriched clusters were classified as either FIT-dependent with a likely function in root iron acquisition (35% out of 159) or FIT-independent with either a function in internal Fe homeostasis in shoots and roots or in -Fe response regulation (28% out of 159; **Figure 6A**). We then calculated the proportion of min-pCREs (present in ≥5 GS-enriched clusters) in each cluster category (**Supplemental Table S5**). Interestingly, the two IDE1-like motifs, C<u>ATGC</u>C, CC<u>ATGC</u> and the related ABI3VP1/B3 TFBM matched <u>ATGC</u>AT, were predominantly identified in FIT-dependent clusters, but IDE2-like CACGCC had no preference for either FIT-dependent or FIT-independent clusters (**Figure 6B**). This suggests that the IDE1-like pCREs tend to be more important for FIT-dependent root Fe acquisition rather than FIT-independent Fe sensing, signaling and distribution. This is also consistent with the role of grass IDE1 in Fe uptake (Kobayashi et al., 2003; Kobayashi et al., 2005). Two ARF TFBM matched min-pCREs (AACGTA/ARF16, GTCGGA/ARF2) were also preferentially found in FIT-dependent clusters. ARFs are involved in auxin signaling, thereby controlling - among other functions - root hair elongation (Pitts et al., 1998; Mangano et al., 2017; Choi et al., 2018). Different studies reported that -Fe responses can be accompanied by an increase of root hair number, elongation of root hairs, deformed or short root hairs (Schmidt et al., 2000; Müller and Schmidt, 2004; Dinneny et al., 2008). ARF2 and ARF16 TFs are root hair growth repressors (Choi et al., 2018), which would be consistent with a short root hair phenotype and down-regulation of respective GO terms under -Fe (Dinneny et al., 2008) (**Supplemental Figure S1**). During -Fe, several root hair-acting genes are co-expressed in a regulon which also contains *IRT2* (Ivanov et al., 2012), indicating a possible connection of ARF TFBM matched min-pCREs with these root hair processes.

**Figure 6.**
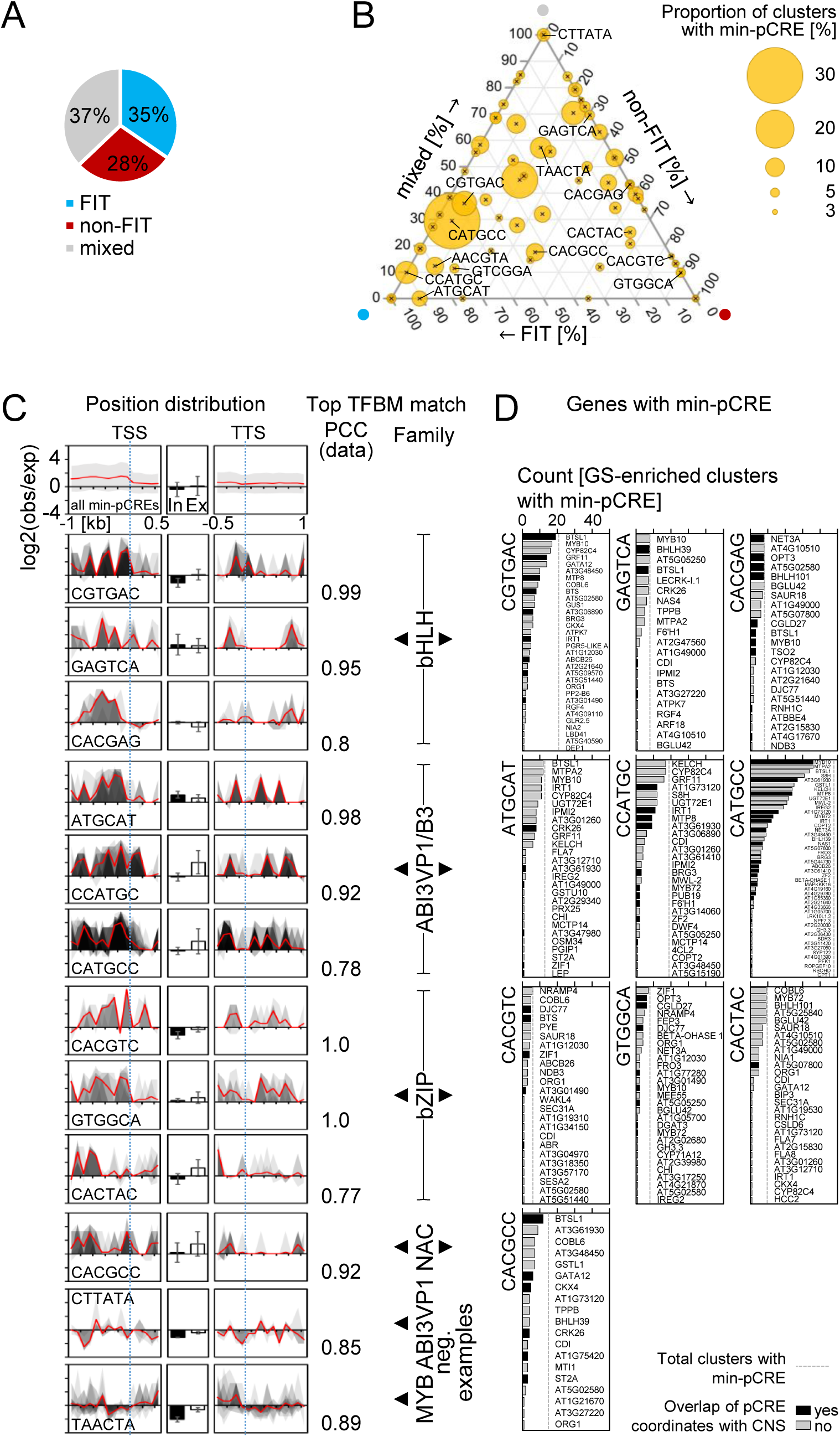
Characteristics of the most informative pCREs (min-pCREs). **A:** Proportion of GS-enriched co-expression clusters enriched for FIT-dependent genes (FIT, blue), FIT-independent genes (non-FIT, red) or both (mixed, gray). **B:** Ternary plot including min-pCREs identified in >3% (n=5) GS-enriched clusters. Position of the min-pCREs corresponds to the normalized proportions of FIT, non-FIT, and mixed clusters in which the min-pCRE was identified. Bubble size corresponds to the overall proportion of GS-enriched clusters with min-pCRE. Labeled min-pCREs are shown in 6C, D or mentioned in the main text. **C:** Positional bias of all (mean with standard deviation; **top**) and selected min-pCREs (**below**) in the putative promoter region (**1^st^ column**), all introns (In) and all exons (Ex) (**2^nd^ column**; mean with standard deviation), and in the putative non-coding region (**3^rd^ column**). 1^st^ and 3^rd^ column: position distributions in all co-expression clusters with min-pCRE (gray areas) with mean distribution (red line). TFBM matches (PCC) for each min-pCRE are shown (**4^th^ column**) and min-pCREs are sorted by TF family. log2(obs/exp): log_2_ of the number of observed (obs) min-pCRE occurrences divided by the number of min-pCRE occurrences in randomized sequences (expected, exp). **D:** Genes which might be regulated by the selected min-pCREs. Count: number of GS-enriched clusters in which the min-pCRE was identified and which included the respective gene having the min-pCRE in its promoter. Dashed line: total number of GS-enriched clusters with the min-pCRE. Genes in which the min-pCRE overlaps with a CNS are designated with black bars. A high-resolution image is available as **Supplemental Figure S7**.

In contrast, three bZIP TFBM matched min-pCREs, GTGGCA, CACGTC and CACTAC, were predominantly identified in FIT-independent clusters. As described in the previous section, bZIPs are involved in ABA signaling. ABA negatively regulates FIT-dependent Fe^3+^ reductase gene *FRO2* and Fe^2+^ importer gene *IRT1* (Séguéla et al., 2008). However, ABA signaling also leads to enhanced apoplastic and vacuolar Fe utilization and root to shoot transport under -Fe (Séguéla et al., 2008; Lei et al., 2014). (Lei et al., 2014) propose that ABA-responsive gene regulation and Fe remobilization and transport are connected through bZIPs, which is consistent with our results that bZIP TFBMs are preferentially found in FIT-independent co-expression clusters of genes involved in Fe mobilization and translocation. Similarly, two TCP TFBMs, GACCAC and ACCCAC, were identified almost exclusively in FIT-independent clusters, which is in agreement with TCP20 regulating *BHLH39* in a FIT-independent manner (Andriankaja et al., 2014). Finally, bHLH TFBMs were identified in all cluster categories with a preference for mixed clusters. This matches the ubiquitous nature of bHLH target motifs (E-/G-boxes), which act at many levels in the -Fe bHLH cascade. In summary, we can propose plausible roles for pCREs as TFBMs in FIT-dependent and FIT-independent processes.

### Distribution and conservation of min-pCREs in co-expression cluster gene promoters

Next, to explore if min-pCREs displayed significant positional bias in the promoter regions of co-expressed genes, we compared the observed min-pCRE frequencies in 100 bp bins of −1000 to +500 bp and of −500 to +1000 bp flanking regions adjacent to the transcription start site (TSS) and transcription termination site (TTS), respectively, with the expected frequencies from shuffled pCRE sequences (according to Uygun et al., 2017) for all 615 min-pCREs (**Supplemental Figure S9**). Furthermore, we examined the non-coding as well as the coding sequences of the transcribed regions. Overall, the distributions of min-pCREs revealed a slight positional bias in the promoter regions (**top**, **Figure 6C**). We investigated the distribution plots separately for ten selected min-pCREs: FIT-dependent ABI3VP1/B3 TFBM-matched min-pCREs (containing the two IDE1-likes), the IDE2-like (NAC TFBM match), FIT-independent bZIP TFBM matches, and bHLH TFBM matches (**Figure 6C**). These min-pCREs had significant location bias in the putative promoters up to 1000 bp upstream of the TSS. Because known CREs often exhibit positional bias (Zou et al., 2011; Heyndrickx et al., 2014; Yu et al., 2016), this provides additional support for these pCREs having regulatory functions in Fe uptake and homeostasis. Interestingly, some of the pCREs common to mixed clusters did not show position bias in any of the genomic regions tested (e.g. ABI3VP1/CTTATA and MYB/TAACTA; **bottom**, **Figure 6C**), indicating that genes of such clusters are less likely to be transcriptionally co-regulated.

Next, we sought to pinpoint the specific processes of Fe homeostasis, which these ten min-pCREs might regulate. To assess this, we determined the genes containing the respective min-pCRE and counted the number of incidents in which these genes were likely regulated by the min-pCRE (**Figure 6D**; **Supplemental Table S6A**). For example, CATGCC was found in 48 GS-enriched clusters, and 36 of those clusters (75%) included the min-pCRE-containing gene *MYB10*, while only four of those clusters (8%) included *MAPKKK16* (**second row right**, **Figure 6D**). We inferred that CATGCC might be regulating predominantly processes in which MYB10 is required. As an additional line of evidence for pCRE functionality, we determined if min-pCREs overlapped with conserved noncoding sequences (CNS) of the Brassicaceae family (Haudry et al., 2013) (**Supplemental Table S6B**). As expected, bHLH TFBM matched min-pCREs were identified in many gene promoters, including *BHLH39*/*BHLH101* (direct targets of bHLH IVc TFs, (Zhang et al., 2015)), *NAS4* (direct PYE target, (Long et al., 2010)), and *IRT1*, *AT3G07720* and *GRF11* (FIT targets; **Figure 6D**; (Sivitz et al., 2012; Yang et al., 2013)). In *BHLH39*/*BHLH101*, *IRT1*, and *GRF11*, the respective min-pCREs overlapped with CNS, further supporting the importance of these motifs. Interestingly, bHLH matched min-pCREs were also located in CNS of *BTS* and *BTSL1*, two genes that negatively regulate Fe uptake by marking positive regulators (e.g. bHLH IVc TFs) for degradation (Selote et al., 2015). If *BTS* were to be regulated by bHLH proteins from the same regulatory cascade, this may indicate a negative feedback loop.

FIT-dependent IDE1-likes CATGCC and CCATGC were located in the *IRT1* promoter, overlapping with CNS (**second row**, **Figure 6D**). Interestingly, both IDE1-likes were found in several genes encoding enzymes and TFs involved in coumarin biosynthesis (*CYP82C4*, *S8H*, *F6′H1*, *MYB72*/*MYB10*; (Kai et al., 2008; Murgia et al., 2011; Fourcroy et al., 2014; Schmid et al., 2014; Zamioudis et al., 2014; Rajniak et al., 2018; Siwinska et al., 2018)). We propose that IDE1 is important for synthesis of Fe chelators in response to low Fe conditions in both monocots and dicots (see Kobayashi et al., 2003; Kobayashi et al., 2005). The IDE2-like min-pCRE was located amongst others in *BTSL1* (within a CNS), *BHLH39*, *ORG1*, and a number of uncharacterized genes. While the role of IDE2 in the Strategy II Fe response has not been comprehensively explored, it is known to regulate expression of *OsYSL2*, a phloem Fe^2+^-nicotianamine transporter (Kobayashi et al., 2003; Ogo et al., 2008). This putative involvement in phloem translocation of Fe suggests that the IDE2-like might preferably associate with FIT-independent gene functions. However, we could not assess the relationship between IDE2 and *YSLs* in this analysis because *YSL1*/*2*/*3* were not expressed above the log2FC>1 threshold.

Next, we explored the bZIP TFBM-matched min-pCREs, since they had the strongest FIT-independent preference. These min-pCREs were located in a number of genes involved in translocation of Fe/Fe chelates or the synthesis of Fe chelators (e.g. *NRAMP4*, *ZIF1*, *OPT3* (Lanquar et al., 2005; Haydon et al., 2012; Mendoza-Cózatl et al., 2014; Zhai et al., 2014)) or in Fe sensing and signaling (e.g. *OPT3*, *BTS*, (Mendoza-Cózatl et al., 2014; Zhai et al., 2014; Selote et al., 2015; Khan et al., 2018)). Furthermore, GTGGCA (matched to bZIP TFBM) overlapped with a CNS of *CGLD27* (**third row**, **Figure 6D**), which has been associated with photoprotection in leaves during -Fe (Ruiz-Sola and Rodríguez-Concepción, 2012; Rodríguez-Celma et al., 2013). However its function in roots remains elusive. Taken together, our findings suggest diverse roles for bZIP TFBMs, including Fe transport and adjustment of the plastid proteome.

In summary, we identified more than 100 -Fe pCREs which, in addition to sharing significant sequence similarity to known TFBMs, were also part of the core sets of pCREs needed for robust prediction of -Fe responses of GS-enriched clusters (min-pCREs). Furthermore, they were preferentially located in promoter regions upstream of the TSS, and even in CNS’ of some genes. Together, these findings indicate that these pCREs might be authentic -Fe CREs. From the biological context of the genes which are likely regulated by some of the pCREs, we were able to greatly improve our understanding of -Fe response regulation in Arabidopsis. For example, our work highlighted that in addition to the bHLH TFBMs, IDE1-like motifs and bZIP TFBMs are likely involved in different responses to -Fe and should be considered of high interest for future work.

## Conclusion

We identified 5,639 pCREs enriched in promoters of co-expressed -Fe-responsive genes that were used as features to predict -Fe-responsive regulation of root-expressed genes on a genome-wide scale. Of those, 173 reliably predicted -Fe response genes of >5% of our defined co-expression clusters (freq-pCREs). Because most of those pCREs were either unique to co-expression clusters enriched for our gold standard Fe acquisition and homeostasis genes, or unique to co-expression clusters lacking those genes, we conclude that our approach had captured motifs specifically regulating different responses during -Fe. To take advantage of the publicly available *in vitro* TF binding information, we compared the freq-pCREs to TFBMs from two studies (Weirauch et al., 2014; O’Malley et al., 2016), and found that our approach had captured known Strategy I -Fe recognition motifs for bHLH and MYB proteins. Our approach also led to novel regulatory connections of bZIP, B3, NAC, and TCP families to Strategy I -Fe response regulation. While bZIP and bHLH TF families are also associated with high salinity stress response (Uygun et al., 2017), other high salinity stress response associated TF families (e.g. WRKY and AP2) were not common among our -Fe pCREs regulating GS-enriched co-expression clusters, highlighting the usefulness of this approach to pinpoint regulators specific to a stress condition.

We inferred possible functions of pCREs which were most important for modeling -Fe responses (min-pCREs) from their enrichment in FIT-dependent or FIT-independent co-expression clusters and their location bias in promoters of particular -Fe-responsive genes (**Figure 6B, D**). Our results provide evidence that B3 TFBM pCREs containing the IDE1 core motif CATGC are linked to coumarin synthesis, indicating that the function of IDE1-like motifs to ensure supply of Fe-chelating compounds for Fe acquisition could be an evolutionarily conserved function at least among flowering plants. While our results highlight the importance of IDE1-like motifs for Fe acquisition, it was not the only prominent -Fe pCRE. This is in contrast to Zn deficiency, where ZDRE seems to be singularly associated with multiple Zn deficiency responses (Assunção et al., 2010), and indicates that despite of overlaps of Zn and Fe homeostasis control (Briat et al., 2015), their transcriptional regulation must follow different mechanisms.

Our results support a concept in which -Fe is not regulated by only one or few regulatory elements. Of the many important pCREs for -Fe response, many share significant similarity with TFBMs of TF families known to undergo hetero-dimerization and protein interaction across families, such as bHLH, MYB, bZIP, TCP, and ABI3VP1 (Bemer et al., 2017). A combinatorial mechanism would dramatically increase the flexibility of transcriptional responses driven by a set of few TFs. It might be that some pCREs not as important in our prediction models, would become informative in combination, as suggested for high salinity stress response (Zou et al., 2011; Uygun et al., 2017). A next step would therefore be to build -Fe prediction models that explicitly account for interactions between pCREs. A limitation of our approach is that our co-expression clusters were based on ATH1 chip microarray data, the only comprehensive -Fe time course transcriptome set available to date. Some important -Fe marker genes (e.g. *FRO2*) are not represented on the chip and others might not have passed our significance threshold because of sensitivity issues with the microarray technology. Additionally, we restricted our analysis to the promoter region 1000 bp upstream of the TSS. While this is expected to cover most important *cis*-acting elements and reduce the occurrence of promoters overlapping with adjacent genes, introns as well as more distal promoter regions are known to harbor *cis*-acting elements (Rose et al., 2008; Rose et al., 2016).

The large number of TFs known to be involved in -Fe-induced up-regulation points towards the importance of transcriptional regulation. However -Fe responses are also heavily controlled at the post-transcriptional and post-translational level (Lingam et al., 2011; Meiser et al., 2011; Sivitz et al., 2011; Selote et al., 2015; Zhang et al., 2015; Gratz et al., 2019). Naturally, our approach cannot cover such regulatory aspects. However, it allows us to predict TF families, that may act upstream of the known -Fe-responsive genes. We suggest TFs of the bZIP, ABI3VP1/B3, NAC, and TCP families as upstream regulators of -Fe response in the root. Because major Fe sinks are located in the shoot, a systemic shoot-to-root signal must exist for proper Fe supply (Vert et al., 2003; Garcia et al., 2013). Integrating shoot transcriptomic data would expand our knowledge on how responses to -Fe stress are coordinated at the whole-plant level.

In conclusion, we demonstrate that our machine learning-based approach can identify pCREs for -Fe-induced gene up-regulation. This strategy can be applied to various stresses and developmental conditions to elucidate regulatory mechanisms, especially when *cis*- and/or *trans*-acting elements were previously elusive (Zou et al., 2011; Uygun et al., 2017). We provide a comprehensive source of potential -Fe response *cis*-regulators for a wide range of -Fe-responsive genes. Because the identified pCREs are potentially involved in enhancing Fe uptake and translocation, they generate potential for future applications in engineering plants with improved plant performance traits, e.g. higher nutritional value because of better Fe allocation and coping with unfavorable soil conditions.

## Methods

### Expression data processing and generation of multiple expression data combinations

Expression data (Affymetrix ATH1) from an -Fe treatment time course experiment with six time points and of four -Fe treated root zones (both Dinneny et al., 2008) were downloaded from Gene Expression Omnibus (GEO, https://www.ncbi.nlm.nih.gov/geo/, GSE10502, GSE10497) in CEL format, preprocessed, normalized and contrasted as described below. AtGenExpress expression data (Affymetrix ATH1) of abiotic stresses ((Kilian et al., 2007), GSE5620-5628 or TAIR-ME00325-330, only data of root samples were used), hormone treatment ((Goda et al., 2008), GSE39384 or TAIR-ME00333-340, ME00343-344, ME00350-352, ME00356) and plant development ((Schmid et al., 2005), GSE5629-5634 or TAIR-ME00319) were downloaded from The Arabidopsis Information Resource (TAIR; https://www.arabidopsis.org/portals/expression/microarray/ATGenExpress.jsp) preprocessed, normalized and contrasted by S. Uygun (Uygun et al., 2016) as described below. Background correction and quantile normalization of CEL files were performed with Robust Multi-Array Average expression measure (RMA) using the Bioconductor affy package (Gautier et al., 2004). The log_2_ fold-change (log2FC) in expression was calculated for all data sets except developmental data by pairwise comparison of treatment and control experiments for each treatment and time point. Contrast matrices and linear model fits were created using R and the Bioconductor LIMMA package (Ritchie et al., 2015; Phipson et al., 2016). Because developmental stages have no control treatment, absolute normalized fluorescence intensity values were used. The *p*-values for log2FC or fluorescence intensities were corrected for multiple testing (adjusted *p*-values=*q*) using the BH method (Benjamini and Hochberg, 1995). Genes were regarded as -Fe responsive if abs(log2FC)≥1, and *q*<0.05 at least at one -Fe treatment time point or in at least one -Fe treated root zone. -Fe deficiency time course data was combined with -Fe root zone expression data or ATGenExpress datasets in different combinations (**Figure 1C**) either including only genes up-regulated (“up”) or all genes up- or down-regulated (“up & down”) in ≥1 -Fe time point or root zone. This resulted in 12 different expression data combinations.

### Co-expression clustering using *k*-means

To cluster genes with similar expression pattern, *k*-means clustering (Hartigan and Wong, 1979) was applied using the Euclidean distance as the similarity measure. Because *k*-means returns a local optimum solution depending on the number of clusters (*k*) created and the random selection of genes as initial “means”, the outcome varies with run (i.e. non-deterministic). Therefore, different *k* (25, 30, 35, 40, 50, 70, 80, 100) were tested and the clustering was repeated up to four times. We build machine learning models (see below) with all clusters generated from expression data combinations (DC) 1, 2, 3 and 5 (**Figure 1C**). To prevent confusion, we point out that the total number of *k*-means-generated clusters used to build models represents several repeated clustering events of always the same two sets of - Fe-responsive genes (up; up & down, see above). The clustering events differ in the DC which was used and in the *k*. Two DC were excluded from the analysis: DC 4 produced identical clusters as DC 1, which were therefore not considered. DC 6 contained different measuring units (log2FC and absolute normalized fluorescence intensity), and could not be handled by the *k*-means algorithm.

### Co-expression clustering by correlation with GS genes

To generate co-expression clusters based on gold standard (GS) genes (**Supplemental Table S1**), each GS gene was used as a query to identify genes with similar expression patterns. Briefly, for each expression data combination (DC; **Figure 1C**), the PCC was calculated between the query gene and each gene in DC using SciPy (http://www.scipy.org, (Jones et al., 2001)). Similar to (Uygun et al., 2016), a random background PCC was calculated representing the null distribution of expression correlation by calculating the PCC of 10,000 randomly selected gene pairs in DC and the 95^th^ percentile of PCCs was tried as the threshold for classifying a pair of genes as significantly correlated. For some DC (mostly those containing only up-regulated -Fe responsive genes), we allowed a significance threshold below 90% down to 45%, because the PCC between random Fe responsive genes was already very high. On the other hand, when >50 genes were considered significantly correlated, the threshold was raised above 95% to 99% to hone in on genes most likely to be co-regulated. In addition, we generated a second version of clusters with >50 genes, containing only the 10 genes with highest PCC. We build machine learning models (see below) with both versions and further used the results from the better performing version only. Percentiles used for each PCC-generated cluster are given in **Supplemental Table S2**. Two DC were excluded from the analysis: DC 4 (up), because the resulting clusters were identical to those generated from DC 1 (up), and DC 6 (up & down), because developmental data seemed to have a disproportional influence on the PCC with the result that even -Fe up- and down-regulated gene pairs were identified as strongly correlating. As in the *k*-means clustering, the total number of PCC-generated clusters used for modeling represents repeated clustering events of the same two sets of -Fe-responsive genes (as described above).

### The co-expression clusters: GO and GS/FIT-dependent/FIT-independent gene enrichment and GO/gene content similarity

Gene ontology (GO) associations for *A. thaliana* were downloaded from TAIR (ftp://ftp.arabidopsis.org/home/tair/Ontologies/Gene_Ontology/ (Berardini et al., 2004)). Biological process (BP) GO annotations were downloaded from GO (http://purl.obolibrary.org/obo/go.obo) and parsed for BP information. Enrichment of GO terms in genes that were significantly differentially regulated (*q*<0.05, abs(log2FC)≥1) in the -Fe time course data set was determined with a Fisher’s exact test (FET, http://www.scipy.org, (Jones et al., 2001)), and *p*-values were corrected for multiple testing (=*q*) using the “qvalue” function in R (Storey, 2002) (**Supplemental Table S7**).

All co-expression clusters were tested for enrichment of GO terms as described above. The similarity of enriched GOs between co-expression clusters was assessed using the Jaccard Index (JI), or the intersection of GOs divided by the union of the GOs, where JI=1 if the exact same GOs were enriched in both co-expression clusters. Co-expression clusters were grouped by hierarchical clustering using the JI with the UPGMA method in the R cluster package (Maechler et al., 2017). Groups containing >20 co-expression clusters and having a within-mean JI that was significantly higher than the mean JI of all clusters were defined as “superclusters”. Biological functions of superclusters were defined through GOs shared by ≥75% (*k*-means clustering) or ≥90% (GS gene correlation; PCC) of the clusters. Similarly, FET with *p*-value correction for multiple testing was used to identify co-expression clusters enriched for (A) -Fe GS genes, (B) FIT-dependent genes, and/or (C) FIT-independent genes.

### *K*-mer enrichment and identification of pCREs predictive of -Fe response using Random Forest (pCRE identification pipeline)

Promoter sequences 1 kb upstream from the transcription start site (TSS) were downloaded from TAIR (ftp://ftp.arabidopsis.org/home/tair/Sequences/blast_datasets/TAIR10_blastsets/upstream_sequences/TAIR10_upstream_1000_20101104). A list of all possible 6-mers of A, T, C, G was generated with the Python itertools function and the Biopython Bio.Seq module (http://biopython.org/ wiki/Biopython, (Cock et al., 2009)). Only one 6-mer for each reverse complement pair was kept (resulting in 2,080 6-mers). Genes were considered -Fe non-responsive if they were not significantly differentially expressed (abs(log2 FC) <0.4) during any time point during the -Fe time course experiment or in any -Fe treated root zone or in four additional -Fe treatment experiments ((Li and Schmidt, 2010): GSE16964, (Long et al., 2010): GSE21443, (Schuler et al., 2011): GSE24348, (Sivitz et al., 2012): GSE40076). The four additional data sets were downloaded in CEL format from GEO and processed as described in the first section of the **Methods** part.

Potentially meaningful *cis*-regulatory elements for -Fe response were identified in two steps, where we first looked for enriched *k*-mers in the promoters of -Fe responsive genes and then determined how well the enriched *k*-mers predicted -Fe response using machine learning. The code for this analysis is available on GitHub (https://github.com/ShiuLab/MotifDiscovery, https://github.com/ShiuLab/ML_Pipeline). For the first step, the promoter sequences of the genes in co-expression clusters (positive set) were searched for enriched 6-mers in comparison to promoter sequences of non-responsive genes (negative set). These enriched 6-mers were elongated by one base and tested again for enrichment. This process was repeated until no longer *k*-mer was more enriched than the shorter *k*-mer. Enrichment was calculated using a one-sided FET (*p*<0.01).

For the second step, to determine which sets of enriched *k*-mers were predictive of -Fe response, we generated features based on presence or absence of each enriched *k*-mer and used these features to build machine learning models using the Random Forest (RF) algorithm (Pedregosa et al., 2011). To avoid building biased models, 50 models were generated for each co-expression cluster by randomly drawing from the negative set to generate balanced (i.e. size positive set equals size of negative set) input datasets. A 10-fold cross-validation approach was used to train and test the models. Briefly, the balanced datasets were divided randomly into ten even groups with a 1:1 ratio of positive to negative class genes. The model was trained on the 1-9 folds and applied to the 10^th^ (and successively trained on 1-8+10 and applied to the 9^th^, etc.). This cross-validation scheme was repeated ten times. Each RF model was made up of 500 decision trees, each trained on a random subset of enriched *k*-mers and of training set genes. The final model performance is represented by the mean F1 score (i.e. F-measure) across all 50 balanced models. The F1 score is the harmonic mean of precision (P=TP/(FP+TP)) and recall (R=TP/(FN+TP)), where TP=true positive, FP=false positive, and FN=false negative. Only co-expression clusters for which the enriched *k*-mers were deemed as good predictors (F1≥0.7) were used in the downstream analysis.

Predictive *k*-mers (then referred to as putative *cis*-regulatory elements, pCREs) were ranked by importance. The importance score is based on the Gini Index, which is a measure of node purity, where important pCREs separate positive from negative class genes well and low ranked pCREs are less informative. To determine how well the models predicted specific -Fe responsive genes, we calculated the percent of times each gene was correctly predicted (TP) out of the 50 balanced replicates.

### pCRE sequence similarity

To assess sequence similarities between the 173 pCREs that were frequently identified (in >5%; freq-pCREs) in GS-enriched or non-enriched co-expression clusters with good model performance (F1≥0.7), sequence dissimilarity of pCRE position weight matrices (PWMs) was calculated by pair-wise PCC distance and a distance matrix was generated using the TAMO package (Gordon et al., 2005). Freq-pCREs were grouped by hierarchical clustering of the PCC distance matrix using the UPGMA method in the R cluster package (Maechler et al., 2017), and visualized in a dendrogram (**Supplemental Figure S4E**). Due to group-wise averaging of PCC distances during hierarchical clustering, the algorithm produced skewed PCC distances of some similar pCRE pairs. Therefore, freq-pCRE clusters were additionally visualized as a network, in which freq-pCREs with PCC distance=0 (identical freq-pCREs or subsets of each other) were connected with black bold edges and freq-pCREs with PCC distance≤0.22 were connected with light gray edges (**Figure 4C**). Highly interconnected nodes were arranged in groups. The network was created using the Cytoscape software (Shannon et al., 2003). To show a consensus of freq-pCREs within a network group, freq-pCRE sequences were aligned using ClustalX (Larkin et al., 2007) with default parameters and a sequence logo was created with weblogo (https://weblogo.berkeley.edu/logo.cgi).

### Identification of most informative pCREs (min-pCREs) by non-linear regression

The most informative pCREs of a co-expression cluster were defined as the minimum set of pCREs (min-pCREs) needed for RF models without sacrificing performance. To identify min-pCREs, for each GS-enriched co-expression cluster, the pCREs used as features were step-wise reduced, with the least important pCREs deleted at each step. First, for pCREs that were subsets of each other (PCC distance=0), the lower ranked one was removed. Then, from this list of pCREs and for successively shorter lists of pCREs (n=40, 30, 25, 20, 15, 12, 10, 8, 6, 5, 4, 3, 2, 1), 10 replicates of RF models were trained on balanced datasets. F1 scores were plotted against the number of pCREs (x) and a non-linear regression curve was fitted to the data points. An exponential recovery function

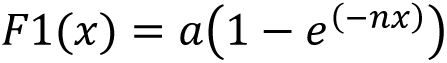

was found to best describe the data behavior. Starting values for variables a and n were approximated by fitting a linear model to the logarithmic transformation of the function. The set of pCREs with the highest F1 closest to the inflection point of the regression curve was defined as min-pCRE set (example in **Supplemental Figure S6**).

### pCRE similarity to TFBMs

*In vitro* binding data of Arabidopsis TFs to genomic DNA (DNA Affinity Purification Sequencing, DAP-seq, (O’Malley et al., 2016)) and TF binding data based on protein-binding microarray data or the TRANSFAC data base (Catalog of Inferred Sequence Binding Preferences, CIS-BP, (Weirauch et al., 2014)) were used (**Supplemental Table S4B**), with DAP-seq TFBMs used over CIS-BP TFBMs when the TF was present in both databases. PWMs of pCREs were compared to PWMs of TFBMs using PCC and the pCREs were classified as similar to (A) a specific TF, (B) a TF family, or (C) to TFs generally, based on the degree of similarity to their best matching TFBM (Uygun et al., 2017). A pCRE was similar to a specific TFBM (A) if the PCC between the pCRE and the TFBM was ≥95^th^ percentile of PCCs between that TFBM and TFBMs from the same TF family. Alternatively a pCRE was similar to TFBMs from a TF family (B) if the PCC between the pCRE and a TFBM from that family was ≥95^th^ percentile of PCCs between TFBMs from that family and TFBMs from other TF families. Finally, a pCRE was similar to TFBMs (C) if the PCC between the pCRE and any known TFBM was ≥95^th^ percentile of PCCs between TFBMs and randomly generated 6-mers. For 95^th^ percentile PCC thresholds see **Supplemental Table S4C**. To determine if pCREs similar to specific TF families were enriched in GS-enriched versus non-enriched co-expression clusters, the percentage of pCREs similar to TFBMs (significance level A or B) from each TF family was calculated for each co-expression cluster category. Then, FET with multiple testing correction (*q*≤0.05) was used to determine if GS-enriched co-expression clusters were enriched for TF families compared to non-GS-enriched co-expression clusters and vice versa.

### Positional distribution of pCREs

To determine the positional distribution of the min-pCREs for each GS-enriched co-expression cluster, min-pCREs were converted to PWMs adjusted to the Arabidopsis background AT (0.33) and CG (0.17) content using the TAMO package (Gordon et al., 2005) and mapped to the promoter sequences ranging from 1000 bp upstream to 500 bp downstream of the transcription start site (1000-TSS-500), using Motility (http://cartwheel.caltech.edu). For comparison, min-pCRE PWMs were also mapped to exons and introns, respectively, and to the region 500 bp upstream and 1000 bp downstream of the transcription termination site (500-TTS-1000). Arabidopsis sequences were downloaded from TAIR (ftp://ftp.arabidopsis.org/Sequences/blast_datasets/TAIR10_blastsets/). Positional distributions were calculated as described in (Uygun et al., 2017). In brief, min-pCREs were mapped to 100 bp bins of 1000-TSS-500 and 500-TTS-1000 and to whole exons and introns. For comparison, min-pCREs were mapped to randomized versions of the sequences. Randomization was performed within each 100 bp bin and in each exon or intron, respectively, in order to maintain nucleotide composition and therefore GC content. Positional distribution was calculated as log2FC of number of observed mappings divided by number of randomly expected mappings (log2FC(observed/expected)).

### pCRE coordinate overlap with CNS coordinates

All 615 min-pCRE PWMs were mapped to the putative promoter region (1 kb upstream of TSS) of -Fe response genes (described above). The min-pCRE coordinates were then compared to the coordinates reported as conserved non-coding sequences (CNS) across nine species in the Brassicaceae family (Haudry et al., 2013) downloaded from the UCSC Genomics Bioinformatics website (http://mustang.biol.mcgill.ca:8885/download/A.thaliana/gff/AT_CNS.gff; **Supplemental Table S6B**).

## Supporting information

Supplemental_Figures_S1-S7_Supplemental_Table_S1_Supplemental_literature

## Author contributions

B.S., P.B., and S.-H.S. conceived the project; B.S., and S.-H.S. designed the research plan; B.S., and C.B.A. analyzed the data; B.S. wrote the original draft; B.S., C.B.A., S.-H.S., and P.B. reviewed and edited the article; P.B., C.B.A, and S.-H.S. acquired funding. P.B. and S.-H.S. agree to serve as the authors responsible for contact and ensure communication.

## Abbreviations

CIS-BP: Catalog of inferred sequence binding preferences
CNS: Conserved non-coding sequence
DAP-seq: DNA affinity purification sequencing
Fe/-Fe: Iron/Iron deficiency
FeS: Iron-Sulfur
FET: Fisher’s exact test
FIT: FE DEFICIENCY-INDUCED TRANSCRIPTION FACTOR
freq-pCRE: Frequent pCRE
GO: (Biological process) Gene ontology
GS: Gold standard
IDE: Iron Deficiency-responsive Element
log2FC: log_2_ fold-change
MA: Mugineic acid
min-pCRE: Minimum set pCRE
PCC: Pearson’s correlation coefficient
pCRE: Putative *cis*-regulatory element
PWM: Position weight matrix
RF: Random Forest
TF: Transcription factor
TFBM: Transcription factor binding motif
TSS: Transcription start site
TTS: Transcription termination site
Zn: Zinc

## Supplemental Material

The following supporting material is available as three supplemental PDF files (1: **Supplemental_Figures_S1-S7_Supplemental_Table_S1_Supplemental_literature**; 2: **Supplemental_Figure_S8**; 3: **Supplemental_Figure_S9**), and as supplemental Excel spreadsheet (**Supplemental_Table_S2**-**S7_spreadsheet**).

**Supplemental Figure S1.** Complete GO enrichment analysis of -Fe-responsive genes.

**Supplemental Figure S2.** GO terms and -Fe GS gene enrichments of the defined co-expression clusters containing up-/down-regulated genes, and mean similarity within designated superclusters of up-regulated and up-/down-regulated genes.

**Supplemental Figure S3.** Co-expression cluster RF model performance of cluster category (GS-enriched and non-enriched) and cluster size.

**Supplemental Figure S4.** Comparison of the pCRE abundance and importance in GS-enriched clusters vs. non-enriched clusters and hierarchical clustering of freq-pCRE sequences.

**Supplemental Figure S5.** Significance of sequence similarity for freq-pCRE from non-enriched clusters and the best matching known TFBM.

**Supplemental Figure S6.** Example of a non-linear regression curve to determine the minimum set of pCREs for a co-expression cluster.

**Supplemental Figure S7.** High-resolution image of **Figure 6**.

**Supplemental Figure S8.** Expression plots of all well-performing co-expression clusters.

**Supplemental Figure S9.** Positional distribution plots of all 615 min-pCREs.

**Supplemental Table S1.** Robust -Fe-responsive GS genes (FIT-dependent/FIT-independent).

**Supplemental Table S2.** Detailed information of all generated co-expression clusters: input expression data combinations, algorithm and parameters used for clustering, enrichment of GS genes, FIT-dependent/FIT-independent/both genes, F1 score, gene content, all identified pCREs, min-pCREs.

**Supplemental Table S3.** List of all pCREs (n=5,639) identified in well-performing clusters (GS-enriched and non-enriched).

**Supplemental Table S4. A:** List of most relevant pCREs (freq-pCREs and min-pCREs) and their similarity to DAP-seq and CIS-BP TFBMs. **B:** DAP-seq and CIS-BP TFBMs used in this study. **C:** TF family 95^th^ percentiles of within, between and random PCC thresholds determining the pCRE-TFBM similarity.

**Supplemental Table S5.** Association of min-pCREs to FIT-dependent, FIT-independent or both cluster categories.

**Supplemental Table S6. A:** List of all 615 min-pCREs with total counts of GS-enriched clusters and genes having the min-pCRE. **B:** Overlap of min-pCRE coordinates with Brassicaceae conserved non-coding sequences (CNS).

**Supplemental Table S7.** *p*- and *q*-values of complete GO enrichment analysis of -Fe-responsive genes (**Supplemental Figure S1**).

## Acknowledgements

We thank Sahra Uygun, Bethany Moore, Nicholas Panchy, and Peipei Wang for providing scripts and help with programming. B.S. is a member of the international graduate school iGRAD-Plant, Düsseldorf. Funding from the German Research Foundation through the DFG International Research Training group 1525 to P.B. is greatly acknowledged. C.B.A. was supported in part by an NSF Graduate Research Fellowship. This work was partly supported by grants to S.-H.S. from the US National Science Foundation IOS-1546617, DEB-1655386, and DGE-1828149, and the US Department of Energy (DOE) Great Lakes Bioenergy Research Center (DOE Office of Science BER DE-SC0018409). This work received funding from Germany’s Excellence Strategy, EXC 2048/1, Project ID: 390686111.

