## Supplemental_Figure_S8 for "Putative *cis*-regulatory elements predict iron deficiency responses in Arabidopsis roots"

**Figure S8. Expression plots of all well-performing (F1≥0.7; n=517) co-expression clusters.**

Clusters are organized according to transcriptomic data combinations (**Figure 1C**). GS-enriched (n=159) and non-enriched (n=358) clusters of each data combination are shown on separate pages. Each cluster plot is labeled with its cluster ID, as in **Supplemental Table S2**, and with its enrichment category (FIT-dependent/FIT, FIT-independent/non-FIT, both/mixed, none (if not GS-enriched)). **Pages 1, 2**: -Fe time course[1] data only (Combination 1**). Pages 3, 4**: Time course + -Fe-treated root zones 1-4[1] (Combination 2). **Pages 5, 6**: Time course + hormones[3] (Combination 4). **Pages 7-10**: Time course + abiotic stresses[2] (Combination 3). **Page 11**: Time course + abiotic stresses + hormones (Combination 5a). **Pages 12-15**: Time course + abiotic stresses (genotoxic data deleted) + hormones (Combination 5b). **Pages 16, 17**: Time course + abiotic stresses + development[4] (Combination 6). Total number of plots: 517 (159 GS-enriched, 358 non-enriched). Blue dotted lines separate transcriptomic data within combinations. [1](Dinneny et al., 2008), [2](Kilian et al., 2007), [3](Goda et al., 2008), [4](Schmid et al., 2005). Units: for all data except development: $\log_2$ fold-change (log2FC); development: fluorescence intensity.

### Fe deficiency time course, GS-enriched

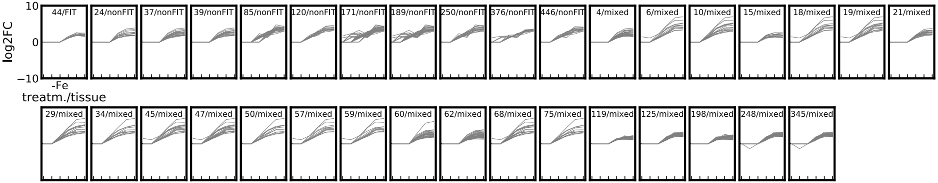

### Fe deficiency time course, non-enriched

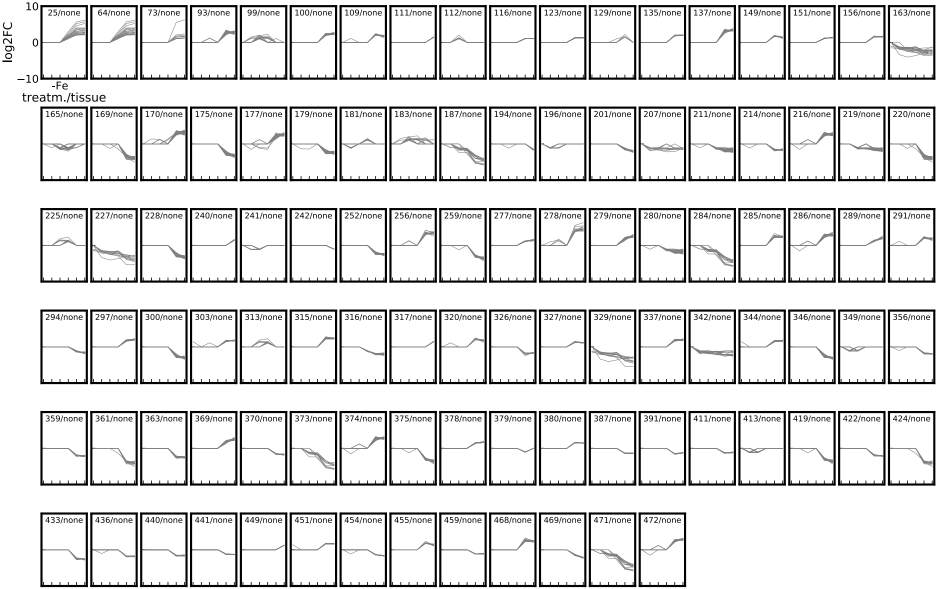

### Fe deficiency time course/root zones, GS-enriched

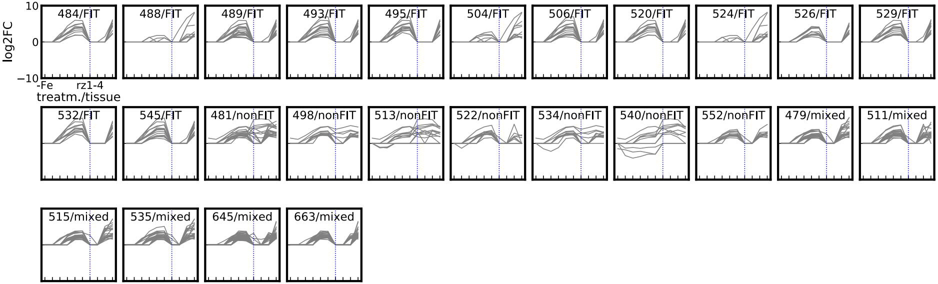

### Fe deficiency time course/root zones, non-enriched

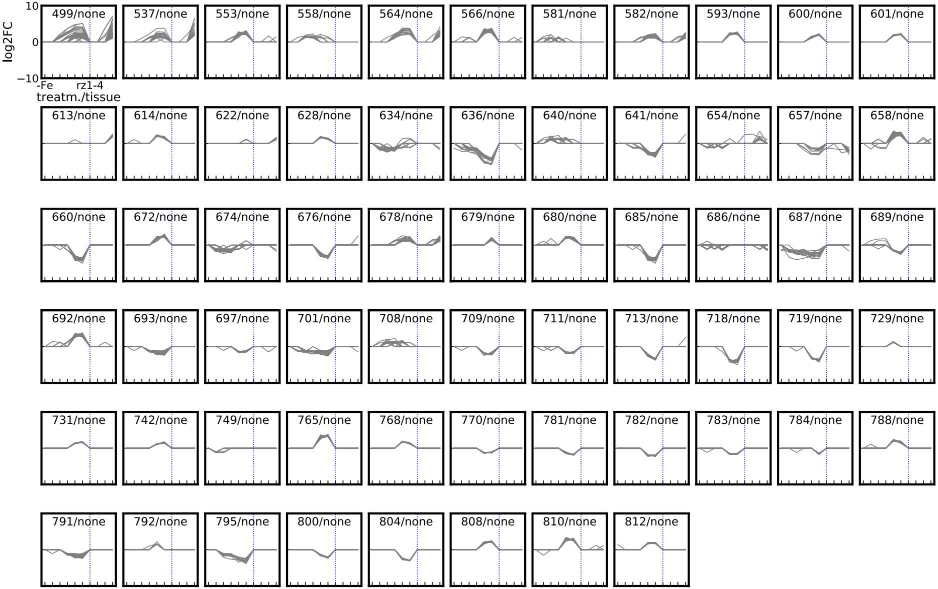

### Fe deficiency time course/hormones, GS-enriched

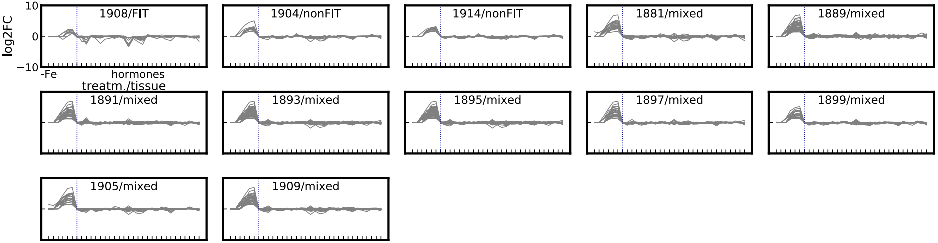

### Fe deficiency time course/hormones, non-enriched

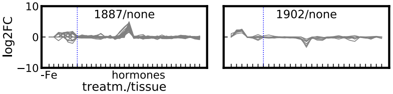

### Fe deficiency time course/abiot. stresses, GS-enriched

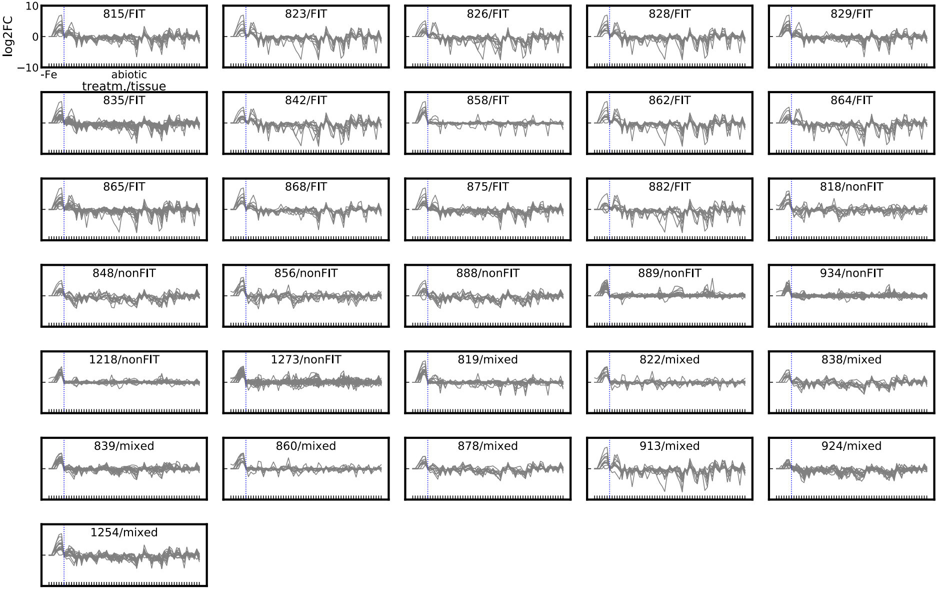

### Fe deficiency time course/abiot. stresses, non-enriched (1)

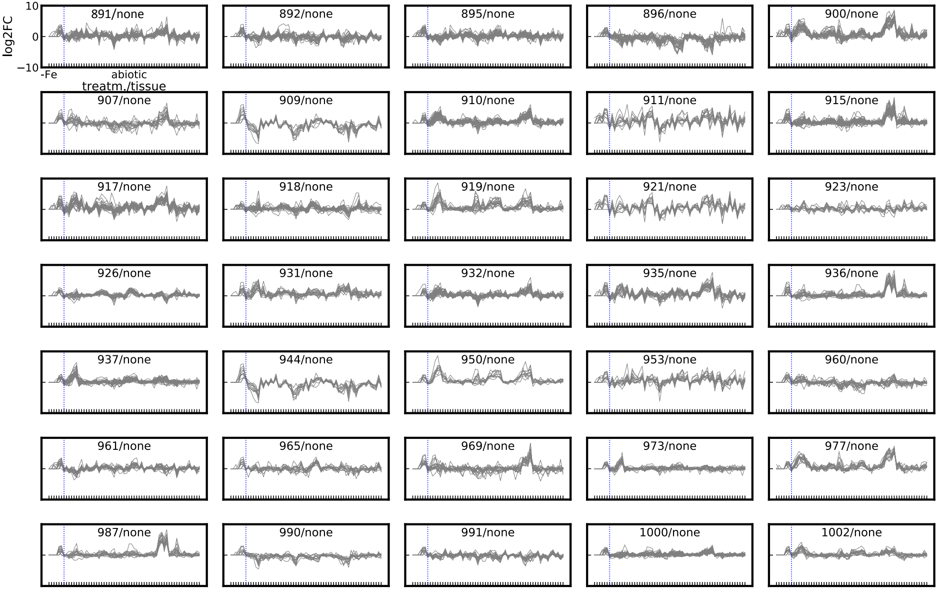

### Fe deficiency time course/abiot. stresses, non-enriched (2)

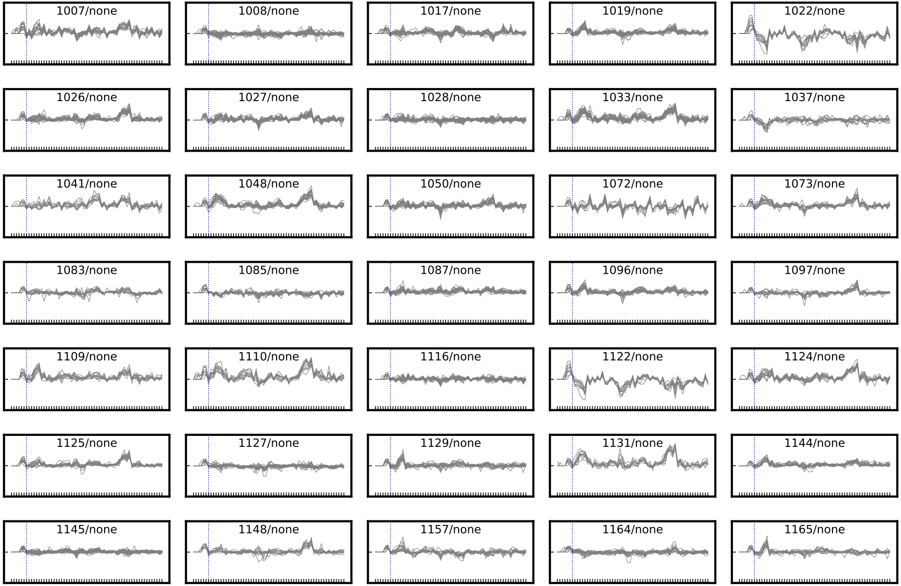

### Fe deficiency time course/abiot. stresses, non-enriched (3)

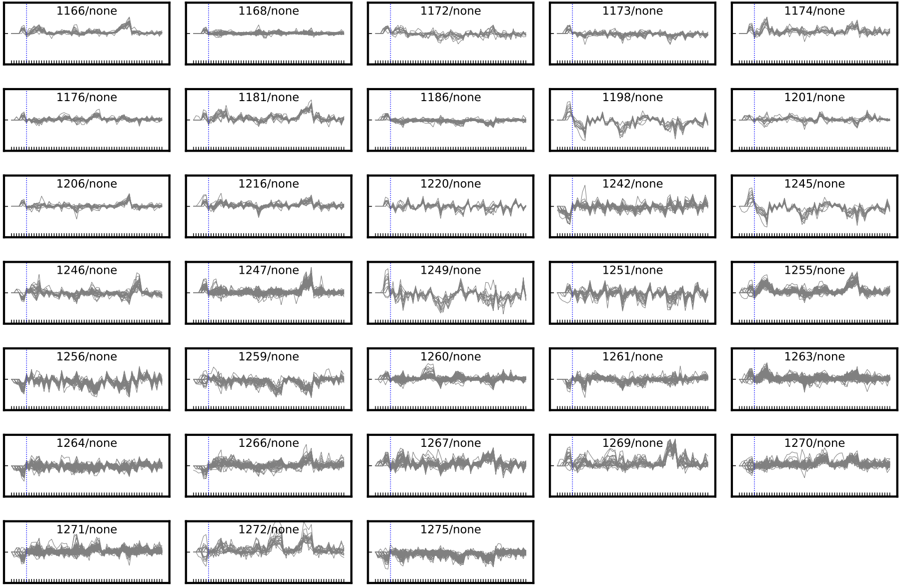

### Fe deficiency time course/abiot. stresses/hormones, GS-enriched

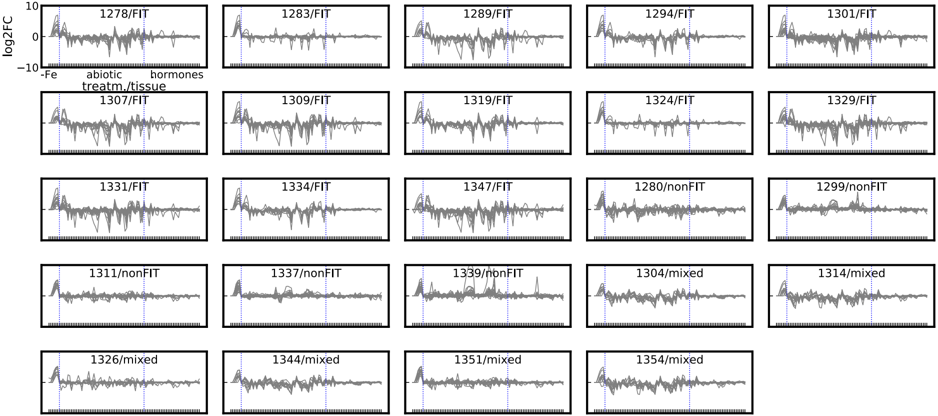

### Fe deficiency time course/abiot. stresses (ng)/hormones, GS-enriched

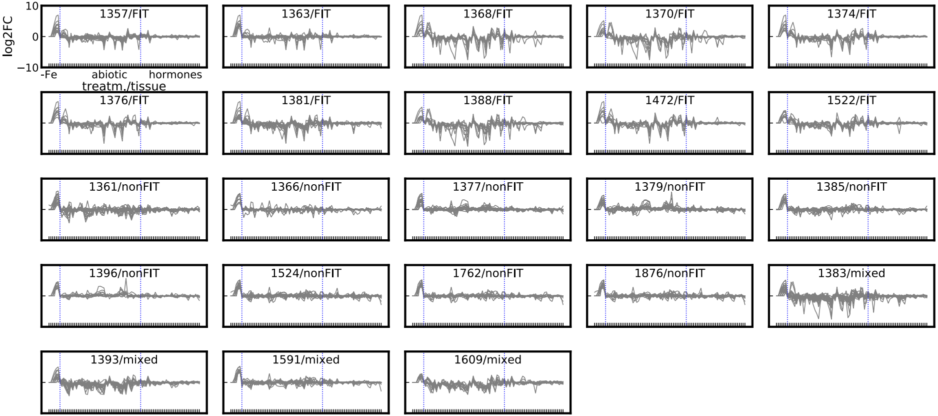

### Fe deficiency time course/abiot. stresses (ng)/hormones, non-enriched (1)

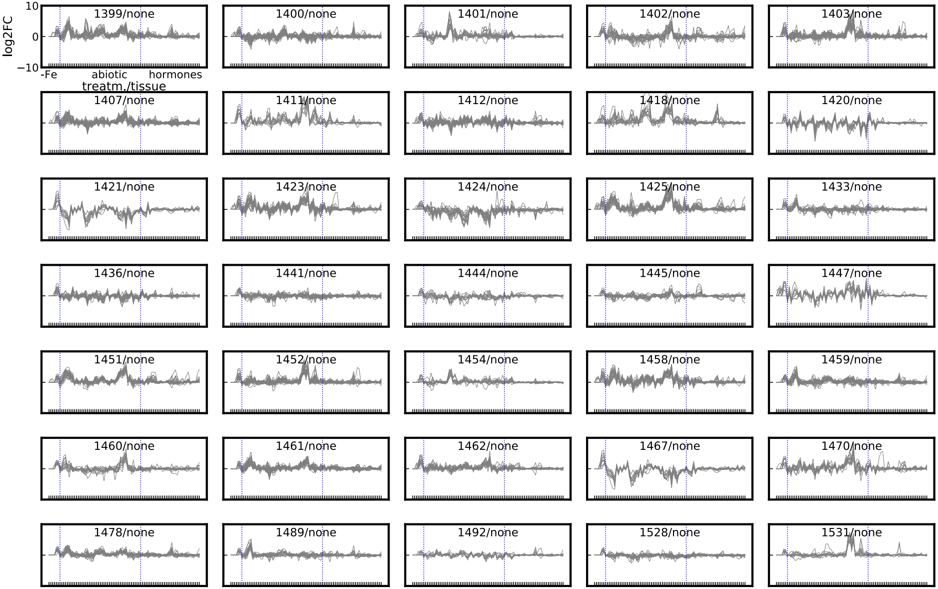

### Fe deficiency time course/abiot. stresses (ng)/hormones, non-enriched (2)

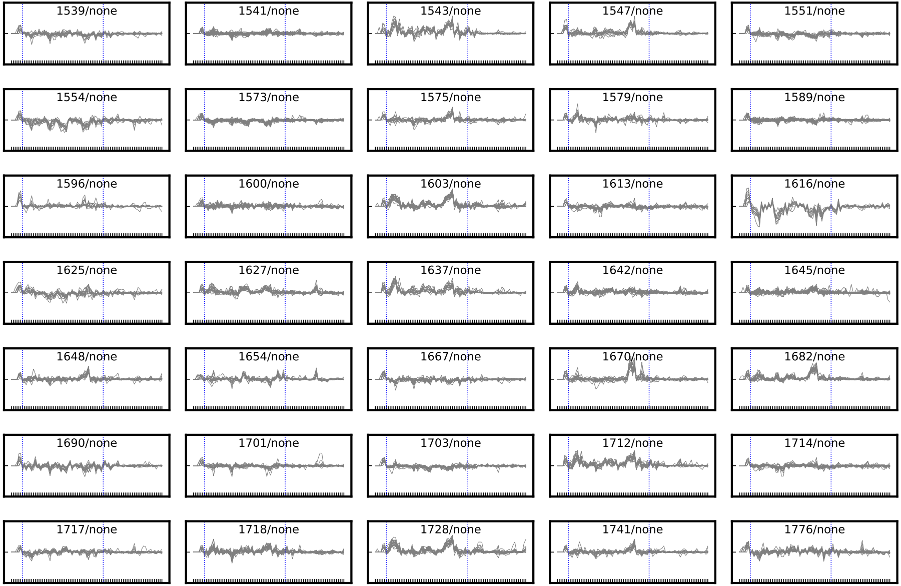

### Fe deficiency time course/abiot. stresses (ng)/hormones, non-enriched (3)

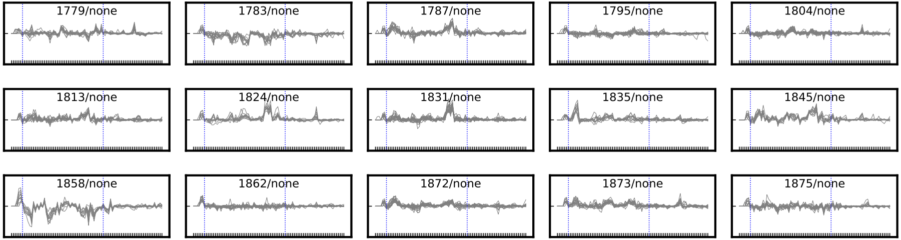

### Fe deficiency time course/abiot. stresses/developm., GS-enriched

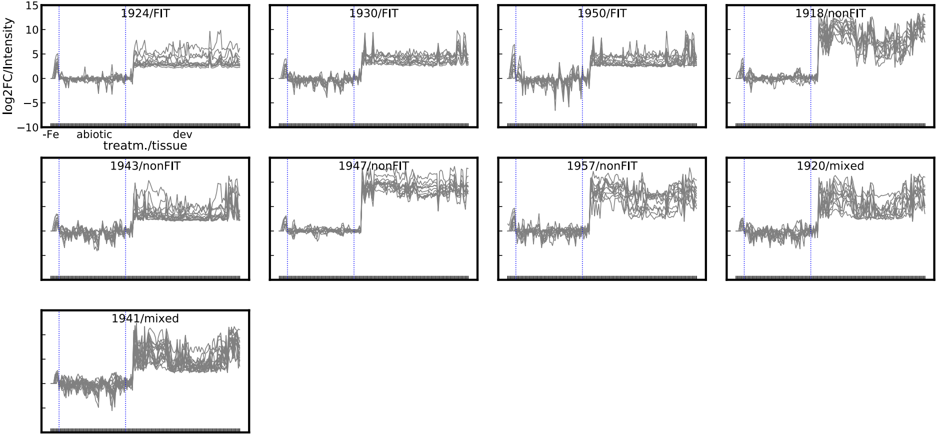

### Fe deficiency time course/abiot. stresses/developm., non-enriched

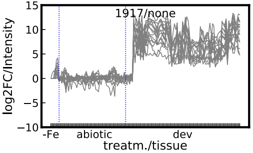


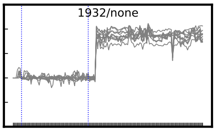
