## Supplemental_Figure_S9 for "Putative *cis*-regulatory elements predict iron deficiency responses in Arabidopsis roots"

**Figure S9.** Positional distribution plots of all 615 min-pCREs.

**Page 1:** Introductory table into min-pCRE ID system (numbered 1-615, sorted by alphabet). **Pages 2-11:** Positional distribution plots. **1st column:** plots of putative promoter region; 1000 bp upstream and 500 bp downstream of TSS (transcription start site). **2nd column:** plots of all introns (In) and all exons (Ex). **3rd column:** plots of putative non-coding region; 500 bp upstream and 1000 bp downstream of TTS (transcription termination site). 1st and 3rd column: position distributions of all co-expression clusters with min-pCRE (gray, filled) plotted as overlay and mean distribution (red, line). 2nd column: bars represent mean with standard deviation. min-pCREs are sorted in columns extended over two pages. **Pages 2+3:** min-pCREs 1-125, **pages 4+5:** min-pCREs 126-250, **pages 6+7:** min-pCREs 251-375, **pages 8+9:** min-pCREs 376-500, **pages 10+11:** min-pCREs 501-615.

Part1

| ID (as in S3/S4/S5_Table) | min-pCRE | Count GS-enriched clusters with min-pCRE | ID in this file |
|---|---|---|---|
| 8 | AAAAACATT | 1 | 1 |
| 46 | AAAAGT | 1 | 2 |
| 50 | AAAAGTC | 2 | 3 |
| 87 | AAACAAC | 2 | 4 |
| 91 | AAACAGC | 1 | 5 |
| 96 | AAACATTA | 1 | 6 |
| 104 | AAACGA | 1 | 7 |
| 112 | AAACGC | 1 | 8 |
| 122 | AAACTA | 7 | 9 |
| 123 | AAACTAC | 1 | 10 |
| 128 | AAACTG | 1 | 11 |
| 150 | AAAGGCT | 1 | 12 |
| 152 | AAAGTA | 9 | 13 |
| 170 | AAAGTCT | 1 | 14 |
| 179 | AAATAAC | 2 | 15 |
| 189 | AAATAG | 3 | 16 |
| 191 | AAATAGT | 1 | 17 |
| 206 | AAATATT | 2 | 18 |
| 209 | AAATATTAT | 1 | 19 |
| 211 | AAATATTT | 2 | 20 |
| 213 | AAATCT | 1 | 21 |
| 215 | AAATCTG | 1 | 22 |
| 230 | AAATGTA | 1 | 23 |
| 235 | AAATTA | 1 | 24 |
| 282 | AACACT | 2 | 25 |
| 286 | AACAGC | 1 | 26 |
| 287 | AACAGG | 1 | 27 |
| 297 | AACATGG | 1 | 28 |
| 298 | AACATT | 1 | 29 |
| 299 | AACATTA | 2 | 30 |
| 310 | AACCCA | 1 | 31 |
| 333 | AACCTC | 1 | 32 |
| 343 | AACGAG | 1 | 33 |
| 352 | AACGCA | 2 | 34 |
| 358 | AACGCC | 2 | 35 |
| 368 | AACGGA | 1 | 36 |
| 370 | AACGGC | 14 | 37 |
| 371 | AACGGCA | 1 | 38 |
| 382 | AACGTA | 15 | 39 |
| 411 | AACTAC | 4 | 40 |
| 412 | AACTACA | 2 | 41 |
| 418 | AACTAG | 1 | 42 |
| 423 | AACTAGT | 1 | 43 |
| 426 | AACTAT | 1 | 44 |
| 427 | AACTATA | 1 | 45 |
| 435 | AACTGA | 1 | 46 |
| 469 | AAGACGA | 2 | 47 |
| 476 | AAGAGT | 1 | 48 |
| 481 | AAGATG | 1 | 49 |
| 483 | AAGCAA | 3 | 50 |
| 484 | AAGCAC | 1 | 51 |
| 501 | AAGCTGC | 2 | 52 |
| 511 | AAGGAG | 1 | 53 |
| 527 | AAGGTC | 1 | 54 |
| 532 | AAGGTTA | 1 | 55 |
| 556 | AAGTCC | 1 | 56 |
| 558 | AAGTCG | 1 | 57 |
| 570 | AAGTGC | 1 | 58 |
| 597 | AATAAT | 3 | 59 |
| 598 | AATAATA | 1 | 60 |
| 599 | AATAATAA | 1 | 61 |
| 624 | AATACTA | 2 | 62 |
| 629 | AATAGA | 4 | 63 |
| 638 | AATAGG | 1 | 64 |
| 640 | AATAGT | 3 | 65 |
| 644 | AATAGTT | 1 | 66 |
| 645 | AATATA | 1 | 67 |
| 655 | AATATAT | 1 | 68 |
| 661 | AATATC | 2 | 69 |
| 665 | AATATG | 8 | 70 |
| 674 | AATATTAG | 1 | 71 |
| 677 | AATATTT | 1 | 72 |
| 680 | AATCAC | 1 | 73 |
| 684 | AATCAT | 3 | 74 |
| 687 | AATCCC | 1 | 75 |
| 688 | AATCCCC | 1 | 76 |
| 696 | AATCTG | 1 | 77 |
| 708 | AATGATA | 1 | 78 |
| 712 | AATGCA | 3 | 79 |
| 721 | AATGCG | 6 | 80 |
| 744 | AATGTG | 2 | 81 |
| 747 | AATTAA | 3 | 82 |
| 756 | AATTAAT | 1 | 83 |
| 760 | AATTAC | 1 | 84 |
| 793 | AATTGC | 13 | 85 |
| 801 | AATTTAA | 1 | 86 |
| 822 | ACAACA | 1 | 87 |
| 823 | ACAACC | 2 | 88 |

Part2

| ID (as in S3/S4/S5_Table) | min-pCRE | Count GS-enriched clusters with min-pCRE | ID in this file |
|---|---|---|---|
| 873 | ACACGA | 1 | 89 |
| 877 | ACACGC | 2 | 90 |
| 885 | ACACGT | 3 | 91 |
| 905 | ACACTA | 9 | 92 |
| 911 | ACACTTA | 1 | 93 |
| 916 | ACAGCA | 5 | 94 |
| 935 | ACATAAC | 3 | 95 |
| 938 | ACATAG | 1 | 96 |
| 940 | ACATAT | 1 | 97 |
| 948 | ACATCA | 1 | 98 |
| 958 | ACATGC | 3 | 99 |
| 963 | ACATGGA | 1 | 100 |
| 967 | ACATTA | 1 | 101 |
| 973 | ACATTC | 1 | 102 |
| 991 | ACCACG | 3 | 103 |
| 996 | ACCAGC | 2 | 104 |
| 999 | ACCATA | 2 | 105 |
| 1001 | ACCATAC | 1 | 106 |
| 1011 | ACCCAC | 14 | 107 |
| 1023 | ACCGAA | 4 | 108 |
| 1045 | ACCGTC | 1 | 109 |
| 1054 | ACCTAC | 3 | 110 |
| 1068 | ACCTCG | 2 | 111 |
| 1069 | ACCTGG | 2 | 112 |
| 1072 | ACCTTA | 3 | 113 |
| 1078 | ACGAAA | 10 | 114 |
| 1080 | ACGAAAAA | 1 | 115 |
| 1081 | ACGAAAT | 2 | 116 |
| 1090 | ACGAAT | 1 | 117 |
| 1113 | ACGAGA | 2 | 118 |
| 1114 | ACGAGG | 2 | 119 |
| 1125 | ACGCAA | 4 | 120 |
| 1126 | ACGCAAA | 3 | 121 |
| 1134 | ACGCAT | 31 | 122 |
| 1141 | ACGCCA | 4 | 123 |
| 1145 | ACGCCG | 1 | 124 |
| 1148 | ACGCGC | 6 | 125 |
| 1151 | ACGCGCC | 2 | 126 |
| 1158 | ACGCGT | 3 | 127 |
| 1166 | ACGCTC | 1 | 128 |
| 1177 | ACGGAG | 2 | 129 |
| 1178 | ACGGCA | 5 | 130 |
| 1191 | ACGGGGT | 1 | 131 |
| 1196 | ACGGTG | 1 | 132 |
| 1197 | ACGTAA | 1 | 133 |
| 1208 | ACGTAG | 1 | 134 |
| 1213 | ACGTAGT | 1 | 135 |
| 1229 | ACGTCA | 1 | 136 |
| 1231 | ACGTCAG | 1 | 137 |
| 1244 | ACGTGA | 1 | 138 |
| 1246 | ACGTGAC | 1 | 139 |
| 1261 | ACGTGG | 4 | 140 |
| 1266 | ACGTGGC | 2 | 141 |
| 1283 | ACGTTA | 2 | 142 |
| 1296 | ACTAAG | 1 | 143 |
| 1306 | ACTACG | 1 | 144 |
| 1310 | ACTACT | 3 | 145 |
| 1316 | ACTAGG | 6 | 146 |
| 1317 | ACTAGGT | 1 | 147 |
| 1318 | ACTAGT | 10 | 148 |
| 1323 | ACTAGTT | 1 | 149 |
| 1324 | ACTATA | 5 | 150 |
| 1325 | ACTATAA | 3 | 151 |
| 1328 | ACTATAT | 1 | 152 |
| 1339 | ACTATTC | 1 | 153 |
| 1346 | ACTCCA | 1 | 154 |
| 1348 | ACTCCG | 2 | 155 |
| 1357 | ACTGAT | 1 | 156 |
| 1365 | ACTGTA | 2 | 157 |
| 1386 | ACTTGTA | 1 | 158 |
| 1400 | AGAAATA | 1 | 159 |
| 1406 | AGAATA | 2 | 160 |
| 1413 | AGAATG | 1 | 161 |
| 1424 | AGACCTT | 1 | 162 |
| 1426 | AGACGA | 2 | 163 |
| 1431 | AGACGC | 2 | 164 |
| 1438 | AGACTG | 2 | 165 |
| 1450 | AGAGGC | 1 | 166 |
| 1453 | AGAGTGA | 1 | 167 |
| 1468 | AGATTA | 3 | 168 |
| 1469 | AGATTAA | 1 | 169 |
| 1472 | AGCAAT | 2 | 170 |
| 1474 | AGCACA | 1 | 171 |
| 1481 | AGCATA | 1 | 172 |
| 1483 | AGCATG | 10 | 173 |
| 1485 | AGCATGT | 1 | 174 |
| 1489 | AGCCAAA | 3 | 175 |
| 1499 | AGCCGG | 2 | 176 |

Part3

| ID (as in S3/S4/S5_Table) | min-pCRE | Count GS-enriched clusters with min-pCRE | ID in this file |
|---|---|---|---|
| 1500 | AGCCGGA | 3 | 177 |
| 1510 | AGCGCC | 1 | 178 |
| 1555 | AGGCAG | 1 | 179 |
| 1558 | AGGCAT | 6 | 180 |
| 1575 | AGGGAAA | 1 | 181 |
| 1576 | AGGGAC | 3 | 182 |
| 1594 | AGGTCA | 1 | 183 |
| 1600 | AGGTTA | 4 | 184 |
| 1604 | AGTAAC | 6 | 185 |
| 1619 | AGTAGG | 4 | 186 |
| 1625 | AGTATA | 5 | 187 |
| 1641 | AGTCAA | 8 | 188 |
| 1647 | AGTCAAC | 5 | 189 |
| 1667 | AGTCAT | 4 | 190 |
| 1682 | AGTCGG | 1 | 191 |
| 1686 | AGTCTA | 1 | 192 |
| 1695 | AGTGAAT | 1 | 193 |
| 1696 | AGTGAC | 1 | 194 |
| 1701 | AGTGAT | 2 | 195 |
| 1704 | AGTGGC | 7 | 196 |
| 1705 | AGTGGCA | 1 | 197 |
| 1711 | AGTGGG | 4 | 198 |
| 1713 | AGTGGGG | 3 | 199 |
| 1715 | AGTGTA | 1 | 200 |
| 1724 | AGTTAAA | 2 | 201 |
| 1728 | AGTTAT | 1 | 202 |
| 1730 | AGTTATAT | 1 | 203 |
| 1734 | AGTTATT | 1 | 204 |
| 1769 | ATAAATA | 2 | 205 |
| 1770 | ATAAATAA | 1 | 206 |
| 1775 | ATAACA | 1 | 207 |
| 1779 | ATAACC | 6 | 208 |
| 1784 | ATAACG | 1 | 209 |
| 1794 | ATAAGC | 4 | 210 |
| 1801 | ATAATA | 2 | 211 |
| 1807 | ATAATAC | 2 | 212 |
| 1820 | ATAATCA | 3 | 213 |
| 1822 | ATAATCC | 1 | 214 |
| 1824 | ATAATG | 2 | 215 |
| 1835 | ATAATTA | 1 | 216 |
| 1841 | ATACAAT | 1 | 217 |
| 1842 | ATACAC | 2 | 218 |
| 1850 | ATACAT | 2 | 219 |
| 1860 | ATACCC | 3 | 220 |
| 1883 | ATACTA | 2 | 221 |
| 1893 | ATACTGA | 1 | 222 |
| 1899 | ATAGAATA | 1 | 223 |
| 1900 | ATAGAC | 1 | 224 |
| 1913 | ATAGGC | 1 | 225 |
| 1939 | ATATAC | 2 | 226 |
| 1940 | ATATACA | 2 | 227 |
| 1948 | ATATAG | 1 | 228 |
| 1956 | ATATAT | 6 | 229 |
| 1967 | ATATATT | 5 | 230 |
| 1971 | ATATCA | 2 | 231 |
| 1974 | ATATCG | 3 | 232 |
| 1979 | ATATGA | 1 | 233 |
| 1990 | ATATTAA | 1 | 234 |
| 1995 | ATATTAT | 1 | 235 |
| 2021 | ATCACG | 1 | 236 |
| 2035 | ATCCCG | 1 | 237 |
| 2037 | ATCCTA | 2 | 238 |
| 2042 | ATCGCA | 4 | 239 |
| 2043 | ATCGCC | 1 | 240 |
| 2055 | ATCGTG | 10 | 241 |
| 2064 | ATCTTC | 1 | 242 |
| 2089 | ATGAGC | 2 | 243 |
| 2091 | ATGAGCT | 1 | 244 |
| 2100 | ATGCAA | 2 | 245 |
| 2101 | ATGCAC | 1 | 246 |
| 2105 | ATGCAT | 13 | 247 |
| 2110 | ATGCATTA | 3 | 248 |
| 2111 | ATGCCA | 2 | 249 |
| 2114 | ATGCCC | 1 | 250 |
| 2117 | ATGCGA | 2 | 251 |
| 2128 | ATGCTG | 1 | 252 |
| 2142 | ATGGCG | 3 | 253 |
| 2149 | ATGGTA | 1 | 254 |
| 2152 | ATGTAA | 5 | 255 |
| 2168 | ATGTGG | 1 | 256 |
| 2187 | ATTAAC | 2 | 257 |
| 2192 | ATTAAG | 15 | 258 |
| 2206 | ATTACA | 3 | 259 |
| 2213 | ATTACG | 2 | 260 |
| 2219 | ATTAGA | 4 | 261 |
| 2221 | ATTAGC | 3 | 262 |
| 2222 | ATTAGCT | 1 | 263 |
| 2226 | ATTATA | 6 | 264 |

Part4

| ID (as in S3/S4/S5_Table) | min-pCRE | Count GS-enriched clusters with min-pCRE | ID in this file |
|---|---|---|---|
| 2236 | ATTATC | 1 | 265 |
| 2238 | ATTATGA | 1 | 266 |
| 2239 | ATTCAA | 1 | 267 |
| 2247 | ATTCCA | 1 | 268 |
| 2248 | ATTCCAC | 2 | 269 |
| 2253 | ATTCGG | 5 | 270 |
| 2266 | ATTGAA | 1 | 271 |
| 2278 | ATTGAG | 3 | 272 |
| 2280 | ATTGCC | 1 | 273 |
| 2284 | ATTGGAT | 2 | 274 |
| 2285 | ATTGTA | 1 | 275 |
| 2286 | ATTGTAT | 1 | 276 |
| 2298 | ATTTAC | 4 | 277 |
| 2301 | ATTTAGA | 4 | 278 |
| 2349 | CAAATAG | 1 | 279 |
| 2365 | CAACCAG | 2 | 280 |
| 2374 | CAACGC | 3 | 281 |
| 2387 | CAACTA | 1 | 282 |
| 2409 | CAAGCG | 1 | 283 |
| 2413 | CAAGGG | 1 | 284 |
| 2429 | CAATAA | 1 | 285 |
| 2433 | CAATAC | 2 | 286 |
| 2445 | CAATTA | 2 | 287 |
| 2446 | CAATTAA | 3 | 288 |
| 2473 | CACACG | 2 | 289 |
| 2477 | CACACGC | 2 | 290 |
| 2479 | CACACGT | 2 | 291 |
| 2506 | CACCACG | 1 | 292 |
| 2530 | CACGAG | 8 | 293 |
| 2534 | CACGCA | 8 | 294 |
| 2541 | CACGCC | 15 | 295 |
| 2542 | CACGCCA | 1 | 296 |
| 2547 | CACGCG | 1 | 297 |
| 2559 | CACGTA | 1 | 298 |
| 2572 | CACGTC | 6 | 299 |
| 2573 | CACGTCA | 1 | 300 |
| 2576 | CACGTCAG | 1 | 301 |
| 2584 | CACGTG | 3 | 302 |
| 2609 | CACTAC | 10 | 303 |
| 2612 | CACTACT | 1 | 304 |
| 2616 | CACTCG | 1 | 305 |
| 2620 | CACTTA | 1 | 306 |
| 2630 | CAGACAA | 1 | 307 |
| 2637 | CAGCAC | 1 | 308 |
| 2656 | CAGGTC | 1 | 309 |
| 2666 | CAGTCG | 1 | 310 |
| 2676 | CAGTTTA | 1 | 311 |
| 2679 | CATAACA | 1 | 312 |
| 2685 | CATACA | 2 | 313 |
| 2692 | CATACG | 2 | 314 |
| 2741 | CATGCA | 10 | 315 |
| 2748 | CATGCC | 48 | 316 |
| 2751 | CATGCG | 3 | 317 |
| 2765 | CATGTA | 1 | 318 |
| 2774 | CATTAC | 1 | 319 |
| 2785 | CATTCC | 3 | 320 |
| 2792 | CATTGG | 1 | 321 |
| 2794 | CATTTA | 1 | 322 |
| 2797 | CATTTAG | 1 | 323 |
| 2824 | CCAAGG | 3 | 324 |
| 2832 | CCACAA | 1 | 325 |
| 2843 | CCACGA | 2 | 326 |
| 2846 | CCACGC | 4 | 327 |
| 2855 | CCACGG | 1 | 328 |
| 2858 | CCACGTC | 3 | 329 |
| 2864 | CCACTA | 3 | 330 |
| 2865 | CCACTC | 2 | 331 |
| 2868 | CCAGCAC | 1 | 332 |
| 2869 | CCAGCC | 1 | 333 |
| 2870 | CCAGCG | 1 | 334 |
| 2879 | CCATAC | 6 | 335 |
| 2881 | CCATACT | 1 | 336 |
| 2887 | CCATGC | 19 | 337 |
| 2895 | CCATTA | 1 | 338 |
| 2897 | CCATTC | 4 | 339 |
| 2907 | CCCACG | 9 | 340 |
| 2911 | CCCACTC | 1 | 341 |
| 2920 | CCCCAC | 4 | 342 |
| 2923 | CCCCACT | 4 | 343 |
| 2951 | CCCTCG | 1 | 344 |
| 2960 | CCGAGG | 2 | 345 |
| 2972 | CCGCGG | 1 | 346 |
| 2980 | CCGGCC | 7 | 347 |
| 2981 | CCGGCCG | 1 | 348 |
| 2992 | CCGTCA | 1 | 349 |
| 3006 | CCTACG | 5 | 350 |
| 3038 | CCTTGA | 1 | 351 |
| 3042 | CGAAAA | 1 | 352 |

Part5

| ID (as in S3/S4/S5_Table) | min-pCRE | Count GS-enriched clusters with min-pCRE | ID in this file |
|---|---|---|---|
| 3048 | CGAAGA | 1 | 353 |
| 3049 | CGAAGC | 1 | 354 |
| 3050 | CGAATA | 2 | 355 |
| 3055 | CGACAC | 3 | 356 |
| 3059 | CGACCA | 1 | 357 |
| 3079 | CGAGGC | 3 | 358 |
| 3085 | CGATGA | 2 | 359 |
| 3086 | CGATTA | 2 | 360 |
| 3104 | CGCATA | 1 | 361 |
| 3105 | CGCATC | 2 | 362 |
| 3108 | CGCCAC | 1 | 363 |
| 3120 | CGCGAC | 1 | 364 |
| 3123 | CGCGCC | 1 | 365 |
| 3140 | CGCTCA | 12 | 366 |
| 3150 | CGCTTA | 5 | 367 |
| 3155 | CGGACC | 1 | 368 |
| 3164 | CGGCCA | 8 | 369 |
| 3166 | CGGCCG | 1 | 370 |
| 3177 | CGGTAC | 1 | 371 |
| 3190 | CGTAAA | 3 | 372 |
| 3194 | CGTAAC | 5 | 373 |
| 3197 | CGTAAG | 6 | 374 |
| 3200 | CGTACA | 1 | 375 |
| 3212 | CGTATAA | 1 | 376 |
| 3217 | CGTATC | 1 | 377 |
| 3223 | CGTCAG | 3 | 378 |
| 3225 | CGTCAGC | 1 | 379 |
| 3235 | CGTCGA | 4 | 380 |
| 3238 | CGTCGAC | 2 | 381 |
| 3244 | CGTGAC | 21 | 382 |
| 3247 | CGTGAG | 1 | 383 |
| 3255 | CGTGCC | 2 | 384 |
| 3262 | CGTGGA | 16 | 385 |
| 3263 | CGTGGAA | 1 | 386 |
| 3267 | CGTGGC | 2 | 387 |
| 3280 | CGTGTC | 3 | 388 |
| 3291 | CGTTAG | 1 | 389 |
| 3299 | CGTTGA | 3 | 390 |
| 3300 | CGTTGAA | 4 | 391 |
| 3303 | CGTTGGA | 1 | 392 |
| 3321 | CTAATA | 3 | 393 |
| 3329 | CTAATTA | 1 | 394 |
| 3333 | CTACAC | 3 | 395 |
| 3354 | CTAGGC | 1 | 396 |
| 3355 | CTAGTA | 4 | 397 |
| 3367 | CTATAC | 1 | 398 |
| 3388 | CTATTCG | 1 | 399 |
| 3406 | CTCAGC | 2 | 400 |
| 3419 | CTCCTCT | 1 | 401 |
| 3426 | CTCTAAA | 1 | 402 |
| 3429 | CTCTACT | 1 | 403 |
| 3442 | CTGAGA | 1 | 404 |
| 3443 | CTGATA | 2 | 405 |
| 3455 | CTGGAA | 1 | 406 |
| 3461 | CTGTAA | 2 | 407 |
| 3477 | CTTATA | 6 | 408 |
| 3493 | CTTGGC | 1 | 409 |
| 3494 | CTTGTA | 1 | 410 |
| 3495 | CTTGTAT | 2 | 411 |
| 3497 | CTTGTC | 1 | 412 |
| 3500 | CTTTAAT | 1 | 413 |
| 3546 | GAACGCA | 4 | 414 |
| 3568 | GAAGGTA | 1 | 415 |
| 3574 | GAAGTGA | 1 | 416 |
| 3576 | GAATAA | 2 | 417 |
| 3601 | GAATTA | 3 | 418 |
| 3611 | GACAAA | 3 | 419 |
| 3623 | GACAGC | 5 | 420 |
| 3639 | GACCAC | 5 | 421 |
| 3645 | GACCGC | 1 | 422 |
| 3656 | GACGAA | 1 | 423 |
| 3674 | GACGTA | 1 | 424 |
| 3680 | GACGTGG | 2 | 425 |
| 3701 | GACTCC | 4 | 426 |
| 3723 | GAGCCCA | 1 | 427 |
| 3733 | GAGGGA | 7 | 428 |
| 3741 | GAGTAC | 1 | 429 |
| 3742 | GAGTCA | 8 | 430 |
| 3763 | GATAAC | 6 | 431 |
| 3773 | GATAGA | 4 | 432 |
| 3789 | GATCTA | 1 | 433 |
| 3802 | GATGGC | 1 | 434 |
| 3807 | GATTAA | 1 | 435 |
| 3808 | GATTAAG | 1 | 436 |
| 3810 | GATTACA | 2 | 437 |
| 3818 | GATTTA | 1 | 438 |
| 3819 | GATTTAA | 1 | 439 |
| 3821 | GCAAAC | 1 | 440 |

Part6

| ID (as in S3/S4/S5_Table) | min-pCRE | Count GS-enriched clusters with min-pCRE | ID in this file |
|---|---|---|---|
| 3827 | GCACAA | 1 | 441 |
| 3838 | GCACGC | 2 | 442 |
| 3854 | GCAGCA | 1 | 443 |
| 3875 | GCATGA | 20 | 444 |
| 3876 | GCATGAA | 1 | 445 |
| 3885 | GCATGC | 3 | 446 |
| 3889 | GCATTA | 4 | 447 |
| 3893 | GCATTAT | 3 | 448 |
| 3917 | GCCCAC | 3 | 449 |
| 3918 | GCCCATA | 1 | 450 |
| 3919 | GCCCCA | 5 | 451 |
| 3941 | GCCTAC | 2 | 452 |
| 3955 | GCGCAC | 1 | 453 |
| 3961 | GCGCTGA | 1 | 454 |
| 3968 | GCGTAA | 1 | 455 |
| 3977 | GCGTCA | 1 | 456 |
| 3980 | GCGTCC | 1 | 457 |
| 3981 | GCGTGA | 6 | 458 |
| 4001 | GCTATA | 1 | 459 |
| 4002 | GCTATAA | 1 | 460 |
| 4017 | GCTGAC | 5 | 461 |
| 4050 | GGACCA | 1 | 462 |
| 4053 | GGACCC | 1 | 463 |
| 4060 | GGAGCC | 11 | 464 |
| 4062 | GGAGCCCA | 3 | 465 |
| 4083 | GGATTA | 1 | 466 |
| 4087 | GGCAAAA | 1 | 467 |
| 4096 | GGCAGA | 1 | 468 |
| 4107 | GGCCAC | 2 | 469 |
| 4115 | GGCCCC | 2 | 470 |
| 4130 | GGCGTA | 1 | 471 |
| 4134 | GGCGTGA | 1 | 472 |
| 4140 | GGGACC | 1 | 473 |
| 4143 | GGGATA | 1 | 474 |
| 4148 | GGGCCA | 1 | 475 |
| 4158 | GGGGCA | 2 | 476 |
| 4172 | GGGTCA | 3 | 477 |
| 4175 | GGGTTA | 1 | 478 |
| 4179 | GGTAAA | 2 | 479 |
| 4196 | GGTCAC | 3 | 480 |
| 4201 | GGTCGA | 3 | 481 |
| 4207 | GGTGAC | 5 | 482 |
| 4210 | GGTGTAA | 1 | 483 |
| 4211 | GGTTAA | 1 | 484 |
| 4216 | GGTTCA | 1 | 485 |
| 4226 | GTAAAC | 1 | 486 |
| 4231 | GTAATA | 1 | 487 |
| 4235 | GTAATCA | 1 | 488 |
| 4237 | GTACAA | 4 | 489 |
| 4242 | GTACCA | 1 | 490 |
| 4252 | GTAGAA | 2 | 491 |
| 4253 | GTAGAAA | 1 | 492 |
| 4262 | GTATAA | 10 | 493 |
| 4276 | GTATTA | 13 | 494 |
| 4278 | GTATTAAA | 3 | 495 |
| 4282 | GTATTAT | 2 | 496 |
| 4285 | GTCAAA | 2 | 497 |
| 4292 | GTCAAC | 15 | 498 |
| 4310 | GTCAGA | 5 | 499 |
| 4311 | GTCATA | 1 | 500 |
| 4318 | GTCCAC | 1 | 501 |
| 4330 | GTCGGA | 8 | 502 |
| 4337 | GTCTTA | 4 | 503 |
| 4343 | GTCTTAT | 1 | 504 |
| 4349 | GTGCAA | 5 | 505 |
| 4354 | GTGCCA | 5 | 506 |
| 4364 | GTGGAA | 2 | 507 |
| 4367 | GTGGAAT | 1 | 508 |
| 4370 | GTGGCA | 8 | 509 |
| 4397 | GTGTCAA | 1 | 510 |
| 4401 | GTGTTA | 4 | 511 |
| 4403 | GTGTTAT | 3 | 512 |
| 4404 | GTTAAA | 2 | 513 |
| 4410 | GTTATA | 9 | 514 |
| 4420 | GTTCTA | 3 | 515 |
| 4426 | GTTGCA | 3 | 516 |
| 4427 | GTTGGA | 1 | 517 |
| 4457 | TAAACTA | 1 | 518 |
| 4462 | TAAAGA | 2 | 519 |
| 4463 | TAAAGTA | 3 | 520 |
| 4467 | TAAATA | 3 | 521 |
| 4468 | TAAATAA | 1 | 522 |
| 4472 | TAAATAG | 2 | 523 |
| 4474 | TAAATTAA | 2 | 524 |
| 4499 | TAACTA | 18 | 525 |
| 4506 | TAAGAA | 1 | 526 |
| 4508 | TAAGCA | 4 | 527 |
| 4509 | TAAGCAC | 1 | 528 |

Part7

| ID (as in S3/S4/S5_Table) | min-pCRE | Count GS-enriched clusters with min-pCRE | ID in this file |
|---|---|---|---|
| 4526 | TAAGTATA | 1 | 529 |
| 4535 | TAATAAT | 1 | 530 |
| 4553 | TAATATG | 2 | 531 |
| 4563 | TAATGA | 1 | 532 |
| 4580 | TAATTA | 8 | 533 |
| 4581 | TAATTAA | 13 | 534 |
| 4583 | TAATTAAC | 1 | 535 |
| 4588 | TAATTAT | 2 | 536 |
| 4593 | TAATTCC | 1 | 537 |
| 4595 | TAATTGC | 1 | 538 |
| 4606 | TACAATA | 1 | 539 |
| 4621 | TACAGA | 5 | 540 |
| 4622 | TACAGAA | 1 | 541 |
| 4640 | TACCATAC | 1 | 542 |
| 4648 | TACCGA | 3 | 543 |
| 4654 | TACCTA | 1 | 544 |
| 4659 | TACGAA | 6 | 545 |
| 4660 | TACGAAA | 1 | 546 |
| 4663 | TACGAAT | 1 | 547 |
| 4685 | TACGTA | 1 | 548 |
| 4707 | TACTAA | 6 | 549 |
| 4715 | TACTACT | 1 | 550 |
| 4726 | TACTCA | 1 | 551 |
| 4733 | TAGAATA | 1 | 552 |
| 4741 | TAGATA | 1 | 553 |
| 4797 | TAGTGGC | 1 | 554 |
| 4803 | TAGTTAC | 1 | 555 |
| 4805 | TAGTTTA | 1 | 556 |
| 4816 | TATAACC | 4 | 557 |
| 4821 | TATAATA | 1 | 558 |
| 4830 | TATACAA | 2 | 559 |
| 4835 | TATACAT | 1 | 560 |
| 4844 | TATAGA | 8 | 561 |
| 4860 | TATATA | 1 | 562 |
| 4866 | TATATACA | 2 | 563 |
| 4874 | TATATAT | 1 | 564 |
| 4904 | TATGAA | 1 | 565 |
| 4910 | TATGATA | 1 | 566 |
| 4911 | TATGCA | 8 | 567 |
| 4912 | TATGCAA | 6 | 568 |
| 4923 | TATGTTA | 1 | 569 |
| 4924 | TATTAA | 3 | 570 |
| 4939 | TATTCA | 1 | 571 |
| 4947 | TATTGA | 1 | 572 |
| 4959 | TATTTAC | 1 | 573 |
| 4979 | TCAACTA | 2 | 574 |
| 5010 | TCACGCC | 1 | 575 |
| 5032 | TCAGCGC | 2 | 576 |
| 5036 | TCATAA | 1 | 577 |
| 5061 | TCATTCC | 1 | 578 |
| 5062 | TCATTCCA | 2 | 579 |
| 5064 | TCATTGG | 1 | 580 |
| 5065 | TCATTTA | 1 | 581 |
| 5088 | TCCCCA | 3 | 582 |
| 5128 | TCGGAA | 2 | 583 |
| 5142 | TCGTAA | 1 | 584 |
| 5161 | TCGTTCC | 1 | 585 |
| 5165 | TCTAAAT | 1 | 586 |
| 5184 | TCTCAA | 3 | 587 |
| 5194 | TCTTAA | 2 | 588 |
| 5210 | TGAAGCA | 1 | 589 |
| 5218 | TGACAA | 3 | 590 |
| 5225 | TGACAGC | 5 | 591 |
| 5259 | TGATAA | 1 | 592 |
| 5269 | TGATTTAA | 1 | 593 |
| 5270 | TGCACA | 2 | 594 |
| 5279 | TGCATGA | 1 | 595 |
| 5281 | TGCATGC | 2 | 596 |
| 5331 | TGGACA | 2 | 597 |
| 5353 | TGGGAA | 1 | 598 |
| 5363 | TGGTAA | 3 | 599 |
| 5373 | TGTACA | 3 | 600 |
| 5374 | TGTACAA | 2 | 601 |
| 5377 | TGTATAA | 1 | 602 |
| 5380 | TGTATCA | 2 | 603 |
| 5418 | TGTTAA | 2 | 604 |
| 5422 | TGTTAAT | 3 | 605 |
| 5434 | TTAACTA | 1 | 606 |
| 5441 | TTAATAAA | 3 | 607 |
| 5444 | TTAATTA | 3 | 608 |
| 5451 | TTAATTTA | 1 | 609 |
| 5484 | TTAGCTA | 1 | 610 |
| 5494 | TTATAA | 3 | 611 |
| 5502 | TTATATA | 1 | 612 |
| 5550 | TTCGTAA | 1 | 613 |
| 5583 | TTGTACA | 1 | 614 |
| 5619 | TTTATTAA | 1 | 615 |

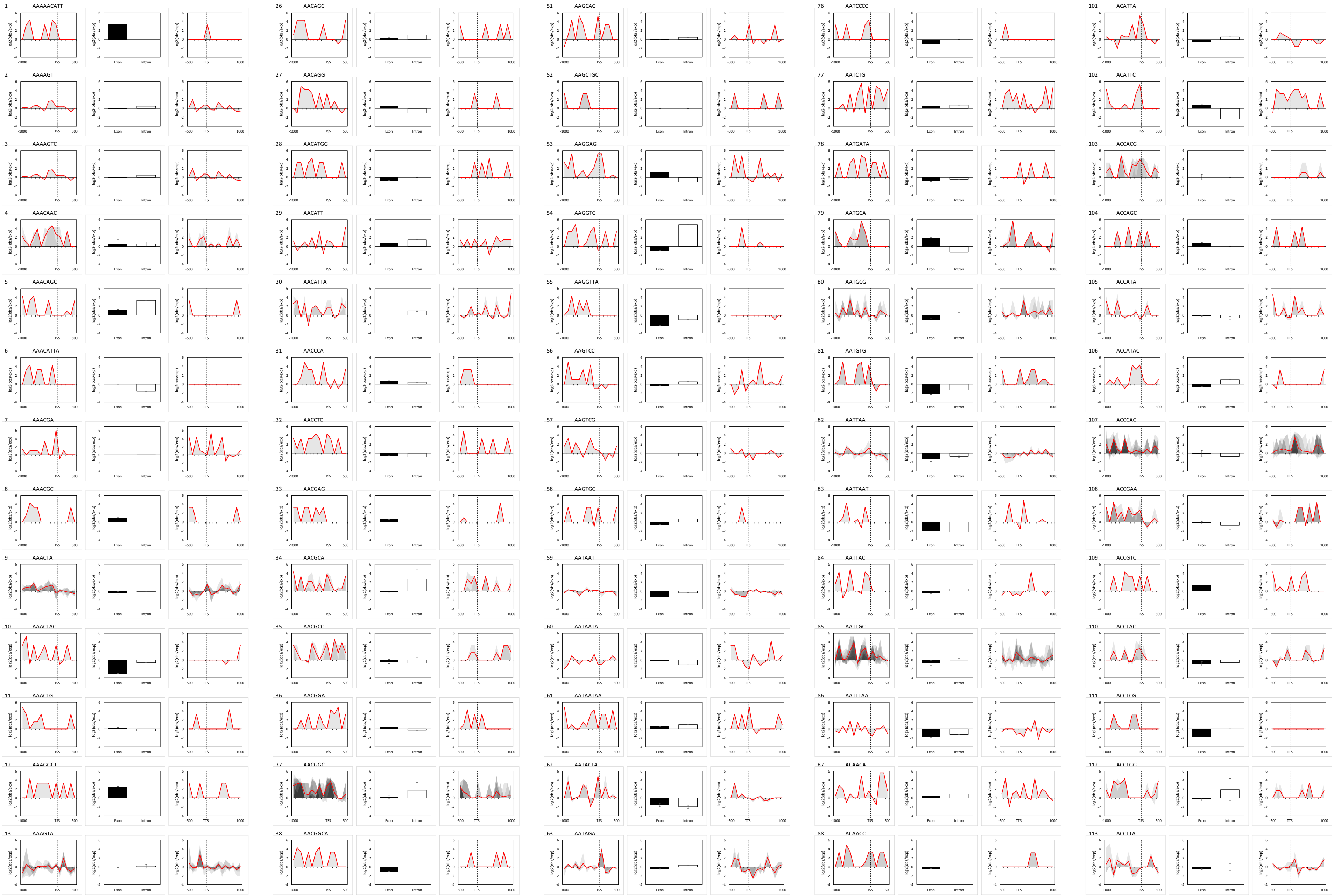
1 AAAAACATT
2 AAAAGT
3 AAAAGTC
4 AAACAAC
5 AAACAGC
6 AAACATTA
7 AAACGA
8 AAACGC
9 AAACTA
10 AAACTAC
11 AAACTG
12 AAAGGCT
13 AAAGTA
26 AACAGC
27 AACAGG
28 AACATGG
29 AACATT
30 AACATTA
31 AACCCA
32 AACCTC
33 AACGAG
34 AACGCA
35 AACGCC
36 AACGGA
37 AACGGC
38 AACGGCA
51 AAGCAC
52 AAGCTGC
53 AAGGAG
54 AAGGTC
55 AAGGTTA
56 AAGTCC
57 AAGTCG
58 AAGTGC
59 AATAAT
60 AATAATA
61 AATAATAA
62 AATACTA
63 AATAGA
76 AATCCCC
77 AATCTG
78 AATGATA
79 AATGCA
80 AATGCG
81 AATGTG
82 AATTAA
83 AATTAAT
84 AATTAC
85 AATTGC
86 AATTTAA
87 ACAACA
88 ACAACC
101 ACATTA
102 ACATTC
103 ACCACG
104 ACCAGC
105 ACCATA
106 ACCATAC
107 ACCCAC
108 ACCGAA
109 ACCGTC
110 ACCTAC
111 ACCTCG
112 ACCTGG
113 ACCTTA

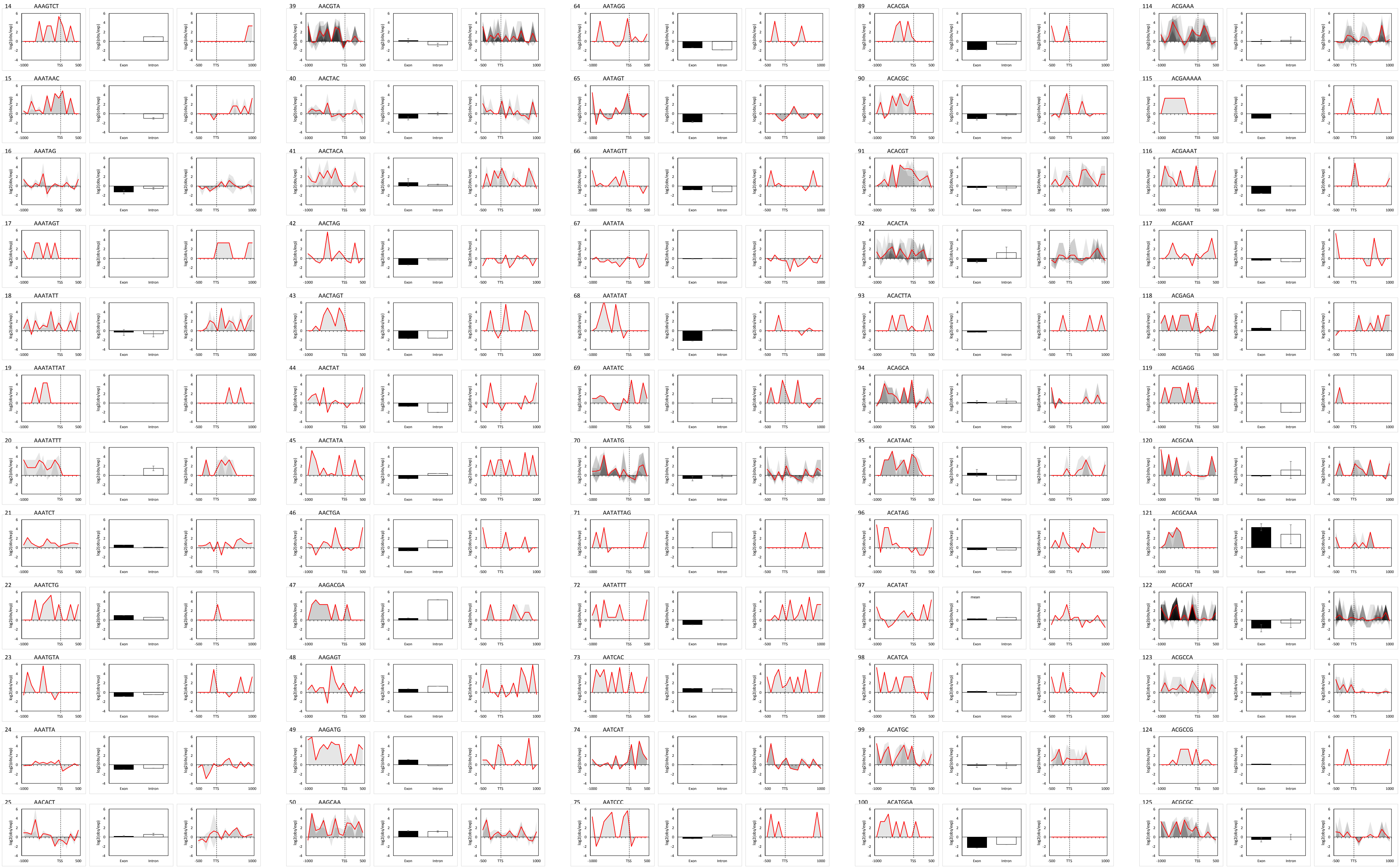

14 AAAGTCT
15 AAATAAC
16 AAATAG
17 AAATAGT
18 AAATATT
19 AAATATTAT
20 AAATATTT
21 AAATCT
22 AAATCTG
23 AAATGTA
24 AAATTA
25 AACACT
39 AACGTA
40 AACTAC
41 AACTACA
42 AACTAG
43 AACTAGT
44 AACTAT
45 AACTATA
46 AACTGA
47 AAGACGA
48 AAGAGT
49 AAGATG
50 AAGCAA
64 AATAGG
65 AATAGT
66 AATAGTT
67 AATATA
68 AATATAT
69 AATATC
70 AATATG
71 AATATTAG
72 AATATTT
73 AATCAC
74 AATCAT
75 AATCCC
89 ACACGA
90 ACACGC
91 ACACGT
92 ACACTA
93 ACACTTA
94 ACAGCA
95 ACATAAC
96 ACATAG
97 ACATAT
98 ACATCA
99 ACATGC
100 ACATGGA
114 ACGAAA
115 ACGAAAAA
116 ACGAAAT
117 ACGAAT
118 ACGAGA
119 ACGAGG
120 ACGCAA
121 ACGCAAA
122 ACGCAT
123 ACGCCA
124 ACGCCG
125 ACGCGC
log2(obs/exp)
TSS
TTS
Exon
Intron

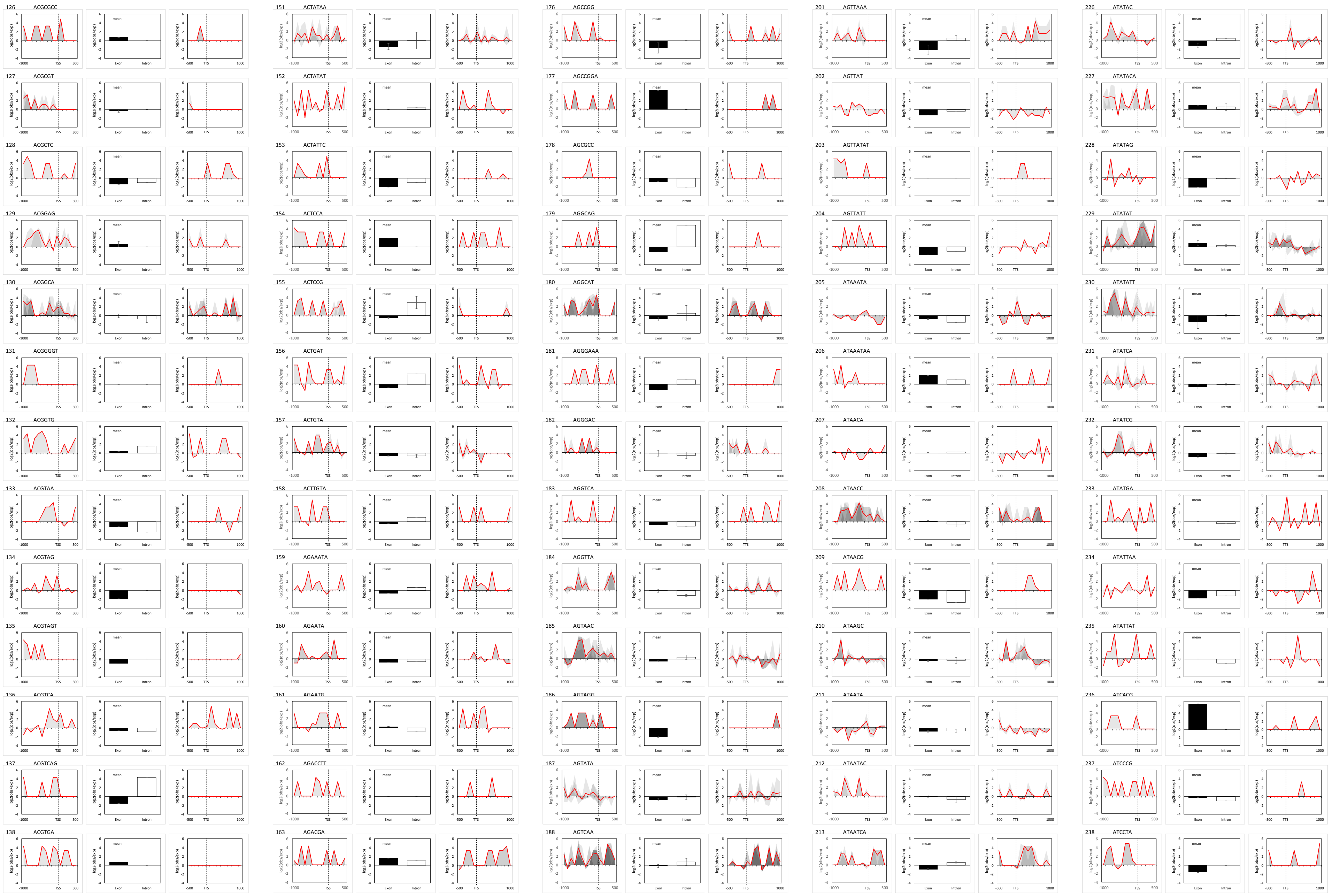
126 ACGCGCC
127 ACGCGT
128 ACGCTC
129 ACGGAG
130 ACGGCA
131 ACGGGGT
132 ACGGTG
133 ACGTAA
134 ACGTAG
135 ACGTAGT
136 ACGTCA
137 ACGTCAG
138 ACGTGA
151 ACTATAA
152 ACTATAT
153 ACTATTC
154 ACTCCA
155 ACTCCG
156 ACTGAT
157 ACTGTA
158 ACTTGTA
159 AGAAATA
160 AGAATA
161 AGAATG
162 AGACCTT
163 AGACGA
176 AGCCGG
177 AGCCGGA
178 AGCGCC
179 AGGCAG
180 AGGCAT
181 AGGGAAA
182 AGGGAC
183 AGGTCA
184 AGGTTA
185 AGTAAC
186 AGTAGG
187 AGTATA
188 AGTCAA
201 AGTTAAA
202 AGTTAT
203 AGTTATAT
204 AGTTATT
205 ATAAATA
206 ATAAATAA
207 ATAACA
208 ATAACC
209 ATAACG
210 ATAAGC
211 ATAATA
212 ATAATAC
213 ATAATCA
226 ATATAC
227 ATATACA
228 ATATAG
229 ATATAT
230 ATATATT
231 ATATCA
232 ATATCG
233 ATATGA
234 ATATTAA
235 ATATTAT
236 ATCACG
237 ATCCCG
238 ATCCTA

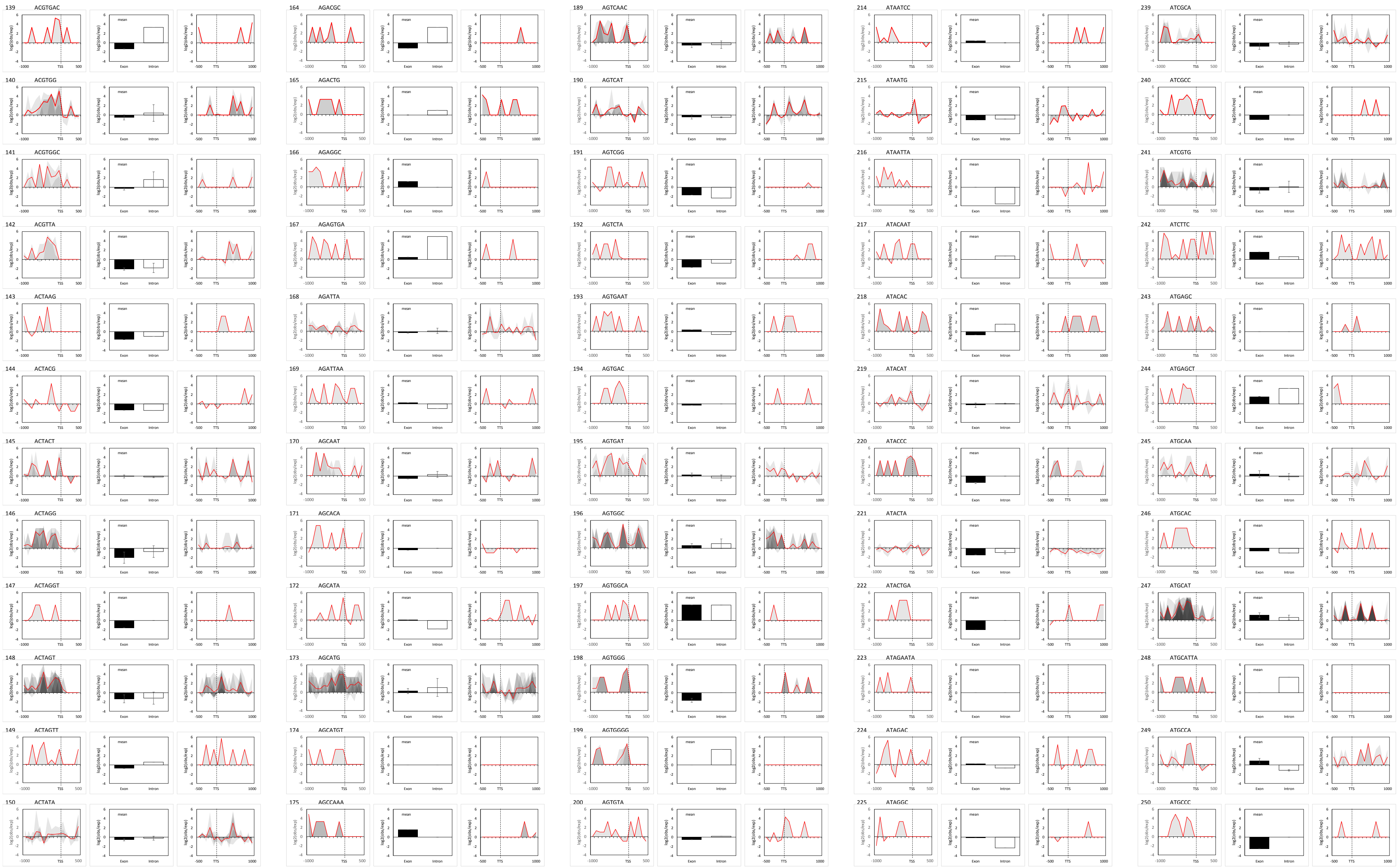
139 ACGTGAC
140 ACGTGG
141 ACGTGGC
142 ACGTTA
143 ACTAAG
144 ACTACG
145 ACTACT
146 ACTAGG
147 ACTAGGT
148 ACTAGT
149 ACTAGTT
150 ACTATA
164 AGACGC
165 AGACTG
166 AGAGGC
167 AGAGTGA
168 AGATTA
169 AGATTAA
170 AGCAAT
171 AGCACA
172 AGCATA
173 AGCATG
174 AGCATGT
175 AGCCAAA
189 AGTCAAC
190 AGTCAT
191 AGTCGG
192 AGTCTA
193 AGTGAAT
194 AGTGAC
195 AGTGAT
196 AGTGGC
197 AGTGGCA
198 AGTGGG
199 AGTGGGG
200 AGTGTA
214 ATAATCC
215 ATAATG
216 ATAATTA
217 ATACAAT
218 ATACAC
219 ATACAT
220 ATACCC
221 ATACTA
222 ATACTGA
223 ATAGAATA
224 ATAGAC
225 ATAGGC
239 ATCGCA
240 ATCGCC
241 ATCGTG
242 ATCTTC
243 ATGAGC
244 ATGAGCT
245 ATGCAA
246 ATGCAC
247 ATGCAT
248 ATGCATTA
249 ATGCCA
250 ATGCCC

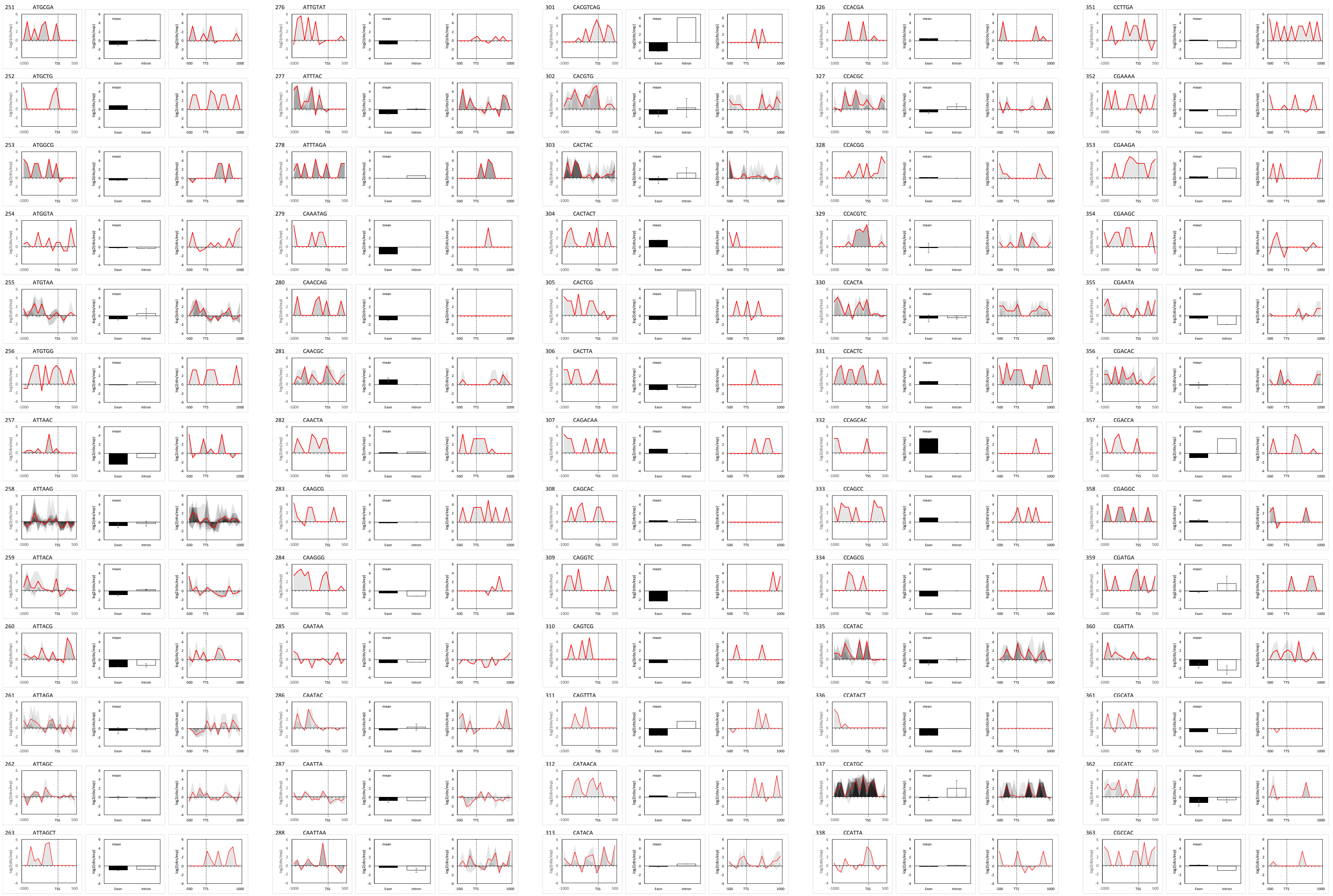
251 ATGCGA
252 ATGCTG
253 ATGGCG
254 ATGGTA
255 ATGTAA
256 ATGTGG
257 ATTAAC
258 ATTAAG
259 ATTACA
260 ATTACG
261 ATTAGA
262 ATTAGC
263 ATTAGCT
276 ATTGTAT
277 ATTTAC
278 ATTTAGA
279 CAAATAG
280 CAACCAG
281 CAACGC
282 CAACTA
283 CAAGCG
284 CAAGGG
285 CAATAA
286 CAATAC
287 CAATTA
288 CAATTAA
301 CACGTCAG
302 CACGTG
303 CACTAC
304 CACTACT
305 CACTCG
306 CACTTA
307 CAGACAA
308 CAGCAC
309 CAGGTC
310 CAGTCG
311 CAGTTTA
312 CATAACA
313 CATACA
326 CCACGA
327 CCACGC
328 CCACGG
329 CCACGTC
330 CCACTA
331 CCACTC
332 CCAGCAC
333 CCAGCC
334 CCAGCG
335 CCATAC
336 CCATACT
337 CCATGC
338 CCATTA
351 CCTTGA
352 CGAAAA
353 CGAAGA
354 CGAAGC
355 CGAATA
356 CGACAC
357 CGACCA
358 CGAGGC
359 CGATGA
360 CGATTA
361 CGCATA
362 CGCATC
363 CGCCAC
log2(obs/exp)
mean
Exon
Intron
-1000
TSS
500
-500
TTS
1000

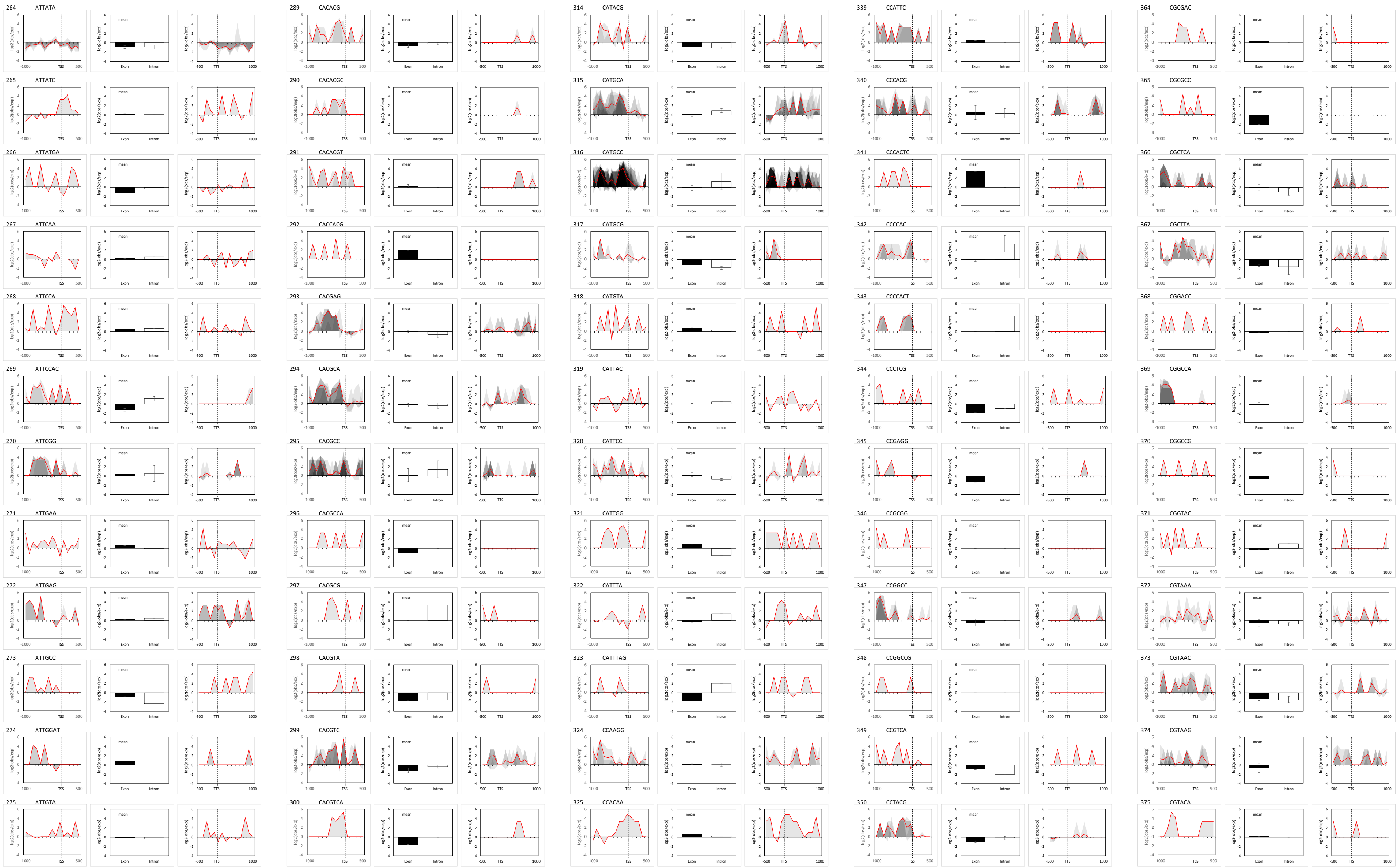

264 ATTATA
265 ATTATC
266 ATTATGA
267 ATTCAA
268 ATTCCA
269 ATTCCAC
270 ATTCGG
271 ATTGAA
272 ATTGAG
273 ATTGCC
274 ATTGGAT
275 ATTGTA
289 CACACG
290 CACACGC
291 CACACGT
292 CACCACG
293 CACGAG
294 CACGCA
295 CACGCC
296 CACGCCA
297 CACGCG
298 CACGTA
299 CACGTC
300 CACGTCA
314 CATACG
315 CATGCA
316 CATGCC
317 CATGCG
318 CATGTA
319 CATTAC
320 CATTCC
321 CATTGG
322 CATTTA
323 CATTTAG
324 CCAAGG
325 CCACAA
339 CCATTC
340 CCCACG
341 CCCACTC
342 CCCCAC
343 CCCCACT
344 CCCTCG
345 CCGAGG
346 CCGCGG
347 CCGGCC
348 CCGGCCG
349 CCGTCA
350 CCTACG
364 CGCGAC
365 CGCGCC
366 CGCTCA
367 CGCTTA
368 CGGACC
369 CGGCCA
370 CGGCCG
371 CGGTAC
372 CGTAAA
373 CGTAAC
374 CGTAAG
375 CGTACA

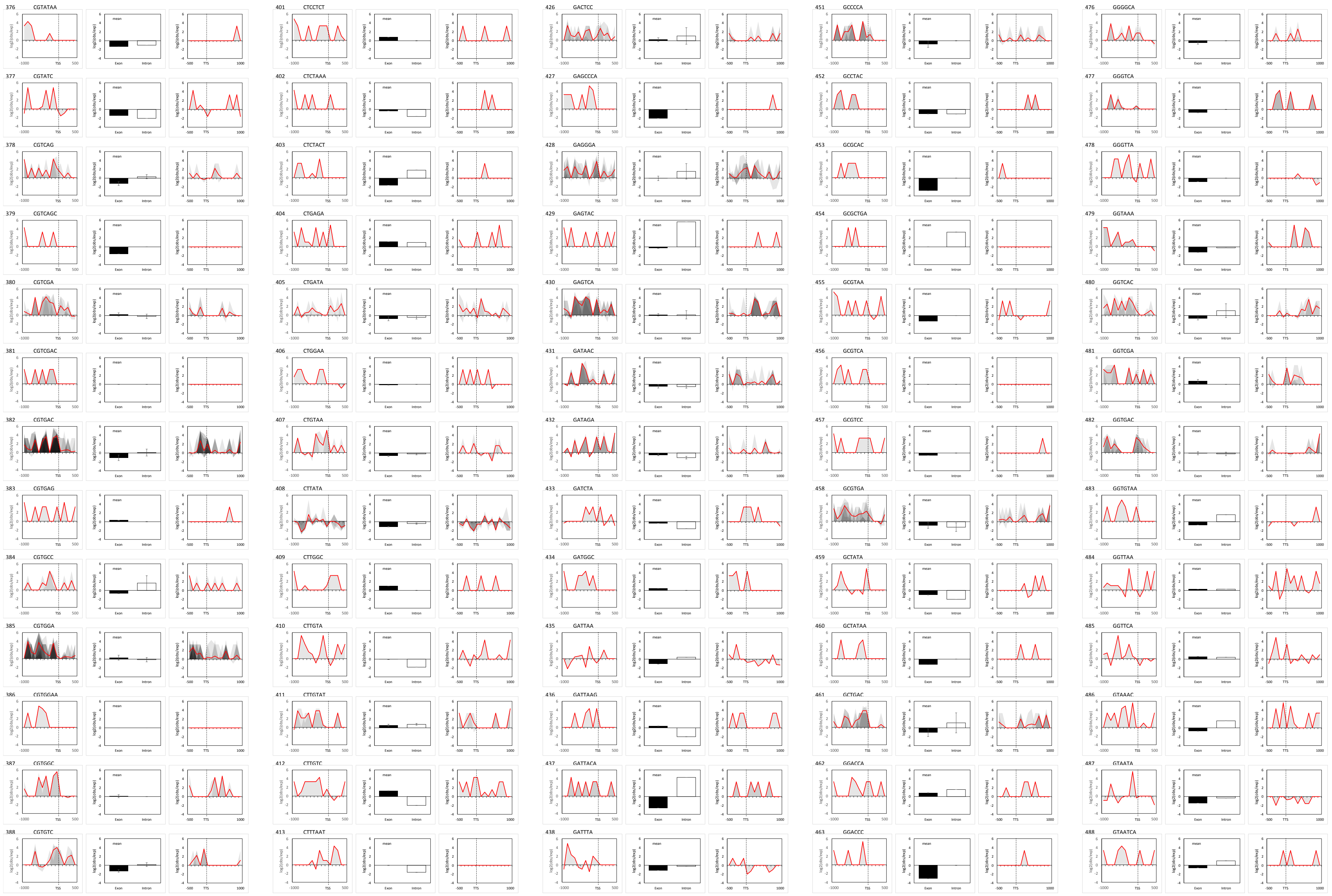

376 CGTATAA
377 CGTATC
378 CGTCAG
379 CGTCAGC
380 CGTCGA
381 CGTCGAC
382 CGTGAC
383 CGTGAG
384 CGTGCC
385 CGTGGA
386 CGTGGAA
387 CGTGGC
388 CGTGTC
401 CTCCTCT
402 CTCTAAA
403 CTCTACT
404 CTGAGA
405 CTGATA
406 CTGGAA
407 CTGTAA
408 CTTATA
409 CTTGGC
410 CTTGTA
411 CTTGTAT
412 CTTGTC
413 CTTTAAT
426 GACTCC
427 GAGCCCA
428 GAGGGA
429 GAGTAC
430 GAGTCA
431 GATAAC
432 GATAGA
433 GATCTA
434 GATGGC
435 GATTAA
436 GATTAAG
437 GATTACA
438 GATTTA
451 GCCCCA
452 GCCTAC
453 GCGCAC
454 GCGCTGA
455 GCGTAA
456 GCGTCA
457 GCGTCC
458 GCGTGA
459 GCTATA
460 GCTATAA
461 GCTGAC
462 GGACCA
463 GGACCC
476 GGGGCA
477 GGGTCA
478 GGGTTA
479 GGTAAA
480 GGTCAC
481 GGTCGA
482 GGTGAC
483 GGTGTAA
484 GGTTAA
485 GGTTCA
486 GTAAAC
487 GTAATA
488 GTAATCA
log2(obs/exp)
mean
Exon
Intron
TSS
TTS

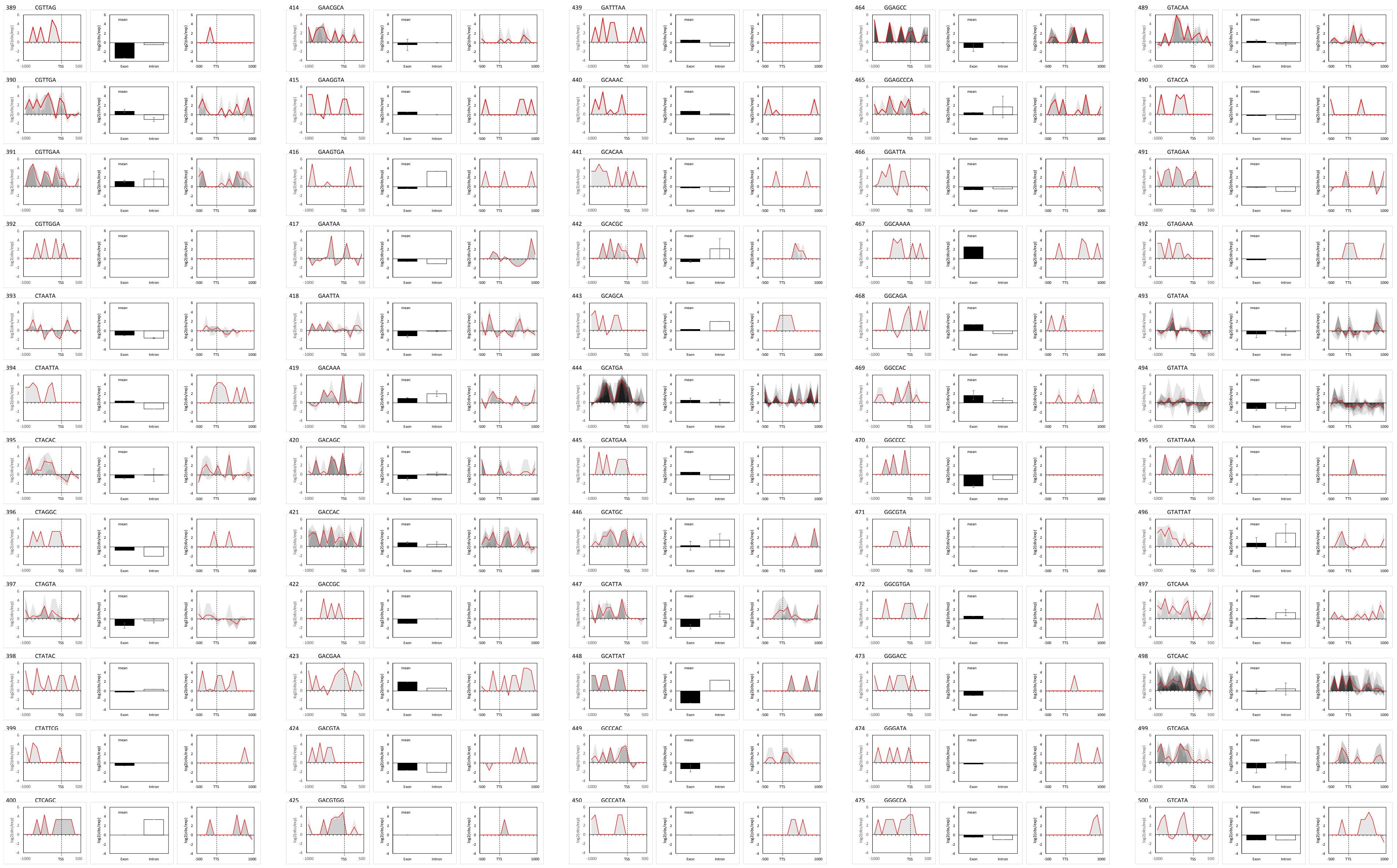

389 CGTTAG
390 CGTTGA
391 CGTTGAA
392 CGTTGGA
393 CTAATA
394 CTAATTA
395 CTACAC
396 CTAGGC
397 CTAGTA
398 CTATAC
399 CTATTCG
400 CTCAGC
414 GAACGCA
415 GAAGGTA
416 GAAGTGA
417 GAATAA
418 GAATTA
419 GACAAA
420 GACAGC
421 GACCAC
422 GACCGC
423 GACGAA
424 GACGTA
425 GACGTGG
439 GATTTAA
440 GCAAAC
441 GCACAA
442 GCACGC
443 GCAGCA
444 GCATGA
445 GCATGAA
446 GCATGC
447 GCATTA
448 GCATTAT
449 GCCCAC
450 GCCCATA
464 GGAGCC
465 GGAGCCCA
466 GGATTA
467 GGCAAAA
468 GGCAGA
469 GGCCAC
470 GGCCCC
471 GGCGTA
472 GGCGTGA
473 GGGACC
474 GGGATA
475 GGGCCA
489 GTACAA
490 GTACCA
491 GTAGAA
492 GTAGAAA
493 GTATAA
494 GTATTA
495 GTATTAAA
496 GTATTAT
497 GTCAAA
498 GTCAAC
499 GTCAGA
500 GTCATA

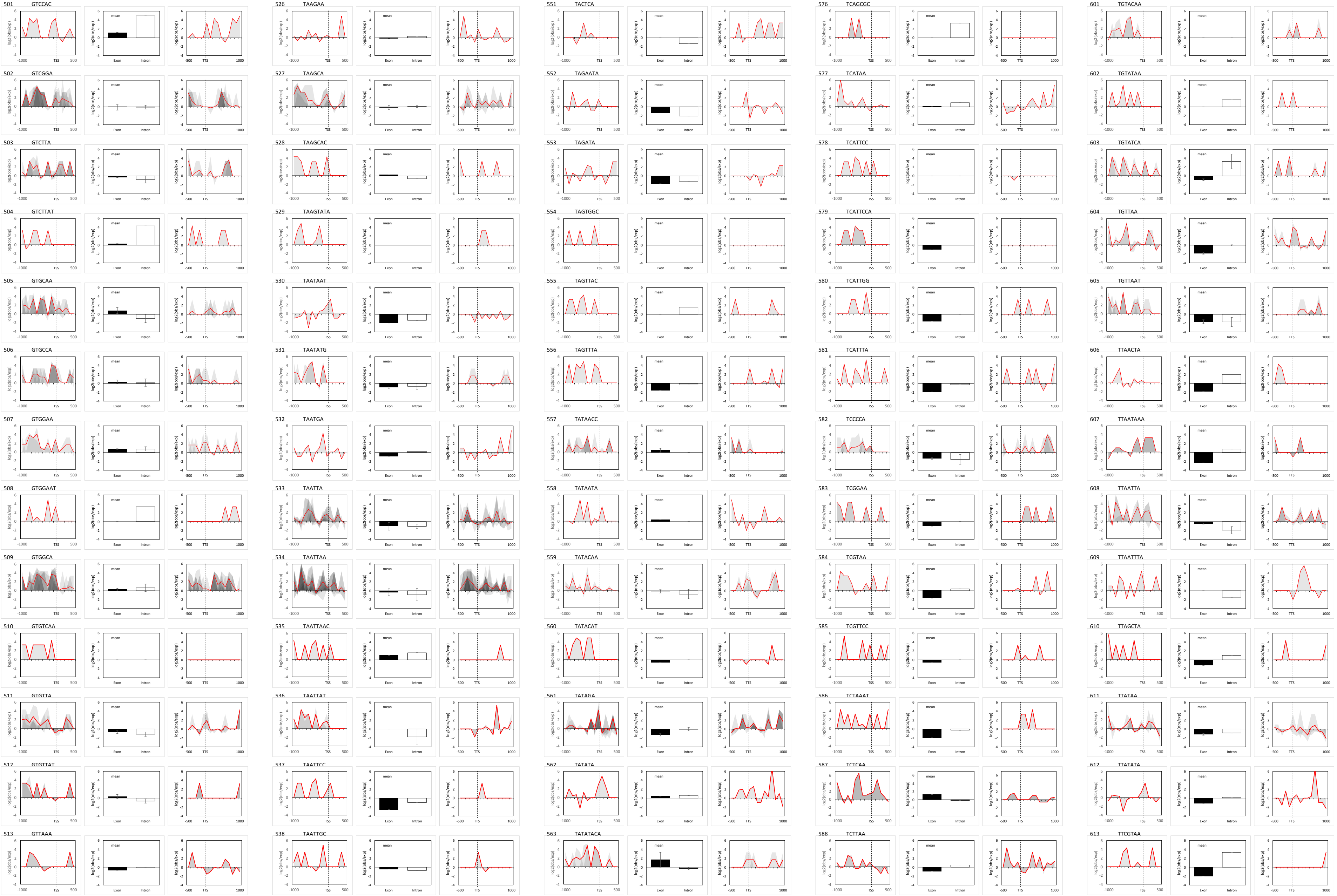

501 GTCCAC
502 GTCGGA
503 GTCTTA
504 GTCTTAT
505 GTGCAA
506 GTGCCA
507 GTGGAA
508 GTGGAAT
509 GTGGCA
510 GTGTCAA
511 GTGTTA
512 GTGTTAT
513 GTTAAA
526 TAAGAA
527 TAAGCA
528 TAAGCAC
529 TAAGTATA
530 TAATAAT
531 TAATATG
532 TAATGA
533 TAATTA
534 TAATTAA
535 TAATTAAC
536 TAATTAT
537 TAATTCC
538 TAATTGC
551 TACTCA
552 TAGAATA
553 TAGATA
554 TAGTGGC
555 TAGTTAC
556 TAGTTTA
557 TATAACC
558 TATAATA
559 TATACAA
560 TATACAT
561 TATAGA
562 TATATA
563 TATATACA
576 TCAGCGC
577 TCATAA
578 TCATTCC
579 TCATTCCA
580 TCATTGG
581 TCATTTA
582 TCCCCA
583 TCGGAA
584 TCGTAA
585 TCGTTCC
586 TCTAAAT
587 TCTCAA
588 TCTTAA
601 TGTACAA
602 TGTATAA
603 TGTATCA
604 TGTTAA
605 TGTTAAT
606 TTAACTA
607 TTAATAAA
608 TTAATTA
609 TTAATTTA
610 TTAGCTA
611 TTATAA
612 TTATATA
613 TTCGTAA

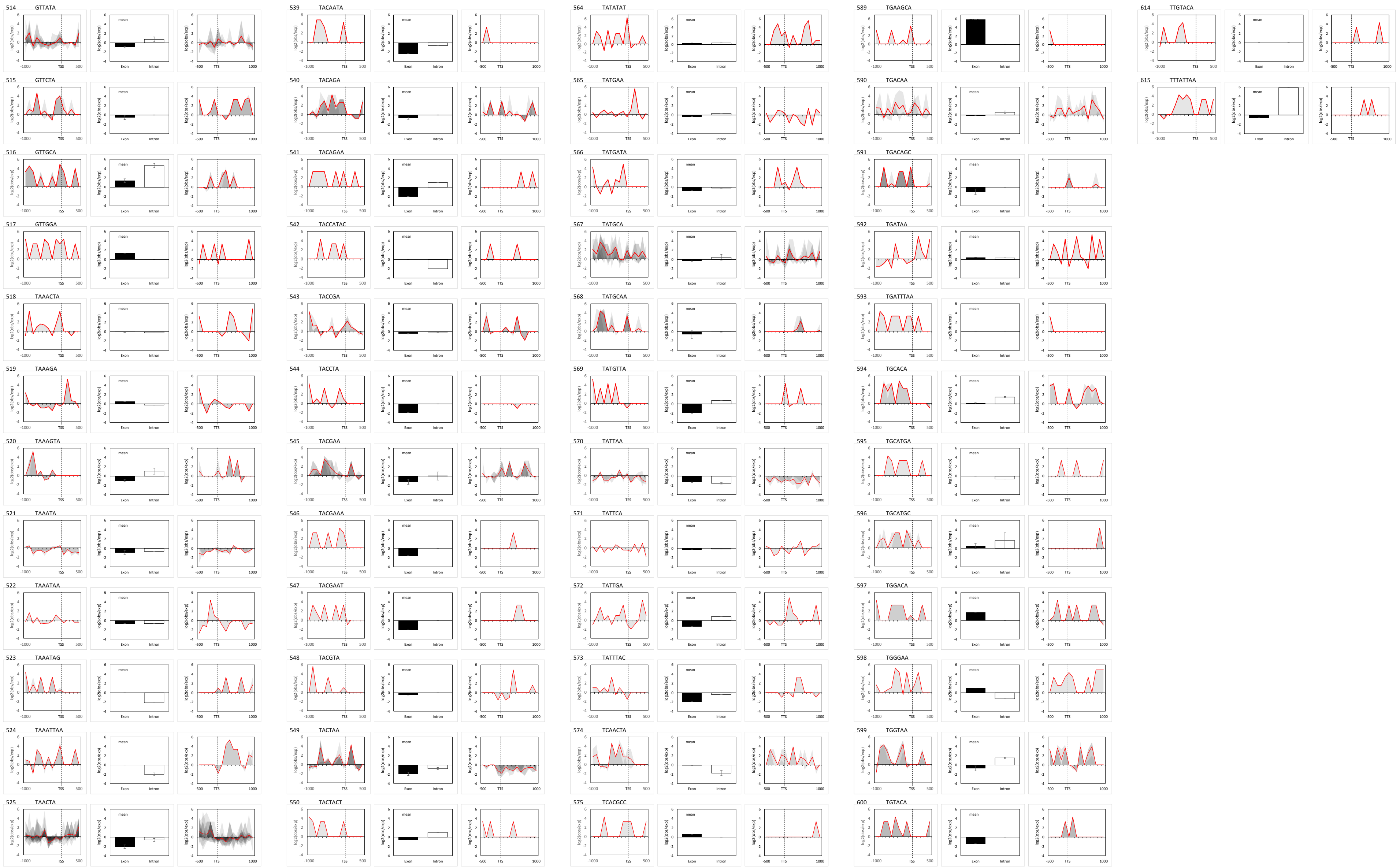

514 GTTATA
515 GTTCTA
516 GTTGCA
517 GTTGGA
518 TAAACTA
519 TAAAGA
520 TAAAGTA
521 TAAATA
522 TAAATAA
523 TAAATAG
524 TAAATTAA
525 TAACTA
539 TACAATA
540 TACAGA
541 TACAGAA
542 TACCATAC
543 TACCGA
544 TACCTA
545 TACGAA
546 TACGAAA
547 TACGAAT
548 TACGTA
549 TACTAA
550 TACTACT
564 TATATAT
565 TATGAA
566 TATGATA
567 TATGCA
568 TATGCAA
569 TATGTTA
570 TATTAA
571 TATTCA
572 TATTGA
573 TATTTAC
574 TCAACTA
575 TCACGCC
589 TGAAGCA
590 TGACAA
591 TGACAGC
592 TGATAA
593 TGATTTAA
594 TGCACA
595 TGCATGA
596 TGCATGC
597 TGGACA
598 TGGGAA
599 TGGTAA
600 TGTACA
614 TTGTACA
615 TTTATTAA
log2(obs/exp)
mean
Exon
Intron
TSS
TTS
