## Supplemental_Figures_S1-S7_Supplemental_Table_S1_Supplemental_literature for "Putative *cis*-regulatory elements predict iron deficiency responses in Arabidopsis roots"

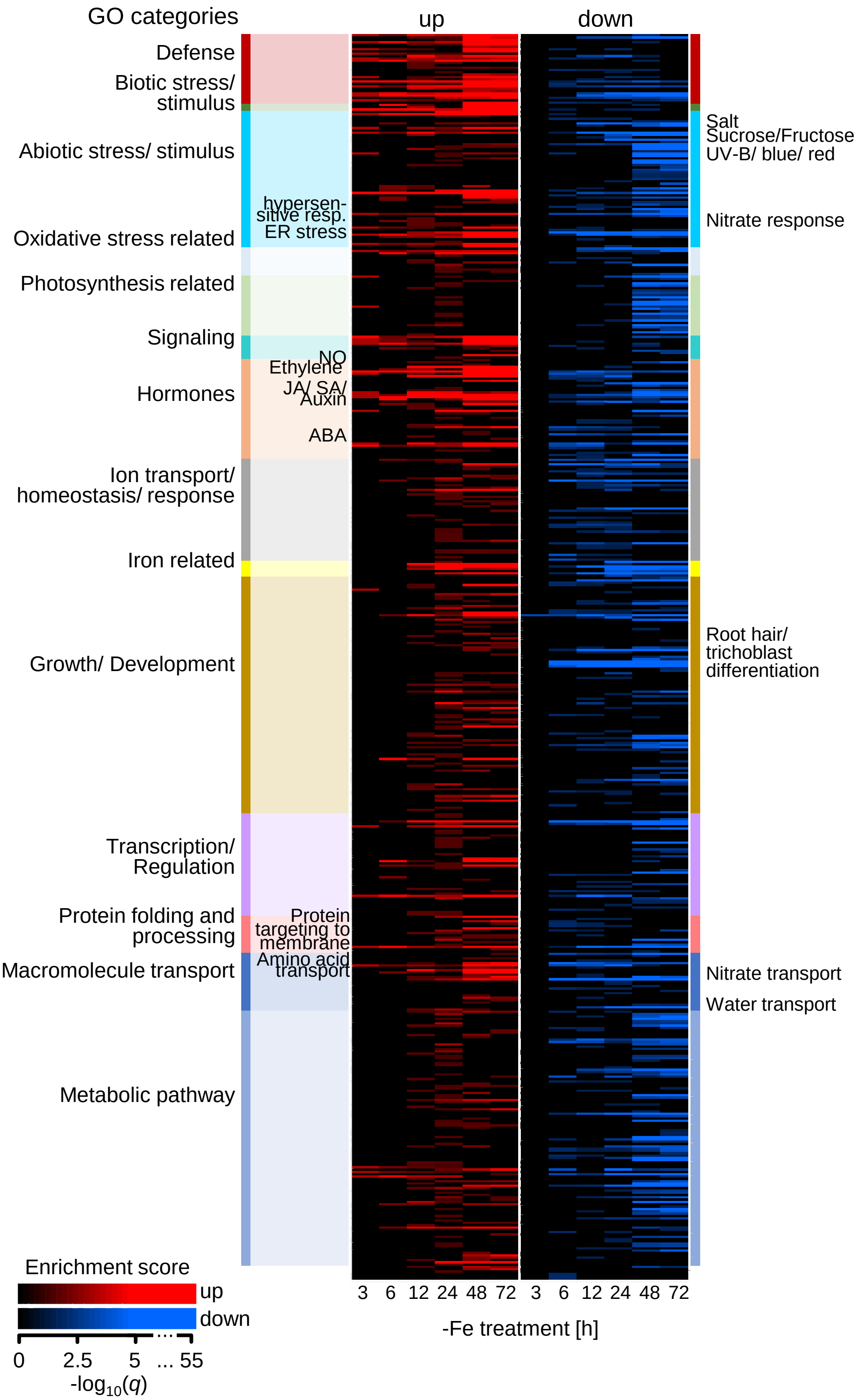


**Figure S1. Complete GO enrichment analysis of -Fe-responsive genes.**
Heatmap of enrichment (FET, $q<0.05$) of GO terms in genes that were significantly up- (red) or down-regulated (blue) ($q<0.05$) at ≥1 of 6 time points in -Fe-treated roots of 6 d-old seedlings (Dinneny et al., 2008). Differential regulation was defined as $\log_2$ fold-change (log2FC) >1 or <-1 (treatment vs. control). Selected highly enriched GO terms are indicated. Maximum color intensity threshold was set to $-\log_{10}(q)=5$ for visualization purposes. (Original *p*- and *q*-values: **Supplemental Table S7**).

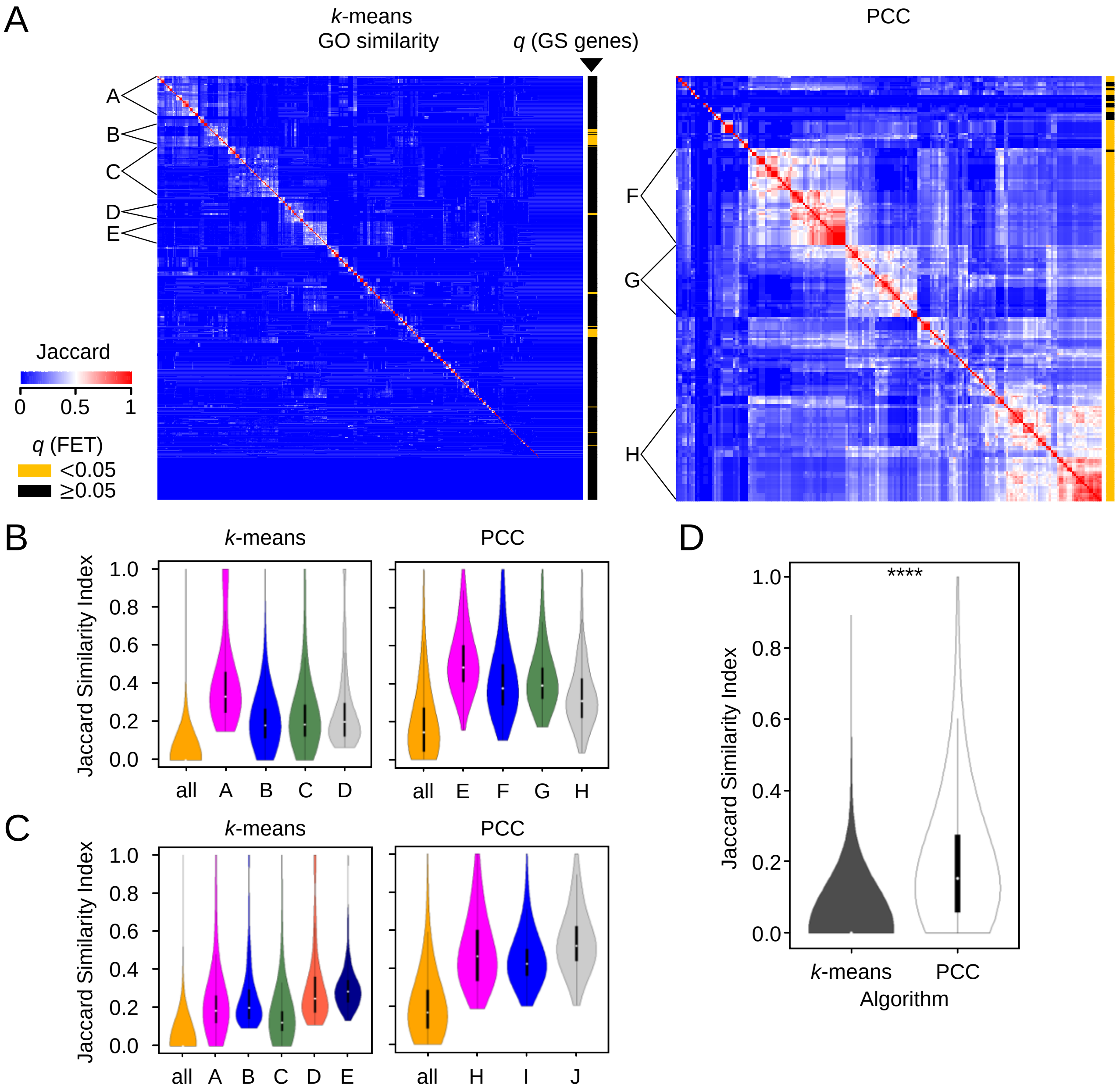


**Figure S2. GO terms and -Fe GS gene enrichments of the defined co-expression clusters containing up-/down-regulated genes, and mean similarity within designated superclusters of up-regulated and up-/down-regulated genes.**

**A:** Heatmap of GO similarity, measured using the Jaccard Index, between co-expression clusters from *k*-means clustering (**left**, n=535) and GS gene correlation (PCC; **right**, n=201), generated from expression data containing up- and down-regulated genes. Clusters were grouped by hierarchical clustering and superclusters (A-H) were defined as groups of >20 clusters that have a within-mean Jaccard Index significantly higher than the mean Jaccard Index of all clusters. Enriched GO terms shared by ≥75% (*k*-means) and ≥90% (PCC) of the clusters in each supercluster are shown (**left**). Co-expression clusters enriched for -Fe GS genes are designated (yellow, **right**). ***k*-means superclusters**: A, Transport nitrate, Response to nitrate. B, Cellular response to Fe/ethylene stimulus/NO. C, Response to red light. D, Regulation of $H_2O_2$ metabolic process, Response to hypoxia, SAR-SA mediated pathway. E, Defense response to fungus, Response to chitin. **PCC superclusters**: F, Cellular response to Fe/ethylene stimulus/NO. G, Transport Pb/Cd/Mn, Homeostasis metal ion/Fe, Pos. regulation of ROS species metabolic process. H, Transport Fe/Zn/nitrate, Cellular response to Fe/ethylene stimulus/NO, Response to Zn/nitrate, Detoxification Zn/Co. All enrichment analyses: FET, $q<0.05$.

**B:** Violin plots showing Jaccard Index distributions of all *k*-means and all PCC clusters (“all”), and of superclusters A-H, generated from expression data containing up-regulated genes (**Figure 2A**). Statistical analysis of each *k*-means or PCC supercluster compared to the respective “all”: Mann-Whitney U ($p<2.2e-16$).

**C:** As in S2B, for superclusters shown in S2A. Mann-Whitney U of each supercluster compared to “all” ($p<2.2e-16$).

**D:** Gene content redundancy of -Fe co-expression clusters, corresponding to **Figure 2A** and **S2A**. Gene content similarity within all *k*-means and all PCC clusters. Clusters were compared in pair-wise manner with the Jaccard Index (1=identical gene content). Statistical analysis comparing *k*-means and PCC mean Jaccard Index: Mann-Whitney U (**** $p<2.2e-16$).

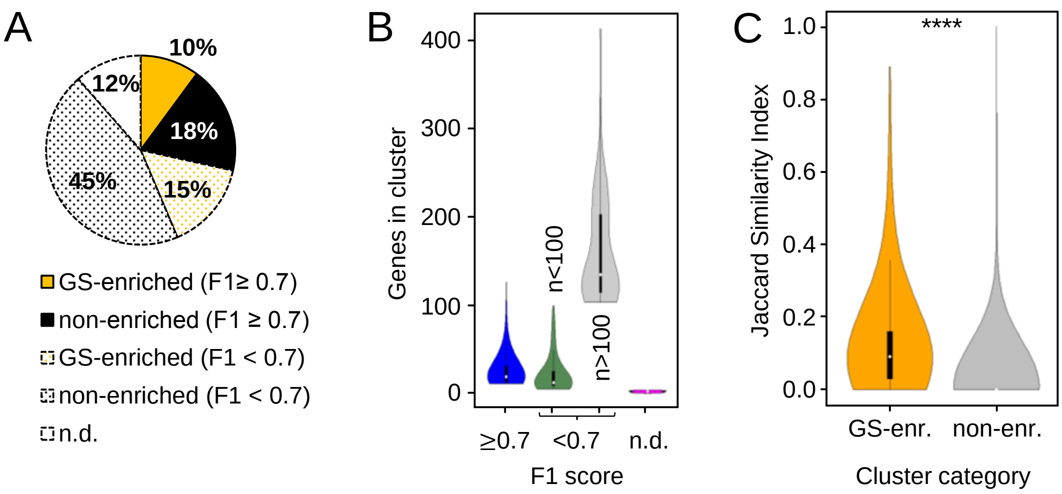


**Figure S3. Co-expression cluster RF model performance of cluster category (GS-enriched and non-enriched) and cluster size.**

**A:** Proportions of co-expression clusters with good (F1≥0.7; filled) and poor (dotted) RF performance organized by cluster category (GS-enriched; yellow, and non-enriched; black), and clusters which did not pass RF (n.d.; white).

**B:** Violin plots of cluster size distributions of all well-performing clusters (n=558), poor-performing clusters with <100 genes (n=1111), poor-performing clusters with >100 genes (n=66) and clusters which did not pass RF (n.d.; n=224).

**C:** Gene content similarity of well-performing GS-enriched (yellow; n=159) and non-enriched (gray; n=358) co-expression clusters (redundant correlation-based clusters not included; see **Methods**). Clusters within each category were compared in pair-wise manner with the Jaccard Index (1=identical gene content). Statistical analysis comparing mean Jaccard Index of both categories: Mann-Whitney U (**** $p$<2.2e-16). Cluster details: **Supplemental Table S2**.

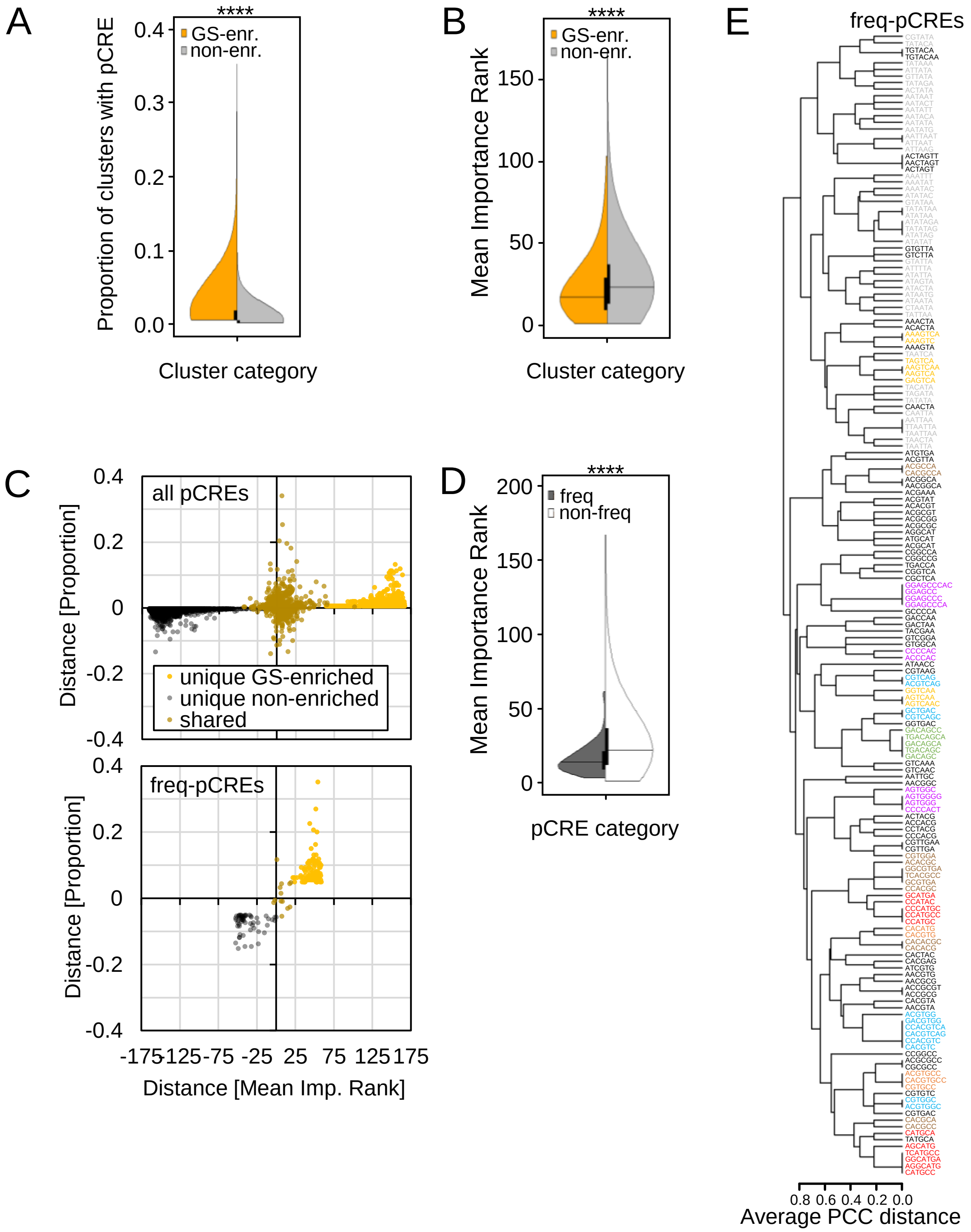


**Figure S4. Comparison of the pCRE abundance and importance in GS-enriched clusters vs. non-enriched clusters and hierarchical clustering of freq-pCRE sequences.**

**A:** Distribution of proportions of GS-enriched (yellow) and non-enriched (gray) clusters in which each pCRE (n=5,639) was identified. Statistical analysis: Mann-Whitney U (**** *p*<2.2e-16).

**B:** Distribution of the mean importance rank (1=most important) of each pCRE across GS-enriched and non-enriched clusters. Statistical analysis: Mann-Whitney U (**** *p*<2.2e-16).

**C:** Distance plots showing mean importance rank and proportion in GS-enriched vs. non-enriched clusters of all 5,639 pCREs (**top**) and 173 freq-pCREs (**bottom**). Distances were calculated as follows: prop[GS] – prop[non] and rank[non]–rank[GS] (positive value: pCRE has a higher proportion and/or is more important in GS-enriched clusters). Note that most shared pCREs in (**top**) become unique freq-pCREs in (**bottom**). In contrast to S4A, B, and Figure 4A, B, which depict shared pCREs twice, each pCRE/freq-pCRE is represented once. Therefore, a preference for GS-enriched or non-enriched clusters can be derived for shared pCREs/freq-pCREs.

**D:** Mean importance rank distributions of freq-pCREs (n=173) and non-freq-pCREs (n=5,466). Statistical analysis: Mann-Whitney U (**** *p*<1.924e-14).

**E:** PCC distances of freq-pCRE PWMs (hierarchical clustering). Due to group-wise averaging of PCC distances during hierarchical clustering, the algorithm produced skewed PCC distances of some pCRE pairs. Therefore, all pair-wise PCC distances were evaluated individually and corrected, if necessary, by hand in the freq-pCRE network (**Figure 4C**). freq-pCRE colors correspond to freq-pCRE network groups in **Figure 4C**, red: group1, orange: group2, pink: group3, gray: group4, yellow: group5, green: group6, brown: group7, blue: group8.

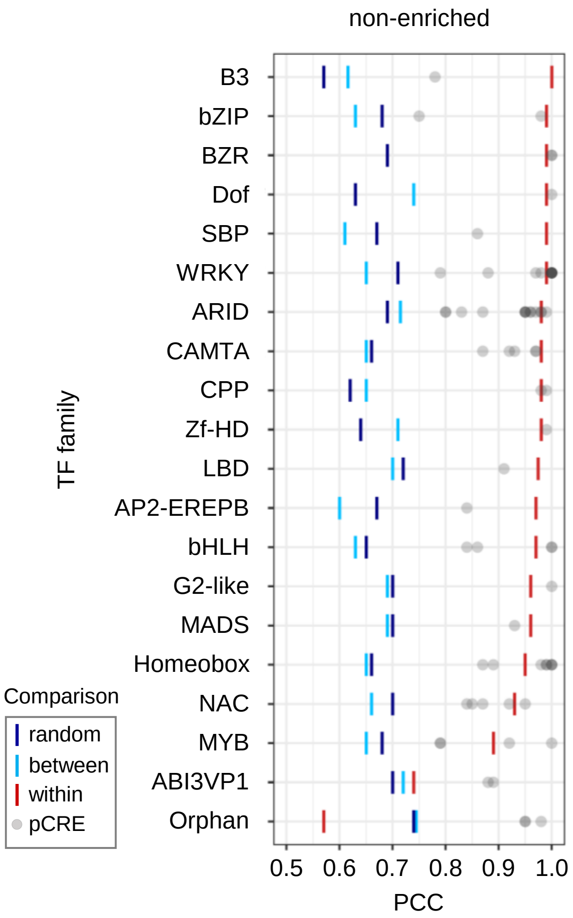


**Figure S5. Significance of sequence similarity for freq-pCRE from non-enriched clusters and the best matching known TFBM.**
Bars represent 95th percentile (PCC) significance thresholds for within TF family (red, pCRE sequence is more similar to a specific TFBM than other TFBMs from the same family), between TF families (light blue, pCRE sequence is more similar to a TFBM in a TF family than TFBMs from other TF families), or random (dark blue, pCRE sequence is more similar to a TFBM from a family than random 6-mers).

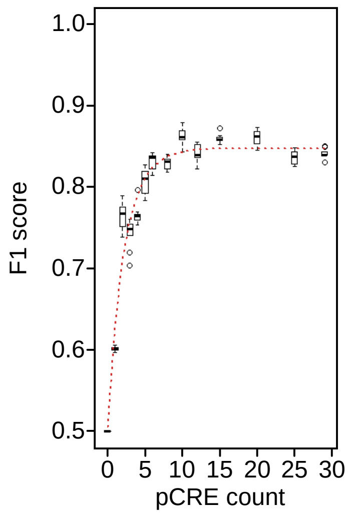


**Figure S6. Example of a non-linear regression curve to determine the minimum set of pCREs for a co-expression cluster.**

The minimum set was defined as the minimum number of pCREs (min-pCREs) needed for the model to perform at its maximum (F1 remains unchanged). Low-ranked pCREs or pCREs that were redundant (one is subset of the other) to a higher ranked pCRE were removed in step-wise manner and the predictive power of the remaining pCREs was tested in 10 RF runs. In this example, the top 10 pCREs were kept. Cluster ID: 513 (**Supplemental Table S2**).

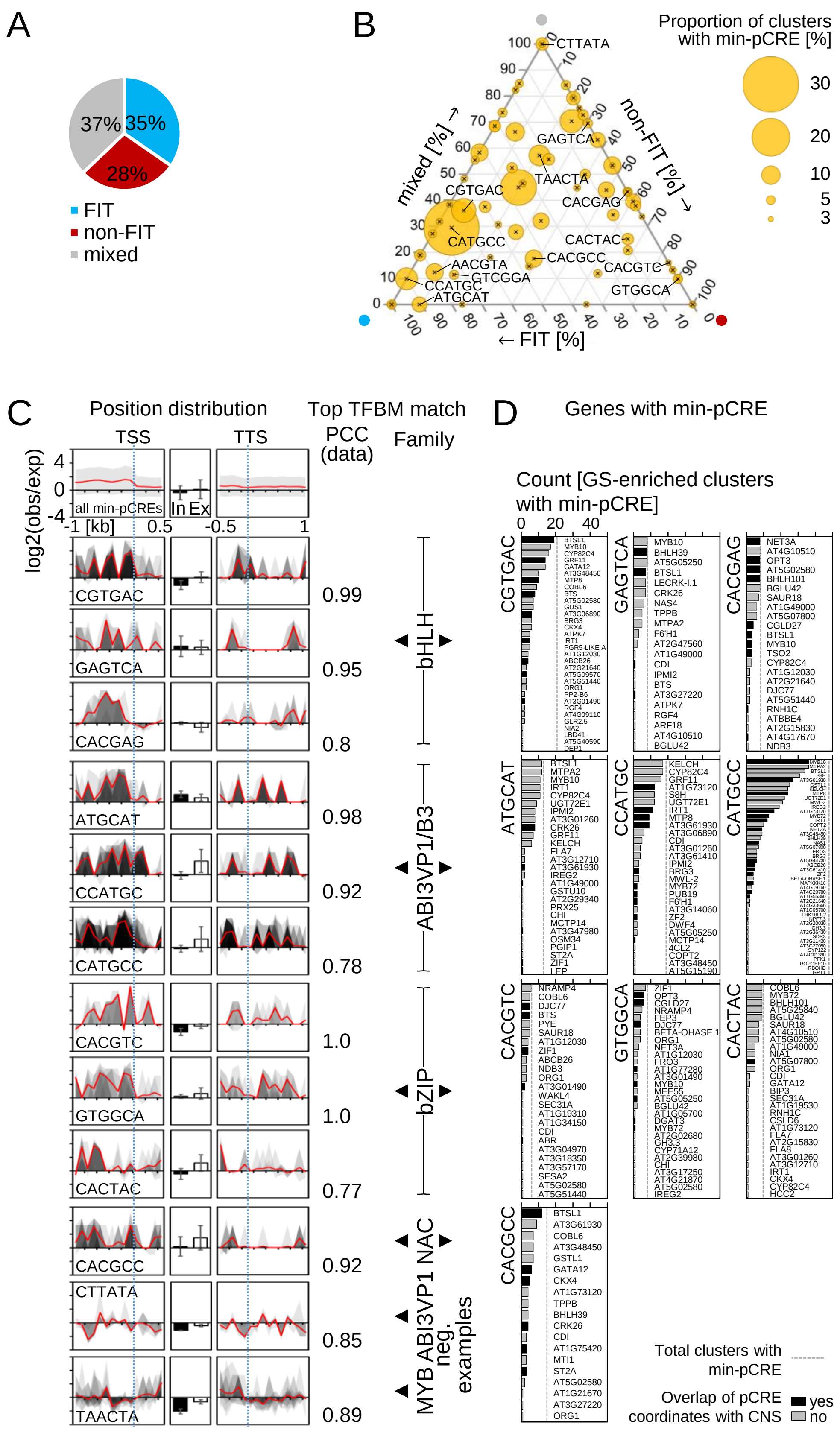


**Figure S7.** High-resolution image of **Figure 6**.

**Table S1**. Robust -Fe-responsive gold standard (GS) genes. -Fe GS genes are reliably up-regulated under -Fe in Arabidopsis roots or seedlings across several independent studies, some of which are transcriptomic and others are targeted. A subset of -Fe GS genes is FIT-dependent whereas other GS genes are FIT-independent.

| AGI | Description | References |
|---|---|---|
| FIT-dependent | | |
| AT1G34760 | ROOT HAIR SPECIFIC 5 (RHS5), GENERAL REGULATORY FACTOR 11 (GRF11), 14-3-3 protein | 1-6, 8, 9, 10 |
| AT1G56160 | MYB72, transcription factor | 1-9, 11, 12 |
| AT3G12820 | MYB10, transcription factor | 1-6, 8, 9, 11 |
| AT1G73120 | F-box/RNI superfamily protein | 1, 2, 4, 5, 7-9 |
| AT2G20030 | RING/U-box superfamily protein | 1, 3, 4, 6, 7, 9 |
| AT3G07720 | Galactose oxidase/kelch repeat superfamily protein | 1-9 |
| AT3G12900 | SCOPOLETIN 8-HYDROXYLASE (S8H) | 1-9, 13-15 |
| AT3G50740 | UDP-GLUCOSYL TRANSFERASE 72E1 (UGT72E1) | 1, 2, 5-9 |
| AT4G31940 | CYTOCHROME P450, FAMILY 82, SUBFAMILY C, POLYPEPTIDE 4 (CYP82C4) | 1, 2, 5-8, 16 |
| AT4G30120 | HEAVY METAL ATPASE (HMA3) | 1, 2, 5, 6, 8 |
| AT4G19690 | IRON-REGULATED TRANSPORTER 1 (IRT1) | 1-9, 17, 18 |
| AT5G03570 | IRON-REGULATED PROTEIN 2 (IREG2) | 1-3, 5-9, 19 |
| AT3G58810 | METAL TOLERANCE PROTEIN A2 (MTPA2), putative zinc transporter | 1-9, 20 |
| AT4G33020 | ZIP9, metal ion transporter | 1, 2, 5, 6, 8 |
| AT5G38820 | Putative amino acid transporter | 1-5, 8, 9 |
| FIT-independent | | |
| AT3G56980 | OBP3-RESPONSIVE GENE 3 (ORG3), bHLH039, transcription factor | 2, 3, 5-8, 21-23 |
| AT5G04150 | bHLH101, transcription factor | 2, 3, 5, 6, 8, 21 |
| AT3G47640 | POPEYE (PYE), bHLH transcription factor | 2, 3, 5-8 |
| AT3G18290 | EMBRYO DEFECTIVE 2454 (EMB2454), BRUTUS (BTS), putative E3 ligase | 2, 3, 5-7, 24 |
| AT1G74770 | BRUTUS LIKE 1 (BTSL1) | 2, 3, 6-8, 25 |
| AT5G53450 | OBP3-RESPONSIVE GENE 1 (ORG1) | 2, 3, 5-8 |
| AT1G56430 | NICOTIANAMINE SYNTHASE 4 (NAS4) | 2, 5-7, 26 |
| AT1G23020 | FERRIC REDUCTION OXIDASE 3 (FRO3) | 2, 3, 5-8, 27 |
| AT4G16370 | OLIGOPEPTIDE TRANSPORTER 3 (OPT3) | 2, 3, 5-8, 28, 29 |
| AT5G67330 | NATURAL RESISTANCE ASSOCIATED MACROPHAGE PROTEIN 4 (NRAMP4), metal ion transporter | 2, 3, 5-7, 30 |
| AT5G13740 | ZINC INDUCED FACILITATOR 1 (ZIF1), small organic molecule transporter | 2, 3, 5-8, 31 |
| AT1G47400 | FE-UPTAKE-INDUCING PEPTIDE3 (FEP3), IRONMAN 1 (IMA1) | 2, 3, 5-8, 32, 33 |
| AT5G05250 | Hypothetical protein | 2, 3, 5-8 |

[1]Colangelo and Guerinot, 2004, [2]Dinneny et al., 2008 (time course, up in at least 1 of 6 time points), [3]Buckhout et al., 2009, [4]García et al., 2010, [5]Long et al., 2010, [6]Yang et al., 2010, [7]Ivanov et al., 2012, [8]Sivitz et al., 2012, [9]Mai et al., 2016, [10]Yang et al., 2013, [11]Palmer et al., 2013, [12]Zamioudis et al., 2015, [13]Siwinska et al., 2018, [14]Tsai et al., 2018, [15]Rajniak et al., 2018, [16]Murgia et al., 2011, [17]Vert et al., 2002, [18]Jakoby et al., 2004, [19]Schaaf et al., 2006, [20]Arrivault et al., 2006, [21]Wang et al., 2007, [22]Yuan et al., 2008, [23]Naranjo-Arcos et al., 2017, [24]Selote et al., 2015, [25]Hindt et al., 2017, [26]Klatte et al., 2009, [27]Mukherjee et al., 2006, [28]Mendoza-Cózatl et al., 2014, [29]Zhai et al., 2014, [30]Lanquar et al., 2005, [31]Haydon et al., 2012, [32]Hirayama et al., 2018, [33]Grillet et al., 2018. Gene annotation based on the The Arabidopsis Information Resource (TAIR) 10.0 genome release, www.arabidopsis.org (accessed 03/2019).
